# Expansion of the Catalytic Repertoire of Alcohol Dehydrogenases in Plant Metabolism

**DOI:** 10.1101/2022.07.24.501124

**Authors:** Chloe Langley, Evangelos Tatsis, Benke Hong, Yoko Nakamura, Christian Paetz, Clare E. M. Stevenson, Jerome Basquin, David M. Lawson, Lorenzo Caputi, Sarah E. O’Connor

**Author notes:** Supporting information for this article is given via a link at the end of the document.

## Abstract

Medium-chain alcohol dehydrogenases (ADHs) comprise a highly conserved enzyme family that catalyse the reversible reduction of aldehydes. However, recent discoveries in plant natural product biosynthesis suggest that the catalytic repertoire of ADHs has been expanded. Here we report the crystal structure of dihydroprecondylocarpine acetate synthase (DPAS), an ADH that catalyses the non-canonical 1,4 reduction of an α,β-unsaturated iminium moiety. Comparison with structures of plant-derived ADHs that catalyse 1,2-aldehyde and 1,2-iminium reductions suggest how the canonical ADH active site can be modified to carry out atypical carbonyl reductions, providing insight into how chemical reactions are diversified in plant metabolism.

---

Alcohol dehydrogenases (ADHs EC 1.1.1.1) are NAD(P)H-dependent medium-chain oxidoreductases found in all kingdoms of life. These enzymes typically catalyse the reversible reduction of aldehydes or ketones to the corresponding alcohol (Figure 1A) ^[1–4]^. The structural motifs of ADHs are highly conserved in all known eukaryotic examples; most notably, a zinc ion involved in catalysis, a second zinc ion involved in maintaining protein structure, and the Rossmann peptide-fold involved in cofactor binding (Figures S1). ADHs have been shown to catalyse many complex biochemical transformations in plant natural product biosynthesis, suggesting that their catalytic repertoire has been expanded. For example, we recently reported the discovery of dihydroprecondylocarpine acetate synthase (DPAS), an ADH involved in vinblastine biosynthesis in the plant *Catharanthus roseus* ^[5]^ and in ibogaine biosynthesis in the phylogenetically related species *Tabernanthe iboga* ^[6]^. Since the product of DPAS is unstable and either immediately decomposes or rearranges in the presence of a downstream cyclase enzyme, the reaction remained unsubstantiated (Figure 1B). However, the cyclised products suggest that DPAS catalyses the 1,4-reduction of an α,β-unsaturated iminium which is an hitherto unprecedented reaction for an ADH. Here, we use isotopic labelling to definitively establish that DPAS catalyses this unusual 1,4-reduction. We report four crystal structures of apo- and substrate-bound DPAS from two phylogenetically related species. These structures reveal, surprisingly, the loss of the catalytic zinc ion from the DPAS active site, indicating that zinc is not strictly required for reduction by ADHs. We also report the structure of the ADH geissoschizine synthase (GS) that catalyses an atypical 1,2 iminium reduction. Comparison of the active site of DPAS and GS with other highly similar ADHs that catalyse either 1,2 aldehyde reduction or 1,2 reduction of an iminium moiety suggests that changes in the proton relay system are also implicated in modulating ADH reactivity. The mechanism and structure of DPAS highlights the catalytic versatility of ADHs. Overall, these findings demonstrate how the active site of ADHs have been modulated to expand the catalytic scope of this large class of enzymes.

**Figure 1.**
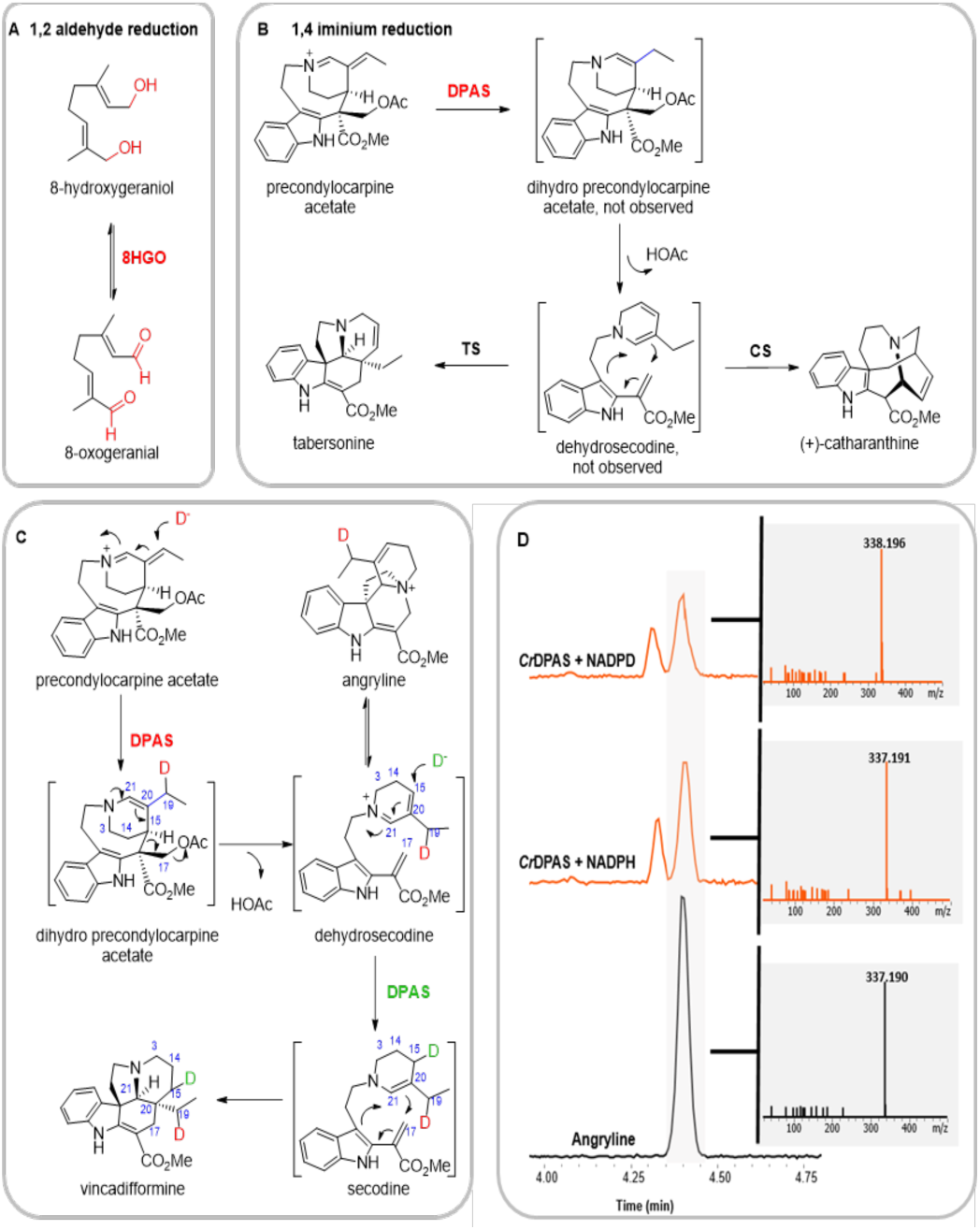
ADH catalysed reduction. **A.** Reversible oxidation/reduction reaction of 8-hydroxygeraniol/8-oxogeranial catalysed by canonical ADH 8HGO. **B.** Proposed 1,4-reduction of precondylocarpine acetate catalysed by DPAS and subsequent cyclisation reaction catalysed by various cyclase enzymes such as tabersonine synthase (TS) or catharanthine synthase (CS). **C.** Isotopic labelling of two 1,4-reductions catalysed by DPAS. **D.** TIC of precondylocarpine acetate reacted with *Cr*DPAS in the presence of NADPH or deuterated NADPD and MS/MS spectra showing +1 mass change indicating deuterium incorporation.

The recently discovered DPAS reduces the substrate precondylocarpine acetate to yield an initial, unstable product, which is hypothesised to be dihydroprecondylocarpine acetate. Dihydroprecondylocarpine acetate is proposed to spontaneously desacetoxylate, likely forming dehydrosecodine, which either decomposes or can serve as a substrate for several cyclase enzymes to form distinct alkaloid scaffolds (Figure 1B) ^[5–8]^. We proposed that the reduction of precondylocarpine acetate proceeds through an irreversible 1,4-α,β-unsaturated iminium reduction at C19, but this could only be inferred due to the extensive rearrangement of the reduced product. Although dihydroprecondylocarpine acetate or dehydrosecodine cannot be isolated, a trapped form of the desacetoxylated product, angyline, is sufficiently stable under acidic conditions to be characterised (Figure 1C) ^[5]^. Therefore, to definitively establish the regioselectivity of the DPAS reduction, we performed the enzymatic reaction in the presence of deuterium-labelled coenzyme, 4-pro-*R*-NADPD (Figure 1D). The isolated deuterated angryline was characterised by NMR which clearly showed deuterium incorporation at C19 (Figure S2). The site of incorporation indicates that DPAS catalyses the 1,4 reduction of the α,β-unsaturated iminium moiety and is thereby the first reported ADH with 1,4 reduction activity (Figure 1C).

**Figure 2.**
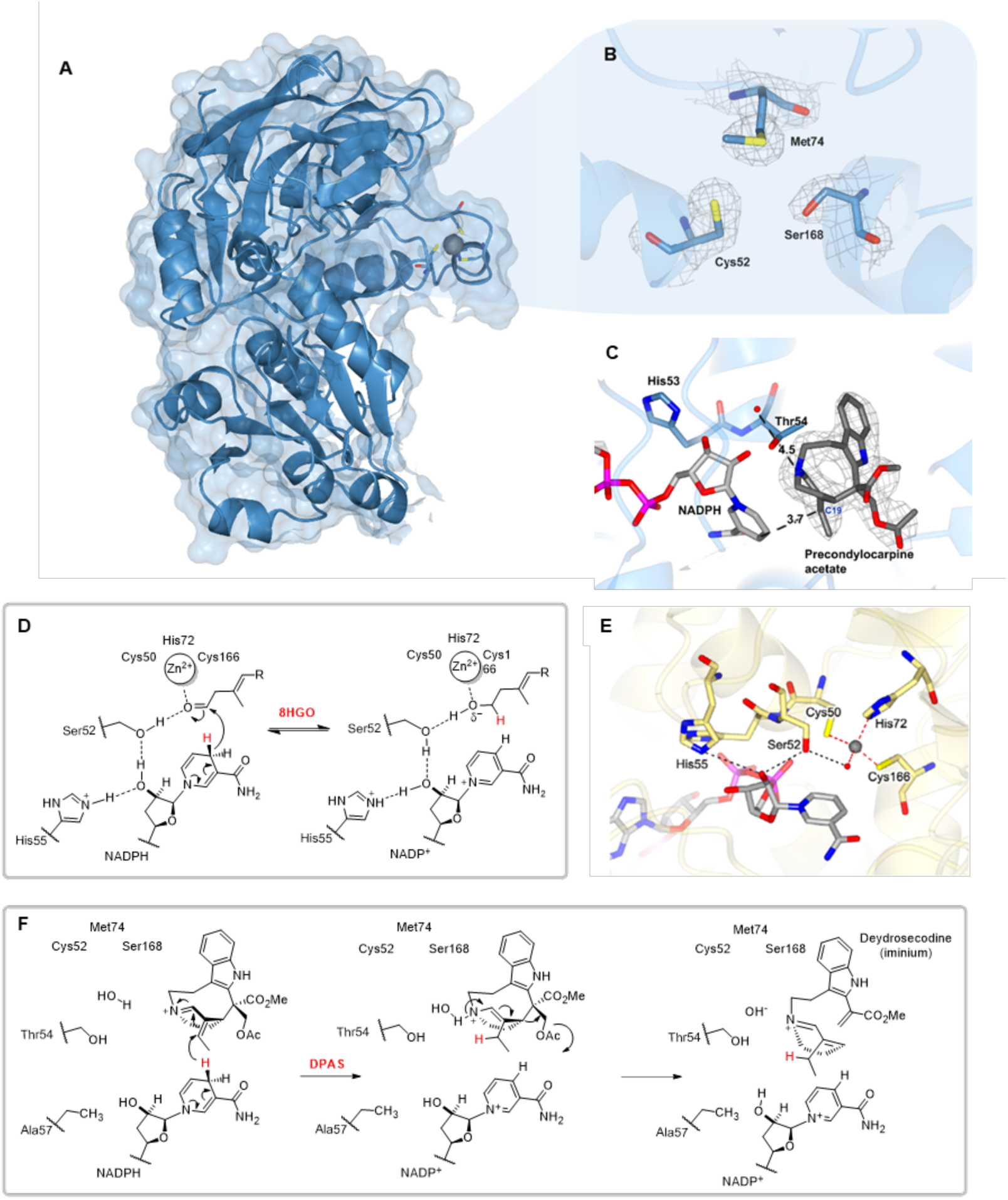
Comparison of DPAS with canonical ADH 8HGO. **A.** Structure of *Ti*DPAS2 structure showing coordination of the structural zinc. **B.** Electron density of *Ti*DPAS2 active site showing lack of catalytic zinc. **C.** Active site of *Ti*DPAS2 showing bound precondylocarpine acetate with docked NADPH. **D.** Proposed mechanism of 8HGO aldehyde reduction. **E.** Active site of *Cr*8HGO with NADP^+^ showing residues involved in catalysis. **F.** Proposed mechanism of 1,4-reduction of an α,β-unsaturated iminium catalysed by DPAS.

Although previous reports have indicated that a dedicated cyclase enzyme is required to form the product vincadifformine ^[9]^, we observed product formation when only DPAS and NADPH is incubated with precondylocarpine acetate (Figure S3). This suggests that after DPAS initially reduces precondylocarpine acetate to form dehydrosecodine, this product can be reduced a second time by DPAS to form secodine, which spontaneously cyclizes to vincadifformine (Figure 1C). Isolation and characterisation of the deuterated vincadifformine product indicate that DPAS catalyses a second 1,4 reduction reaction at C15 of dehydrosecodine (Figure 1C, Figure S4-9). Moreover, CD measurement of the isolated vincadifformine suggest that the product is racemic, consistent with a non-enzymatic cyclization (Figure S10).

**Figure 3.**
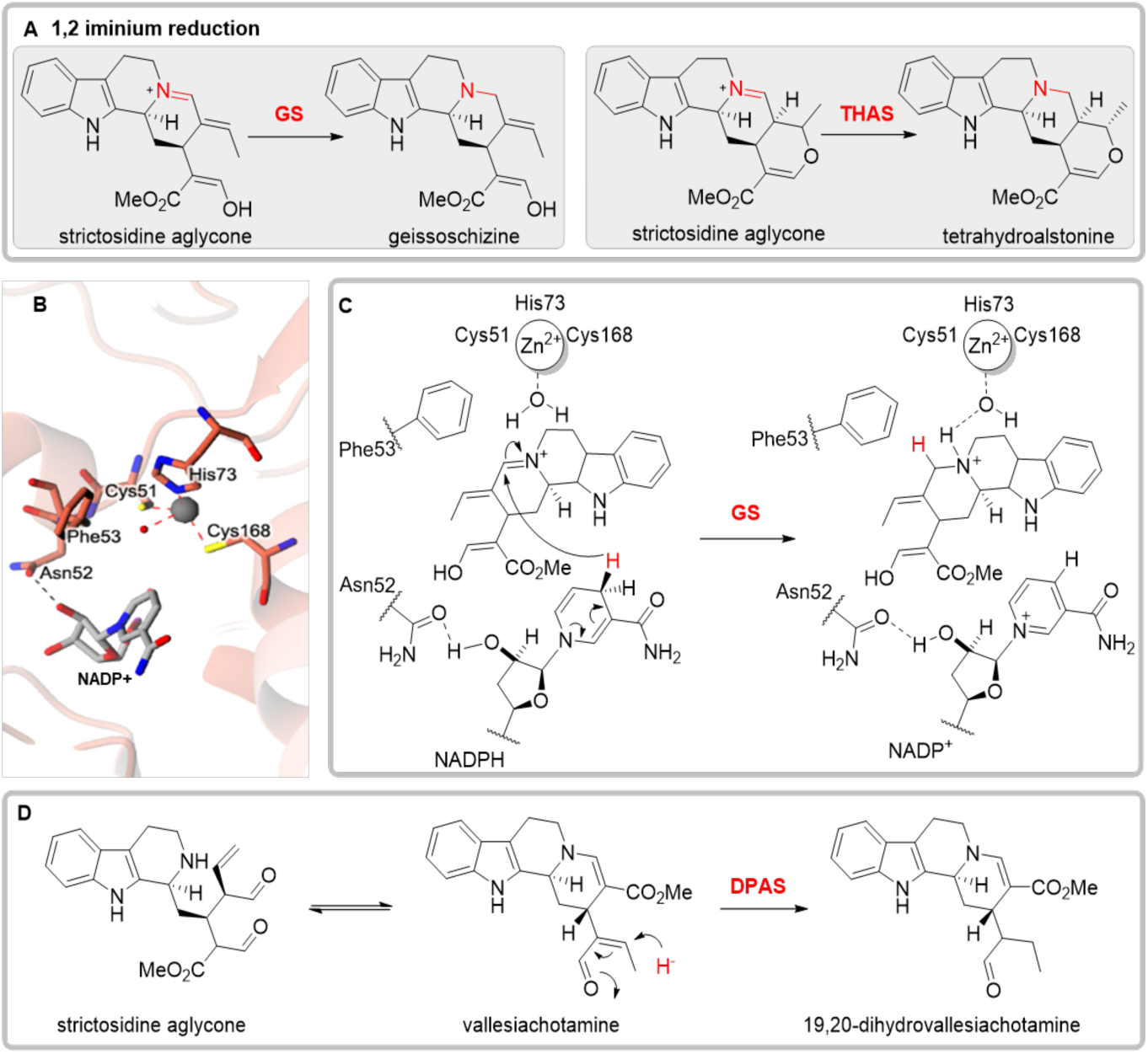
Mechanism of strictosidine aglycone reduction. **A.** 1,2-iminium reductions of strictosidine aglycone catalysed by ADHs GS and THAS. **B.** Active site of GS co-crystallised with NADP^+^ showing residues involved in coordination of the catalytic zinc and loss of the proton relay. **C.** Proposed mechanism of 1,2-iminium reduction catalysed by GS. **D.** Proposed mechanism of 1,4-aldehyde reduction of strictosidine aglycone catalysed by DPAS.

**Figure 4.**
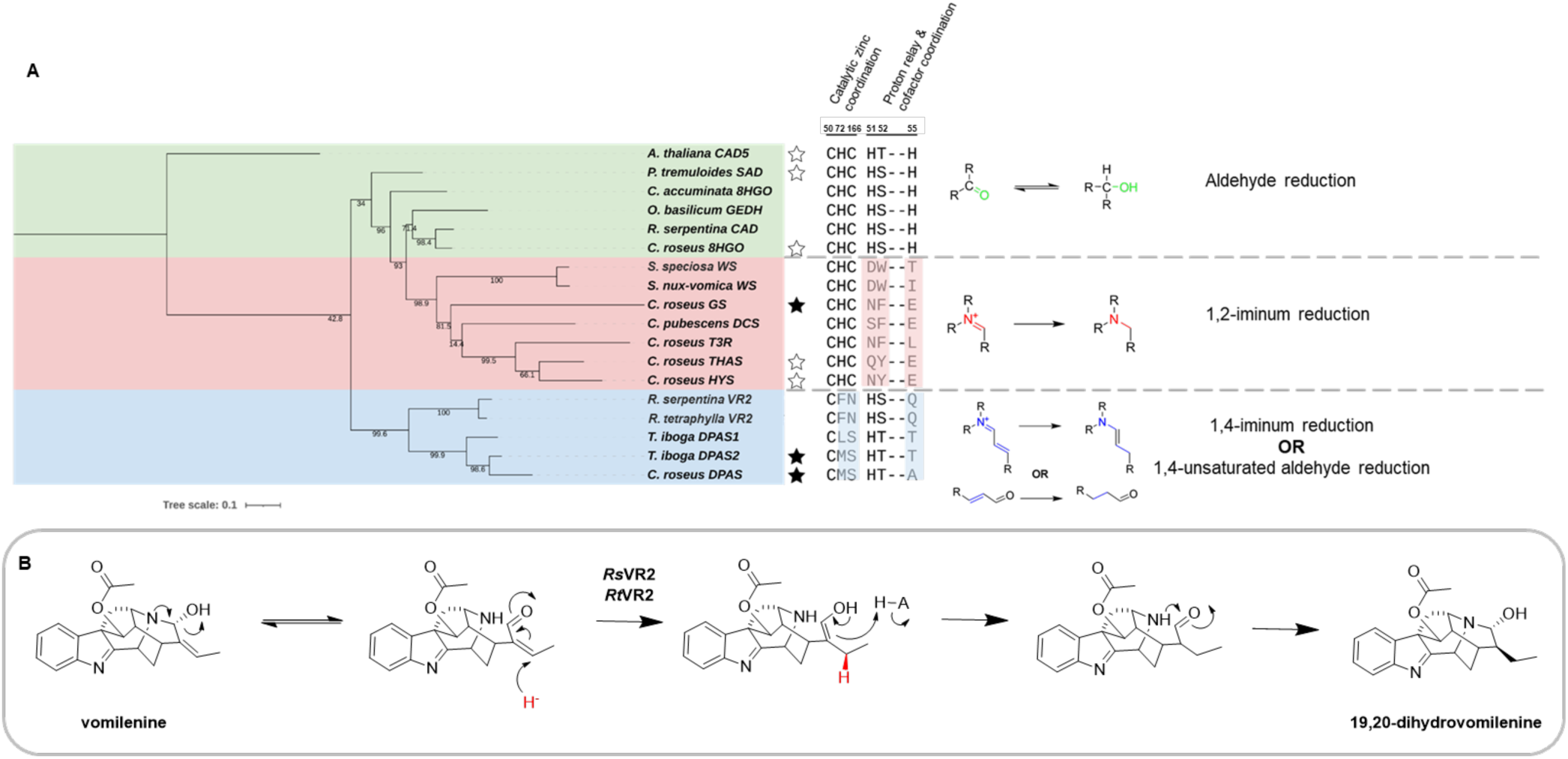
Phylogenetic distribution of non-canonical ADHs in plantae. **A.** Tree of maximum likelihood of previously characterised plant ADHs that perform atypical reductions and conservation of residues coordinating the catalytic zinc and involved in the proton relay. Residue numbering based on *Cr*8HGO structure, stars indicate proteins with structure solved in either previously (line) or in this study (filled). See Table S8 for sequences used. **B.** Proposed mechanism of *Rauwolfia* VR2 1,4 unsatuated aldehyde reduction of vomilenine.

Having established that DPAS catalyses a 1,4 reduction reaction, we solved the crystal structure of this enzyme to understand the structural basis for the switch in catalytic activity compared to canonical 1,2 aldehyde reducing ADHs. The partial structure of DPAS from the vinblastine producing plant *C. roseus* (*Cr*DPAS, 2.16 Å, Figure S11, Table S3), as well as the full structure of the apo- and substrate-bound DPAS from the phylogenetically related alkaloid producing plant *T. iboga* (*Ti*DPAS2, 86% amino acid identity to *Cr*DPAS, 2.42 Å apo-, 1.88 Å precondylocarpine acetate-bound, and 2.24 Å stemmadenine acetate-bound respectively, Figure 2A and 2C, Figure S12, Table S4–6) were solved by X-ray crystallography. Despite the presence of NADP+ in DPAS crystallisation conditions, none of the solved structures had sufficient density for the cofactor. Therefore, to assess the position of the bound substrate relative to the cofactor, NADPH was docked into the active site of DPAS co-crystallised with the substrate precondylocarpine acetate. The 4-pro-*R* hydride of NADPH was measured to be 3.7 Å from C19 of the substrate which is consistent with hydride addition at this carbon (Figure 2C). Additionally, the 4-pro-*R* hydride was 4.6 Å from C15, which is the site of the second reduction to form secodine (Figure 2C), although we note that dehydrosecodine may bind in a different orientation within the substrate cavity (Figure S12).

To establish a point of comparison for DPAS, we used the previously reported structure of a canonical ADH from the vinblastine pathway of *C. roseus,* 8HGO, which catalyses the reversible oxidation/reduction of 8-hydroxygeraniol/8-hydroxygeranial (PDB 6K3G, 60.67% amino acid identity to DPAS, Figure 2D) ^[10]^. The active site of 8HGO contains all of the highly conserved residues involved in ADH catalysis (Figure 2E, Figure S1). Specifically, the catalytic zinc is coordinated by two cysteine thiol groups (Cys50, Cys166), a histidine side chain (His72) and a water molecule ^[4,11]^. The structural zinc - which is distal to the substrate binding pocket and maintains protein domain structure - is tetrahedrally coordinated by four thiol groups (Cys98, Cys106, Cys109, and Cys117). In the canonical ADH reduction mechanism, the aldehyde of the substrate is thought to bind to the catalytic zinc, displacing the water molecule, and the hydride group is transferred from the cofactor to the electrophilic carbon of the aldehyde moiety. The catalytic zinc likely acts as a Lewis acid during catalysis, stabilizing the alkoxide intermediate that forms during the reduction. The reduction is also facilitated by a proton relay system composed of a hydrogen-bonding network between a histidine side chain (His55 in 8HGO), the 2’O ribose of the cofactor, a hydroxyl-group (typically a serine or threonine side chain, Ser52 in 8HGO) and the aldehyde of the substrate (Figure 2D) ^[12]^. The residues involved in binding the catalytic zinc and the proton relay have been demonstrated to be essential for catalysis in numerous studies of ADHs ^[2,3,12,13]^.

Although electron density for the structural zinc ion was clearly observed in all *Ti*DPAS2 structures (the *Cr*DPAS structure lacked electron density for this region), surprisingly, the substrate pocket lacked electron density at the expected site of the catalytic zinc (Figure 2B, Figure S11). This has previously only been reported in a prokaryote ^[14]^. The lack of the catalytic zinc in DPAS is substantiated by the observation that two of the conserved catalytic zinc coordinating residues are mutated (His74Met and Cys168Ser, Figure S1). The zinc ion plays a crucial catalytic role as a Lewis acid in aldehyde reduction. However, the lack of negatively charged intermediate in the 1,4 reduction of an unsaturated iminium system may obviate the need for a Lewis acid.

To investigate the catalytic mechanism of DPAS, we performed site directed mutagenesis of key residues involved in coordination of the catalytic zinc and the proton relay (Figure S13). A DPAS mutant in which the zinc ion coordinating residues were restored (Met74His, Ser168Cys) still reduced precondylocarpine acetate (Figure S14). However, the Met74His mutant had decreased levels of the over reduced vincadifformine product. In contrast, the Ser168Cys mutant showed relative increased vincadifformine production. We speculate that the removal of the catalytic zinc in DPAS enables a larger substrate pocket to better accommodate the substrate. Computational estimation found *Ti*DPAS2 had a larger substrate cavity than other structures of ADHs involved in monoterpene indole alkaloid (MIA) biosynthesis (Figure S15). The change in the ratio between the single (angryline) and the double (vincadifformine) reduced product in these mutants could be due to modulating the shape of the binding pocket and the manner in which the substrates bind. Due to the instability of the substrate and products, accurate product quantification required for steady-state kinetics was impractical, so only qualitative conclusions were drawn from these mutational analyses.

Canonical ADHs rely on a proton relay enabled by conserved Ser/Thr and His residues (Figure 2D). The hydroxyl moiety of Thr54 in TiDPAS (corresponding to Ser52 in 8HGO) was 3.6 Å from the nitrogen of the substrate’s iminium suggesting that this residue may act in the proton relay (Figure 2C). However, while enzyme activity was abolished in a Thr54Phe mutant, the Thr54Ala mutant was active (Figure S14). Therefore, while the size of this residue may impact substrate binding, a proton relay involving a hydroxyl group is not essential for catalysis. The γN of a highly conserved histidine which typically acts as a proton donor during catalysis (His55 in 8HGO, Figure 2D), is mutated to Ala57 in *Cr*DPAS and Thr57 in *Ti*DPAS2. We speculated a nearby histidine (His53) may be acting as a proton donor, however His53Ala mutation did not affect DPAS activity, indicating this residue is not essential for catalysis of the 1,4 reduction reaction (Figure S14). Collectively, these findings suggest the 1,4-α,β-unsaturated iminium reduction by DPAS does not require a catalytic zinc or a proton relay, but reintroduction of these features did not abolish activity (Figure 2F). The inherent reactivity of the unsaturated iminium substrate compared to aldehyde substrates means that the enzyme may not require a Lewis acid to stabilize the reaction as the reduction proceeds. A number of ordered water molecules in the DPAS structure could take over the role of proton donor (Figure 2C). Correct positioning of the substrate relative to the NADPH cofactor may be sufficient for reduction of this already activated substrate.

In addition to the 1,4 reduction catalysed by DPAS, one additional non-canonical reduction reaction has been reported for eukaryotic ADHs: the 1,2 reduction of an iminium moiety. GS catalyses the irreversible 1,2 iminium-reduction of strictosidine aglycone to form geissoschizine, another intermediate in vinblastine biosynthesis (Figure 3A) ^[8,15]^. Additionally, tetrahydroalstonine synthase (THAS) is an 1,2 iminium reductase that reduces an alternate isomer of strictosidine aglycone to generate the alkaloid tetrahydroalstonine (Figure 3A) ^[16,17]^. The apo- and holo structures of THAS were previously solved (PDB accessions 5FI3 and 5H83), but the mechanistic basis behind the switch in activity from aldehyde to iminium reduction remains poorly understood. In light of the insights from the DPAS structure, we solved the holo structure of GS to 2.00 Å resolution (PDB 8A3N, Figure S16, Table S7) and compared the THAS and GS structures with the 1,4 reduction mechanism of precondylocarpine acetate.

GS and THAS (47.7% and 53.8% amino acid identity to DPAS respectively) each crystallised with both the catalytic and structural zinc ions bound. Mutagenesis of GS residues involved in the coordination of the catalytic zinc (His73 and Cys168) resulted in the abolishment of GS activity, suggesting that, as in the canonical ADH mechanism, the catalytic zinc is essential for 1,2 reduction of the iminium (Figure S17). Additionally, GS and THAS each had mutations in highly conserved residues involved in the proton relay observed in canonical ADH reduction (Figure 3B, Figure S1). THAS was previously postulated to use Tyr56 (corresponding to Ser52 in 8HGO) in the proton relay ^[16]^ but careful inspection of the THAS structure in the context of this study suggests that the hydroxyl group may not be appropriately positioned for catalysis. Moreover, GS contains a Phe residue in this position (Phe53). No substantial change in reduction activity was observed for GS Phe53Tyr mutant or the corresponding THAS Tyr56Phe mutant, suggesting the hydroxyl-group has no catalytic role (Figure S18). GS and THAS also both lack the highly conserved histidine residue involved in the proton relay (His55 in 8HGO replaced with Glu56 in GS and Glu59 in THAS). Although THAS Glu59 is positioned 3.6 Å from the 2’O of bound-NADP^+^, GS Glu56 is positioned 7.6 Å from this moiety which is not consistent with a role for this residue in cofactor binding or catalysis (Figure 3B). This residue has been previously reported to affect the stereoselectivity of the reduction in THAS, likely due to the orientation of the cofactor in the substrate pocket ^[16]^. These findings indicate the proton relay present in canonical ADHs is not required for iminium 1,2-reduction. Given that iminium moieties are typically more activated than aldehydes in the context of reduction, the proton relay found in canonical aldehyde reducing ADHs may not be required. We hypothesise that the water molecule in the fourth coordination position of the catalytic zinc remains bound upon substrate binding. This water could hydrogen bond to the substrate iminium, serving as the proton donor, with the zinc ion acting as a Lewis base (Figure 3C, Figure S16).

GS has been reported to be able to reduce precondylocarpine acetate, albeit it at substantially reduced efficiency compared to DPAS (Figure S19) ^[8]^. This suggests that the 1,2 iminium reducing ADHs can also catalyse a 1,4 reduction provided the substrate can productively bind in the active site. Conversely, assay of DPAS with strictosidine aglycone, the substrate of GS and THAS, showed that DPAS could also turnover this substrate (Figure S20). However, NMR characterisation of the major product revealed a previously unreported compound that we named 19,20-dihydrovallesiachotamine (Figure S21–38). This structure indicates that DPAS catalyses the 1,4 reduction of the α,β-unsaturated carbonyl in strictosidine aglycone at C19 (Figure 3D, Figure S12). Assay of strictosidine aglycone with DPAS mutants showed that removal of the hydroxyl moiety of Thr54 – either by mutating to Phe or Ala – abolished enzyme activity. This suggests DPAS Thr54 has a catalytic role in the presence of less electronically activated substrate strictosidine aglycone. Additionally, mutation of the residues corresponding to catalytic zinc binding and proton relay in DPAS resulted in either loss of activity with strictosidine aglycone, or 1,2-iminium reduction of an alternative rearrangement of strictosidine aglycone to form tetrahydroalstonine (Figure S39). Mutation of these residues likely impact substrate binding, highlighting the plasticity of the active site of this enzyme. Increased substrate promiscuity within the ADH active site may be observed in these cases since certain catalytic requirements– zinc ion and proton relay– are not strictly required for the reduction of these substrates.

Phylogenetic analysis of previously characterised plant ADHs revealed three clades that grouped according to conservation of residues involved in the coordination of the catalytic zinc, and residues involved in proton relay and cofactor coordination (Figure 4A). Interestingly, we found the predicted catalytic activity based on residues involved in the coordination of the catalytic zinc and proton relay discussed in this study largely correlated with the experimentally verified activities of these enzymes (Figure S40. The phylogenetic distribution of these atypical ADHs is limited to the Apocynaceae, Loganiaceae and Rubiaceae families – plant families known to produce MIAs. However, this distribution may increase as more enzymes involved in plant secondary metabolism are discovered. The identification of sequence motifs may allow better predictions of enzyme function and mechanism. For example, the previously reported vomilenine reductase 2 (VR2) from *Rauwolfia* (a member of the Apocynaceae family) is in the same phylogenetic clade as DPAS (66% amino acid identity) and is also missing the conserved residues important in coordinating the catalytic zinc (Phe73 instead of His and Asn167 instead of Cys) and the proton relay (Gln56 instead of His) ^[18]^. After inspection of the substrate and product of VR2, we propose a mechanism for VR2 based on the mechanism of DPAS that involves a 1,4 unsaturated aldehyde reduction at C19 (Figure 4B).

ADHs are an ancient and widespread protein family that until recently had a limited reported catalytic repertoire. Here we show the structural basis of how ADHs can catalyse two distinct variations on carbonyl reduction. These enzymes make a valuable addition to other known 1,2-imine reductases (IREDs) ^[19]^. Furthermore, these findings demonstrate how ADH function has been co-opted to perform atypical chemistry by mutations in key catalytic residues, enabling various biotechnological and gene discovery applications, and sheds light on the evolution of chemical diversity in plants.

## Acknowledgements

We gratefully acknowledge the Max Planck Society and the ERC (788301) for funding. ET is financially supported by National Natural Sciences Foundation of China, Research Fund for International Excellent Young Scientists (Grant 32150610477).

## Supporting Information

### Experimental Procedures

#### Chemicals and molecular biology reagents

All solvents used for extractions, chemical synthesis and preparative HPLC were HPLC grade, and solvents used for UPLC/MS were MS grade. All solvents were purchased from Sigma Aldrich. Carbenicillin, kanamycin sulfate, isopropyl β-D-thiogalactoside (IPTG) salts were purchased from Sigma. Synthetic genes were purchased from IDT. All gene amplifications and mutations were performed using Platinum II Superfi DNA Polymerase (Thermo Fisher). Constructs were transformed into vectors using In-Fusion kit (ClonTech Takara) and colony PCR was performed using Phire II mastermix (Thermo Fisher) according to manufacturer’s instructions. PCR product purification was performed using Zymoclean Gel DNA Recovery kit (Zymo). Plasmid purification was performed using the Wizard Miniprep kit (Promega). Strictosidine, precondylocarpine acetate, stemmadenine acetate, angryline, vincadifformine, 19-*E*-geissoschizine and tetrahydroalstonine were enzymatically prepared and purified as previously described ^[1–4]^.

#### Cloning and mutagenesis

Cloning of *Cr*DPAS, *Ti*DPAS1, *Ti*DPAS2, *Cr*GS and *Cr*THAS has been previously reported ^[1,2,4,5]^. Full-length *Cr*DPAS, *Ti*DPAS2, GS and THAS were amplified by PCR from the codon optimized synthetic genes listed in Table S2 using corresponding primers listed in Table S1. *Thermoanaerobacter brockii* alcohol dehydrogenase (*Tb*ADH) synthetic gene (Table S2) was cloned into the pOPINF vector. DPAS, GS and THAS mutants were generated by overlap extension PCR as previously reported ^[6]^. PCR products were purified from 1% agarose gel and ligated into the BamHI and KPNI restriction sites of pOPINK vectors for small-scale GS and GS mutants. All other ADHs were cloned into pOPINF vector. pOPINF and pOPINK were a gift from Ray Owens (Addgene plasmid # 26042 and # 41143 ^[7]^). Constructs were ligated into vectors using the In-Fusion kit (Clontech Takara).

#### Expression and purification of proteins in *E. coli*

Constructs were transformed into chemically-competent *E. coli* Stellar cells (Clontech Takara) by heat shock at 42°C for 30 seconds and selected on LB agar containing 50μg/mL carbenicillin or kanamycin for pOPINF or pOPINK constructs respectively. Positive colonies were screened by colony PCR using primers listed in Table S1 and grown overnight at 37°C shaking at 200 r.p.m. Plasmids were then isolated and constructs were sequence verified. Plasmids were transformed into chemically competent *E. coli* SoluBL21 cells by heat shock for 30 seconds at 42°C and selected on LB agar containing 50 μg/mL carbenicillin or kanamycin for pOPINF or pOPINK constructs respectively. For small scale protein purification, 10 mL starter cultures of LB with 50 μg/mL of the respective antibiotic and a colony of transformed construct in SoluBL21 cells were grown at 37°C 200 r.p.m. overnight. Media (100 mL 2xYT media) containing 50 μg/mL antibiotic was inoculated with 1 mL of the starter culture and grown until OD_600_ of 0.6 was reached. For large scale purification, 20 mL starter cultures of LB with antibiotic and a colony of transformed construct in SoluBL21 cells were grown at 37°C 200 r.p.m. overnight. Media (1L 2xYT media) containing 50 μg/mL carbenicillin was inoculated with 10 mL of starter culture and grown until OD600 of 0.6 was reached. Once cultures had reached the desired OD_600_, cultures were transferred to 18°C 200 r.p.m shaking incubator for 30 minutes before protein expression was induced by addition of 300 μM IPTG, after which cultures were grown for an additional 16 hours.

#### *Cr*PAS insect cell expression

N-terminal His_6_-tagged *Cr*PAS was expressed in Sf9 insect cells as previously described ^[1]^. Cells were harvested by centrifugation and the pellets frozen at –80°C until large-scale purification.

#### *Cr*DPAS, *Cr*GS and *Cr*THAS small-scale protein expression and purification

Cells were harvested by centrifugation at 4000 × *g* for 15 minutes and re-suspended in 10 mL buffer A1 (50 mM Tris-HCl pH 8, 50 mM glycine, 500 mM NaCl, 5% glycerol, 20 mM imidazole) with addition of EDTA-free protease inhibitor cocktail (Roche Diagnostics Ltd.) and 10 mg lysozyme (Sigma). Cells were lysed at 4°C using a sonicator (40% amplitude, 2 seconds on, 3 seconds off cycles for 2 minutes) and centrifuged at 35000 × *g* to remove insoluble cell debris. The supernatant was collected and filtered with 0.2 um PES syringe filter (Sartorious) and purified by addition of 150 μL washed Ni-NTA agarose beads (QIAGEN). Samples were incubated on a rocking incubator at 4°C for 1 hour. Beads were washed by centrifuging at 1000 × *g* for 1 minute to remove the supernatant, and then the beads were resuspended in 10 mL of A1 Buffer. This step was performed a total of three times. Protein was eluted by resuspending the beads in 600 μL of buffer B1 (50 mM Tris-HCl pH 8.0, 50 mM glycine, 500 mM NaCl, 5% glycerol, 500 mM imidazole) before centrifuging for 1000 × *g* for 1 minute and then collecting the supernatant. This elution step was repeated to remove all Ni-NTA bound protein. Proteins were buffer exchanged into buffer A4 (20 mM HEPES pH 7.5, 150 mM NaCl) and concentrated using 10K Da molecular weight cut off centrifugal filter (Merck) and stored at –80°C.

#### *Cr*DPAS, *Ti*DPAS2, *Cr*GS, *Cr*SGD, *Cr*PAS and *Tb*ADH large-scale purification

Cells were harvested by centrifugation at 3200 × *g* for 15 minutes and re-suspended in 50 mL buffer A1 (50 mM Tris-HCl pH 8, 50 mM glycine, 500 mM NaCl, 5% glycerol, 20 mM imidazole) with addition of EDTA-free protease inhibitor cocktail (Roche Diagnostics Ltd.) and 10 mg lysozyme (Sigma). Dithiothreitol (Sigma) (final concentration of 0.05 mM) was additionally added to all buffers in purification of *Cr*DPAS, *Ti*DPAS2 and *Cr*GS for crystallisation. Cells were lysed at 4°C using a cell disruptor at 30 KPSI and centrifuged (35000 × *g*) to remove insoluble cell debris. The supernatant was collected and filtered with 0.2 μm PES syringe filter (Sartorious) and purified using an AKTA pure FPLC (Cytiva). Sample was applied at 2 mL/min onto a His-Trap HP 5mL column (Cytiva) and washed with 5 column volumes (CV) of buffer A1 before being eluted with 5 CV of buffer B1. Protein was detected and collected using the UV 280 nm signal and then further purified on a Superdex Hiload 16/60 S200 gel filtration column (Cytiva) at a flow rate of 1 mL/min using buffer A4. Proteins were finally buffer exchanged into buffer A4 and concentrated using 10K Da molecular weight cut off centrifugal filter (Merck) before being snap frozen in liquid nitrogen and stored at −80°C.

For crystallisation of *Cr*DPAS,*Ti*DPAS2 *and Cr*GS, protein after gel filtration was incubated on a rocker overnight at 4°C with 3C protease to cleave the 6xHis-tag. Proteins were then passed through a 1mL HisTrap column (Cytiva) to remove the cleaved tag. Proteins were then buffer exchanged into buffer A4 (20 mM HEPES pH 7.5, 150 mM NaCl) containing 0.05 mM tris(2-carboxyethyl)phosphine (Sigma) and concentrated using 10K Da molecular weight cut off centrifugal filter (Merck) and stored at – 80°C.

#### Synthesis of NADPD

Deuterated pro-*R*-NADPD was produced in vitro as previously described ^[8]^ with minor modifications. A 20 mL reaction mixture containing 2 mM NADP^+^, 4 mM d_8_-isopropanol, 1 mM semicarbazide and 5 μM *Tb*ADH in 50 mM ammonium bicarbonate buffer at pH 7.5 was incubated at 30°C. The progression of the reaction was monitored spectrophotometrically at 340 nm. When no significant increase in absorbance was observed (approximately 3 hours), 300 μL of Ni-NTA agarose beads (Qiagen) was added and the sample incubated rocking at room temperature for 30 minutes. The reaction was centrifuged to remove the Ni-NTA beads bound to *Tb*ADH, and the supernatant was filtered through a 45 μm glass filter and lyophilized to remove the unreacted d_8_-isopropanol, the acetone that forms during the reaction and the buffer. The residue, containing primarily NADPD, was stored at –20°C until use.

#### In vitro enzyme assays

Enzymatic assays with precondylocarpine acetate were performed in 50 mM HEPES buffer (pH 7.5) with 50 μM precondylocarpine acetate in MeOH (not exceeding 5% of the reaction volume), 250 μM NADPH cofactor (Sigma) and 150 nM enzyme to a final reaction volume of 100 μL. Reactions were incubated for 30 minutes at 30°C and shaking at 60 r.p.m. before being quenched with 1 volume of 70% MeOH with 1% H_2_CO_2_. Enzymatic assays with strictosidine aglycone were performed in 50 mM HEPES buffer (pH 7.5), 100 μM strictosidine and 1 mM SGD to a final reaction volume of 100 μL. Assays were incubated for 30 minutes at 30°C and shaking at 60 r.p.m before 500 nM of ADH enzyme and 250 μM NADPH was added. As control, the reactions were performed without the addition of ADH enzyme. Reactions were incubated for a further 30 minutes at 30°C shaking at 60 r.p.m. before being quenched with 1 volume of 70% MeOH with 0.1% H_2_CO_2_. All enzymatic assays were centrifuged at 14000 × *g* for 15 minutes and the supernatant analysed by UPLC-MS.

#### UPLC-MS

All assays were analysed using a Thermo Scientific Vanquish UPLC coupled to a Thermo Q Exactive Plus orbitrap MS. For assays using precondylocarpine acetate, chromatographic separation was performed using a Phenomenex Kinetex C18 2.6 μm (2.1 × 100 mm) column using water with 1% H_2_CO_2_ as mobile phase A and acetonitrile with 1% H_2_CO_2_ as mobile phase B. Compounds were separated using a linear gradient of 10-30% B in 5 minutes followed by 1.5 minutes isocratic at 100% B. The column was then re-equilibrated at 10% B for 1.5 minutes. The column was heated to 40°C and flow rate was set to 0.6 mL/min. For assays using strictosidine aglycone, separation was carried out using a Waters Acquity BEH C18 1.7 μm (2.1 × 50 mm) using 0.1% NH_4_OH in water as mobile phase A and acetonitrile as mobile phase B. Compounds were separated using a linear gradient of 10-90% B in 9 minutes followed by 2 minutes isocratic at 90% B. The column was re-equilibrated at 10% B for 3 minutes. The column was heated to 50°C and flow rate was set to 0.4 mL/min. MS detection was performed in positive ESI under the following conditions: spray voltage was set to 3.5 kV ~ 67.4 µA, capillary temperature set to 275°C, vaporizer temperature 475°C, sheath gas flow rate 65, sweep gas flow rate 3, aux gas flow rate 15, S-lens RF level to 55 V. Scan range was set to 200 - 1000 *m/z* and resolution at 17500.

#### Production and isolation of *d*-angryline and *d_2_*-vincadifformine

*d*-angryline was produced enzymatically from stemmadenine acetate using the same protocol previously described for the synthesis of angryline but replacing NADPH with NADPD ^[6]^. Briefly, 0.25 mg of stemmadenine acetate, 40 μM flavin adenine dinucleotide (FAD) and 5 μg of *Cr*PAS were combined in a total volume of 500 μL in 50 mM TRIS-HCl buffer pH 8.5 and incubated at 37°C to form precondylocarpine acetate (reaction progress was monitored by LC-MS, *m/z* 395.19). After 2 hours, 1 mg of NADPD and 9 μg of *Cr*DPAS were added to the reaction and incubated for 20 minutes at 37 °C to obtain *d*-angryline (*m/z* 338.19). Multiple reactions were prepared to obtain sufficient product for NMR characterization. After completion, the reactions were snap frozen in liquid nitrogen and stored at –80 °C.

*d_2_*-vincadifformine was also produced enzymatically, but in this case NADPD was generated directly in the reaction mixture using an alcohol dehydrogenase from *E. coli* (Merck product 49854). Multiple 500 μL reactions were prepared to obtain sufficient product for NMR characterization. Each reaction contained 400 μM NADP^+^, 0.89 μg *d_8_*-isopropanol, 1 μg of *Tb*ADH, 10 μg stemmadenine acetate, 0.8 μM *Cr*PAS and 0.8 μM DPAS in 50 mM HEPES buffer pH 7.5. The reactions were incubated at 30 °C for 1 hour, snap frozen in liquid nitrogen and stored at –80 °C until purification of the final product.

*d*-angryline and *d_2_*-vincadifformine were purified by semi-preparative HPLC on an Agilent 1260 Infinity II HPLC system. The reactions were thawed and 500 μL of 90:9:1 MeOH:H_2_O:H_2_CO_2_ was added to the deuterated samples. The samples were filtered through 0.2 μm PTFE disc filters (Sartorius) to remove the precipitated enzymes and injected onto a Phenomenex Kinetex XB-C18 5 μm (250 × 10 mm) column. Chromatographic separation was performed using 0.1% H_2_CO_2_ in water as mobile phase A and acetonitrile as mobile phase B. A linear gradient from 10% B to 40% B in 15 minutes was used for chromatographic separation of the compounds followed by a wash at 40% B for 5 minutes and a re-equilibration step to 10% B for 5 minutes. Flow rate was 6 mL/min. Elution of *d*-angryline and *d_2_*-vincadifformine was monitored at two wavelengths, 330 and 254 nm. Fractions containing the compounds of interest were collected, dried under reduced pressure and stored at –80 °C until further analysis.

#### Production and isolation of 19,20-dihydrovallesiachotamine

19,20-dihydrovallesiachotamine was produced enzymatically from 100 μM strictosidine reacted with 100 μM *Cr*SGD in 50 mM HEPEs buffer pH 7.5 in a 100 mL reaction at 30°C. After 90 minutes, 500 nM of *Cr*DPAS and 250 μM NADPH was added and the reaction monitored. After 2 hours a further 500 nM *Cr*DPAS was added to a final concentration of 1 μM and left for a further 3 hours until the reaction reached completion. The sample was snap frozen in liquid nitrogen and stored at –80 °C. For purification, the sample was thawed on ice and filtered through a 0.2 μm PTFE disc filter (Sartorius) to remove the precipitated enzymes and then passed through a Supelco DSC-18 column (MilliporeSigma) and eluted with methanol. Eluent was dried down in a rotovap and resuspended in 1.5 mL methanol. The product was purified on an Agilent 1290 Infinity II semi-preparative HPLC system using a Waters XBridge BEH C18 5 μm (10 × 250mm) column and using 0.1% NH_4_OH in water as mobile phase A and acetonitrile as mobile phase B. Compounds were separated using a linear gradient of 10-65% B in 25 minutes followed by 10 minutes column re-equilibration at 10% B. Flow rate was set to 7mL/min. Compound was detected by measuring UV 290 nm and 254 nm signal. Fractions containing the compound of interest were collected and dried down using a rotovap and stored at –20 °C until NMR analysis.

#### NMR of *d*-angryline, *d*-vincadifformine and 19,20-dehydrovallesiachotamine

For *d*-angryline, NMR spectra were measured on a 400 MHz Bruker Advance III HD spectrometer (Bruker Biospin GmbH, Rheinstetten, Germany). NMR spectra for 19,20-dehydrovallesiachotamine, (–)-vincadifformine and *d_2_*-(±)-vincadifformine were measured on a 700 MHz Bruker Advance III HD spectrometer (Bruker Biospin GmbH, Rheinstetten, Germany). For spectrometer control and data processing Bruker TopSpin ver. 3.6.1 was used. MeOH-*d_3_* was used as a solvent and all NMR spectra were referenced to the residual solvent signals at δH 3.31 and δC 49.0, respectively.

#### ECD measurement

ECD spectra were measured at 25 °C on a JASCO J-810 spectropolarimeter (JASCO cooperation, Tokyo, Japan) using a 350 µL cell. Spectrometer control and data processing was accomplished using JASCO spectra manager II.

#### ECD spectral calculations for (-)-vincadifformine

Based on the structure determined from NMR analysis a molecular model was created in GaussView ver.6 (Semichem Inc., Shawnee, Kansas, USA) and optimized using the semi-empirical method PM6 in Gaussian (Gaussian Inc., Wallingford, Connecticut, USA). The resulting structure was used for conformer variation with the GMMX processor of the Gaussian program package. Resulting structures were DFT-optimized with Gaussian ver.16 (APFD/6-31G(d)). A cut-off level of 4 kcal/mol was used to select conformers which were subjected to another DFT optimization on a higher level (APFD/6-311G+(2d,p)). All structures up to a deviation of 2.5 kcal/mol from the lowest energy conformer were used to determine the ECD-frequencies in a TD-SCF calculation on the same level as the former DFT optimization. The ECD curve was calculated from the Boltzmann-weighed contributions of all conformers with a cut-off level of two percent.

Experimentally measured ECD data and calculated data were compared using SpecDis ver.1.71 ^[9]^.

#### Protein crystallisation

Purified *Cr*DPAS and *Ti*DPAS2 were crystallised by sitting-drop vapour diffusion on MRC2 96-well crystallisation plates (SwissSci) with 0.3 uL protein and 0.3 uL precipitant solution drops dispensed by Oryx8 robot (Douglas Instruments). *Cr*DPAS was crystallised using JCSG screen (Jena Biosciences) with 1.26 M ammonium sulfate, 100 mM TRIS buffer pH 8.5 and 200mM lithium sulfate. Crystallisation condition with additional 1 mM NADP^+^ and 25% ethylene glycol was used as cryoprotectant. *Ti*DPAS2 was initially screened using PEG/Salt screen (Jena Biosciences) before condition optimization. Apo-*Ti*DPAS2 was crystallised in 17% w/v PEG 3350, 200 mM ammonium chloride and 0.75 mM angryline (no electron density corresponding to angryline was observed in the structure). 17% w/v PEG 3350, 220 mM ammonium chloride, 1 mM NADP^+^, 1 mM angryline and 25% ethylene glycol was used as cryoprotectant. Stemmadenine acetate-bound *Ti*DPAS2 was crystallised in 23% w/v PEG 3350, 250 mM sodium sulfate and 0.75 mM stemmadenine acetate, 23% w/v PEG 3350, 200 mM sodium sulfate, 1 mM NADP^+^, 1 mM stemmadenine acetate and 25% ethylene glycol was used as cryoprotectant. Precondylocarpine acetate-bound *Ti*DPAS2 was crystallised in 25% w/v PEG 3350, 180 mM sodium sulfate and 0.75 mM precondylocarpine acetate. 23% w/v PEG 3350, 200mM sodium sulfate, 1 mM NADP^+^, 1 mM precondylocarpine acetate and 25% ethylene glycol was used as cryoprotectant. *Cr*GS was crystallised in 25% w/v PEG 3350, 100 mM TRIS buffer pH8.0; 20% v/v ethylene glycol was added to this condition for the cryoprotectant. All crystals were soaked in the corresponding cryoprotectant before flash-cooling in liquid nitrogen.

#### X-ray data collection, processing and structure solution

X-ray data sets for *Cr*DPAS and *Ti*DPAS2 structures were recorded on the 10SA (PX II) beamline at the Paul Scherrer Institute (Villigen, Switzerland) at wavelength of 1.0 .Å using a Dectris Eiger3 16M detector with the crystals maintained at 100K by a cryocooler. Diffraction data were integrated using XDS ^[12]^ and scaled and merged using AIMLESS ^[14]^; data collection statistics are summarized in Table S3–7. Structure’s solution was automatically obtained by molecular replacement using the structure of tetrahydroalstonine synthase from *C. roseus* (PDB accession code 5FI3) as template with which *Cr*DPAS and *Ti*DPAS2 share 54% and 56% amino acid identity respectively. In all cases the map was of sufficient quality to enable 90% of the residues expected for a homodimer to be automatically fitted using Phenix autobuild ^[10]^. The models were finalized by manual rebuilding in COOT ^[11]^ and refined using in Phenix refine.

X-ray data for *Cr*GS was recorded at 100 K on beamline I03 at the Diamond Light Source (Oxfordshire, UK) using a Pilatus3 6M hybrid photon counting detector (Dectris), with crystals maintained at 100 K by a Cryojet cryocooler (Oxford Instruments). Diffraction data were integrated and scaled using XDS ^[12]^ via the XIA2 expert system ^[13]^ then merged using AIMLESS ^[14]^ A summary of the data processing is presented in Table S7. A template for molecular replacement was prepared with CHAINSAW ^[15]^ from the structure of tetrahydroalstonine synthase from *C. roseus* (PDB accession code 5FI3) with which *Cr*GS shares 57% amino acid sequence identity. The structure was solved by molecular replacement using PHASER ^[16]^, giving two copies of the subunit in the asymmetric unit, which formed the homodimeric assembly expected for this class of enzyme. After restrained refinement in REFMAC5 ^[17]^ at 2.0 Å resolution, the protein component of the model was completely rebuilt using BUCCANEER ^[18]^. The model was finalized after several iterations of manual editing in COOT ^[11]^ and further refinement in REFMAC5 incorporating TLS restraints. The model statistics are reported in Table S8.

All structures are in the PDB database under the following accessions: X (*Cr*DPAS), X (apo-*Ti*DPAS2), X (precondylocarpine acetate-bound *Ti*DPAS2), X (stemmadenine acetate-bound *Ti*DPAS2), 8A3N (*Cr*GS).

#### Docking simulations

Ligands were docked into the active site of *Ti*DPAS and *Cr*GS using AutoDock Vina on the Webina webserver using default parameters ^[19,20]^. Coordinates of ligands were generated by PDBQTConvert. When assessing the results, we selected ligand orientations in which the 4-pro-*S* hydride of NADPH was in close proximity to the carbon being reduced; this orientation was not always the lowest possible energy solution. Results were visualised using PyMOL. Cavity pocket size estimation was computed using CASTp3.0 using default parameters ^[21]^. Results were visualised using Chimera.

#### Phylogenetic analysis

Nucleic acid sequences of ADH genes were aligned using MUSCLE5 ^[22]^. A maximum likelihood phylogenetic tree was constructed using IQTree using a best-fit substitution model followed by tree reconstruction using 1000 bootstrap alignments and the remaining parameters used default settings ^[23]^. Tree visualisation and figures were made using iTOL version 6.5.2 ^[24]^.

## Supporting Figures

**Figure S1.**
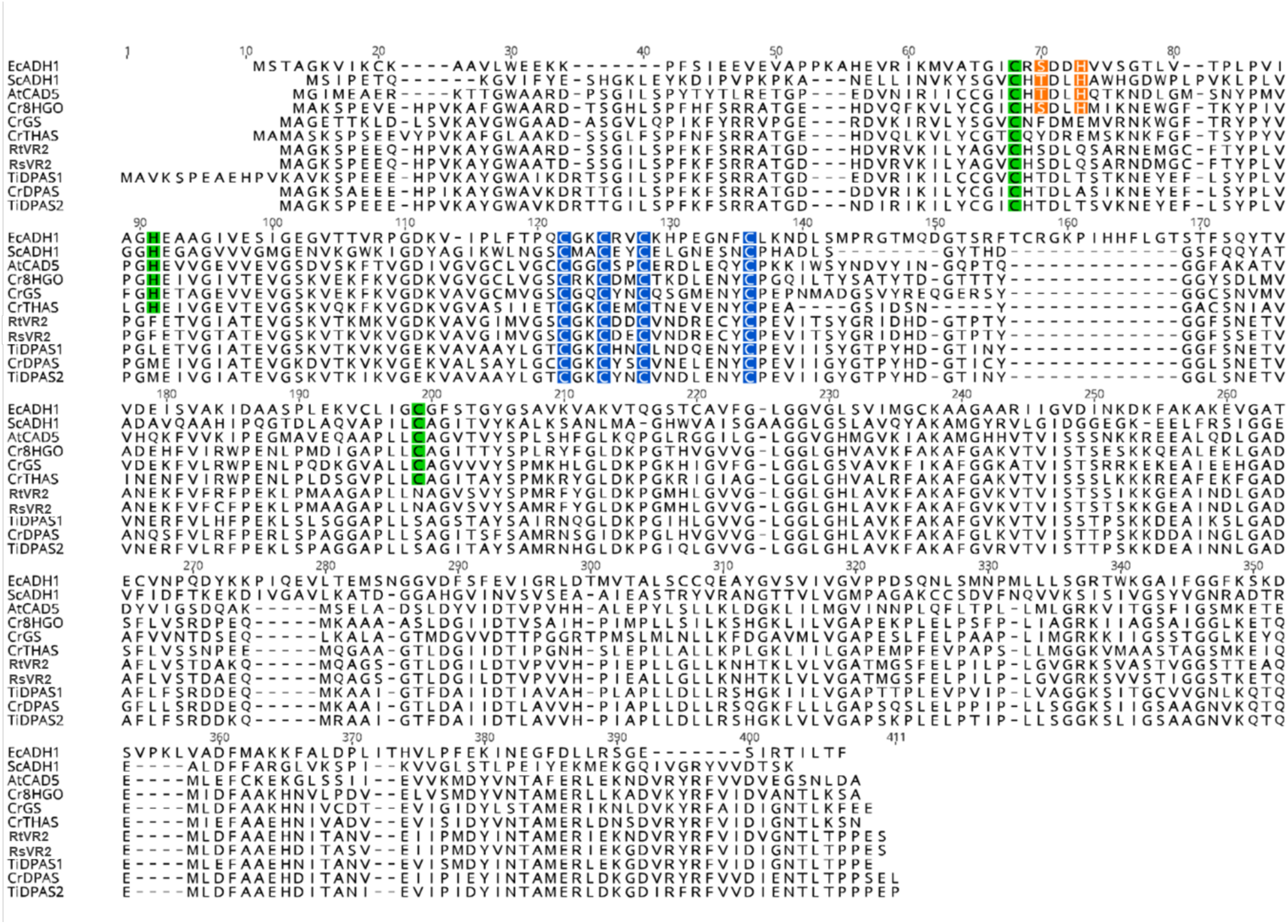
MUSCLE amino acid sequence alignment of ADHs highlighting key residues. Catalytic zinc coordinating residues are labelled in green, structural zinc coordinating residues in blue and proton relay residues in orange. Protein names and uniprot accessions: *Equus caballus* alcohol dehydrogenase (*Ec*ADH1) P00327; *Saccharomyces cerevisiae* alcohol dehydrogenase 1 (*Sc*ADH1), P00330; *Arabidopsis thaliana* cinnamyl alcohol dehydrogenase 5 (*At*CAD5), O49482; *Catharanthus roseus* 8-hydroxygeraniol dehydrogenase (*Cr*8HGO), Q6V4H0; *Catharanthus roseus* geissoschizine synthase (*Cr*GS), W8JWW7; *Catharanthus roseus* tetrahydroalstonine synthase (*Cr*THAS), A0A0F6SD02; *Rauwolfia tetraphylla* vomilenine reductase 2 (*Rt*VR2) A0A0U4BHM2, *Rauwolfia serpentina* vomilenine reductase 2 (*Rs*VR2), A0A0U3S9Q3; *Tabernanthe iboga* dihydroprecondylocarpine acetate synthase 1 (*Ti*DPAS1), A0A5B8XAH0; *Catharanthus roseus* dihydroprecondylocarpine acetate synthase (*Cr*DPAS), A0A1B1FHP3; *Tabernanthe iboga* dihydroprecondylocarpine acetate synthase 2 (*Ti*DPAS2), A0A5B8X8Z0.

**Figure S2.**
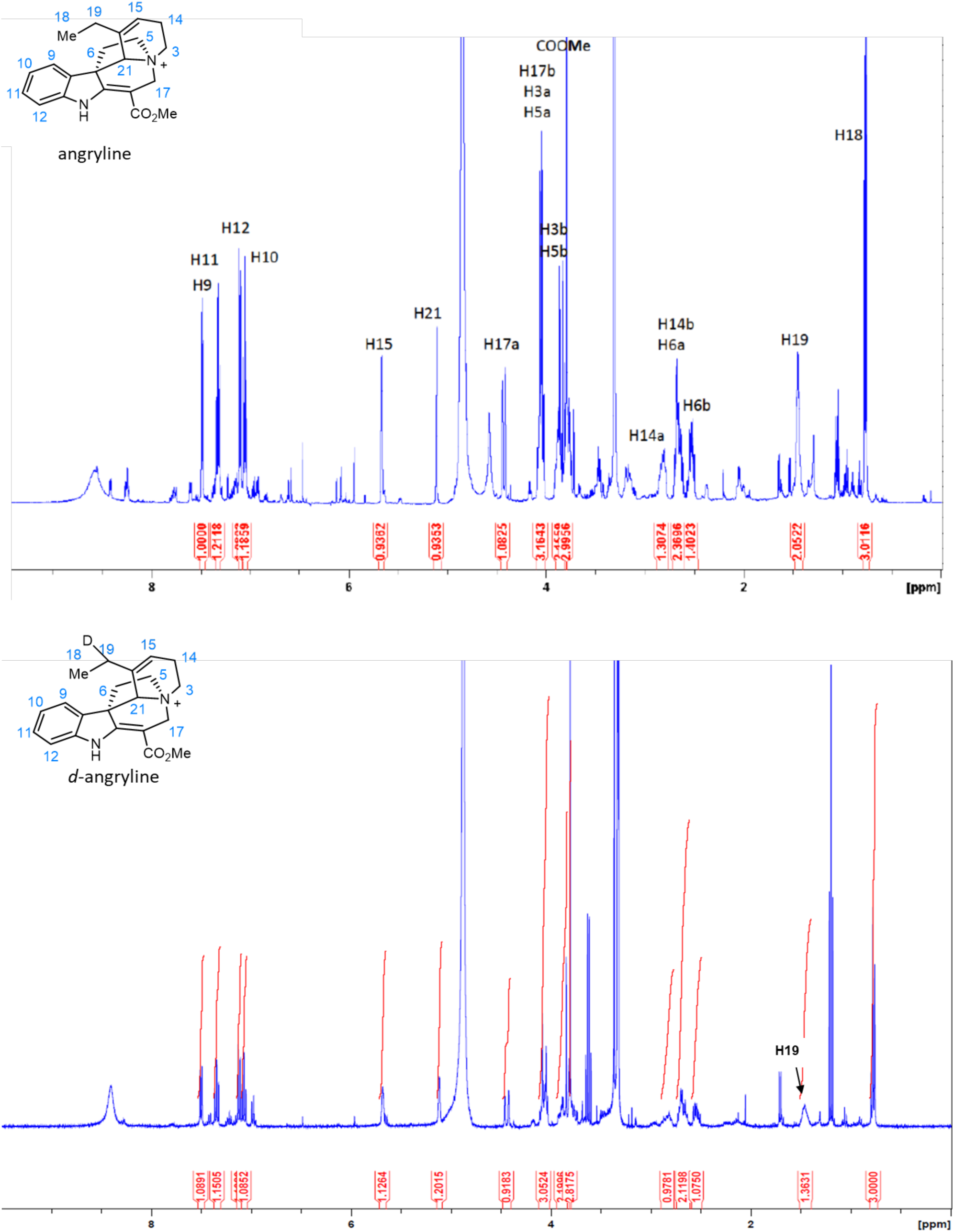
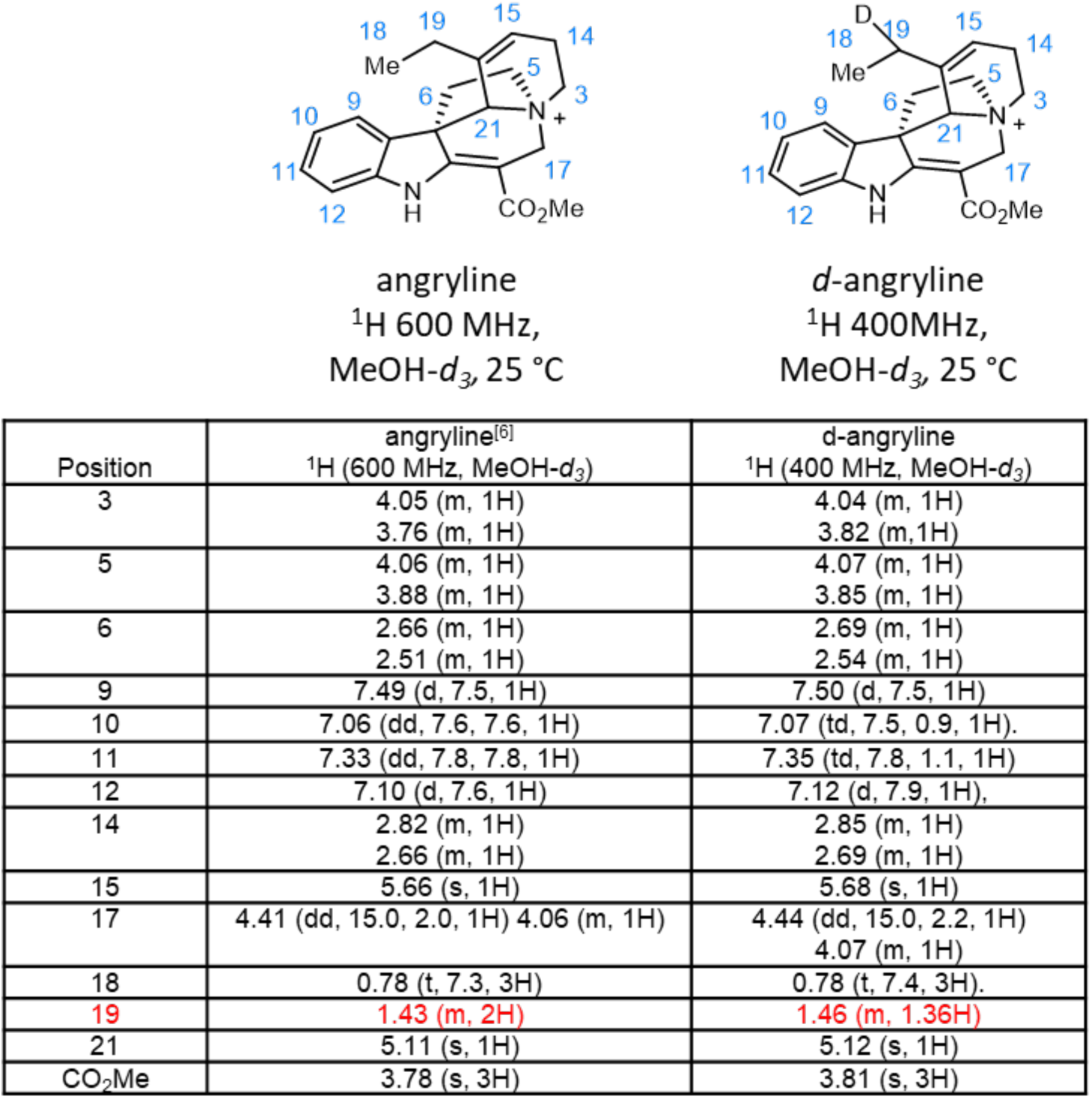
Comparison of ^1^H NMR data for angryline and *d*-angryline showing position of deuterium incorporation. Angryline was characterised by NMR in a previous study ^[6]^.

**Figure S3.**
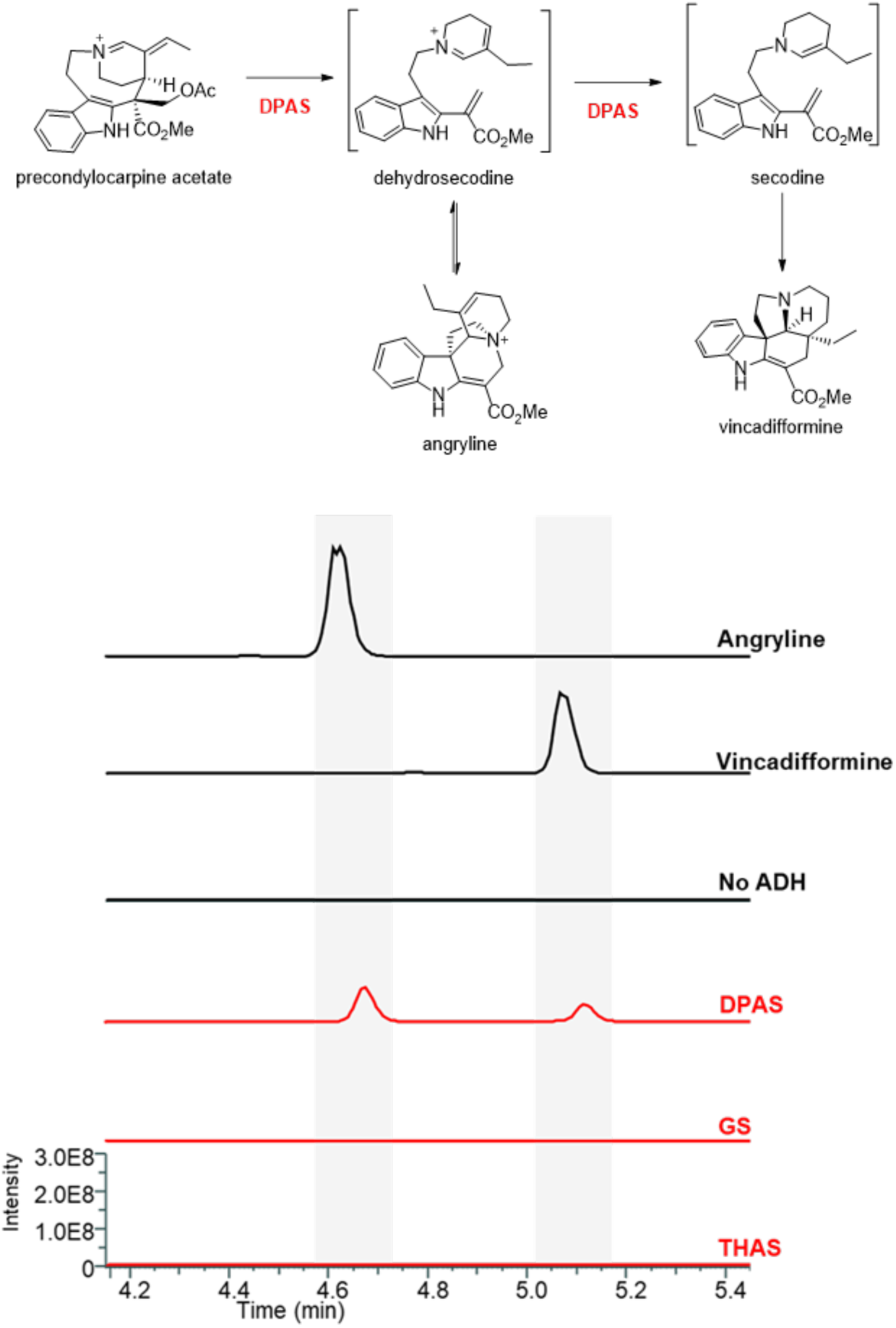
LC-MS chromatograms showing formation of angryline and vincadifformine in absence of cyclase with ADHs and NADPH. EIC *mz* 337.00-341.00.

**Figure S4.**
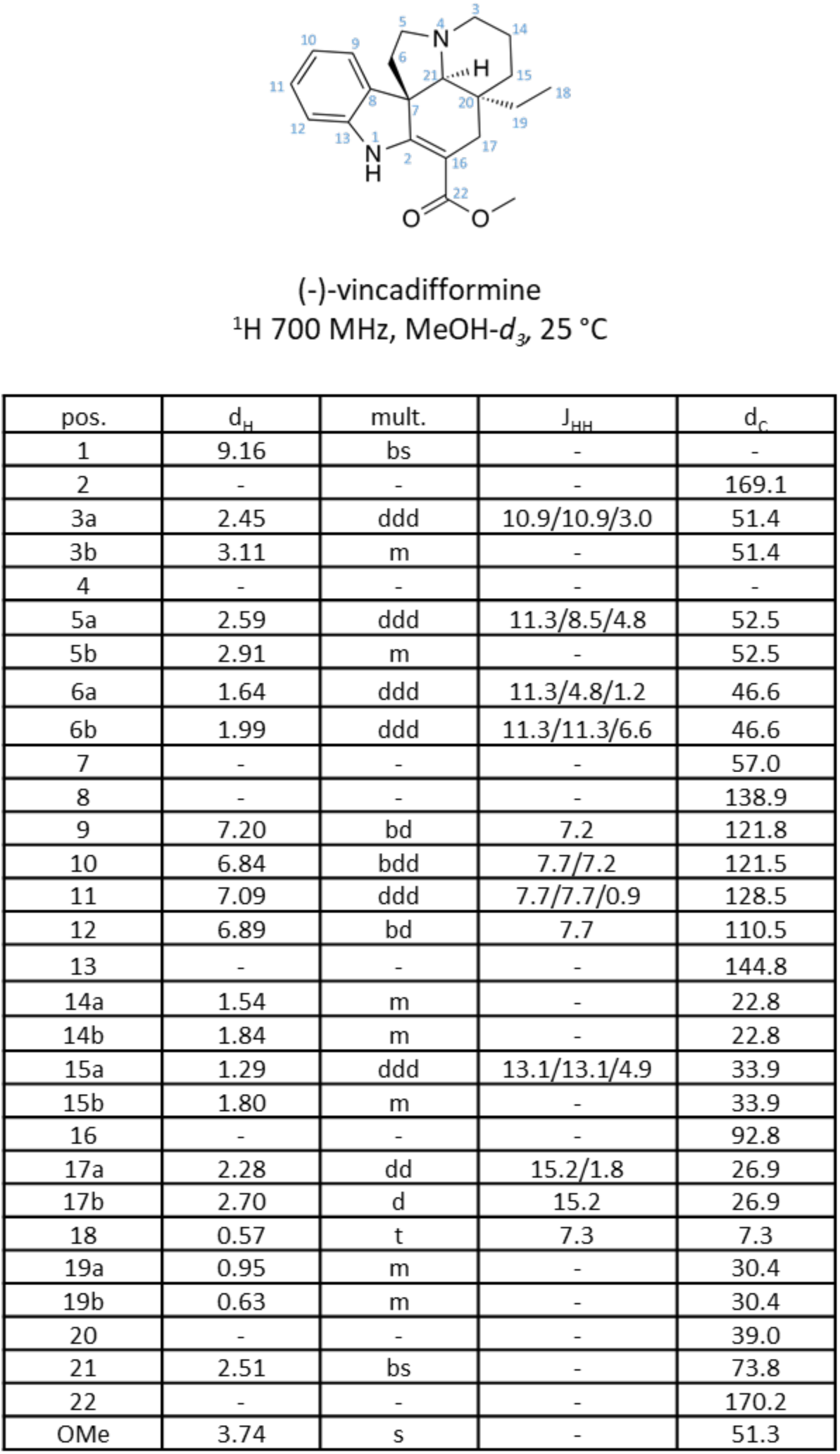
^1^H NMR data for (-)-vincadifformine in MeOH-*d_3_*.

**Figure S5.**
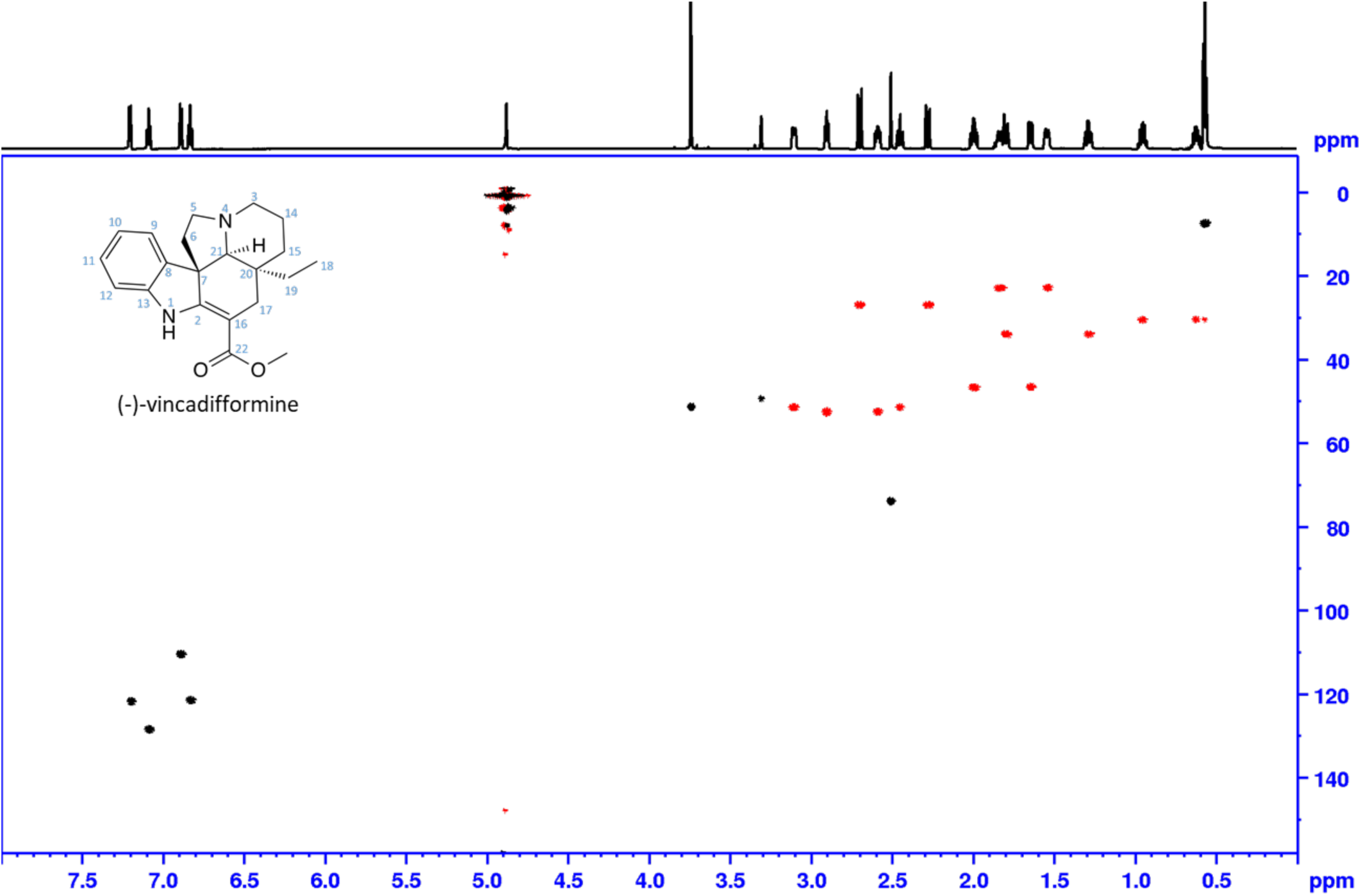
^1^H NMR data for *m/z* 339, (–)-vincadifformine (standard). Phase sensitive HSQC, full range in MeOH-*d_3_*. Shaded areas mark impurity and solvent, red: CH2, black: CH, CH3. NMR data of (–)-vincadifformine in chloroform-*d* has been previously reported^[25,26]^.

**Figure S6.**
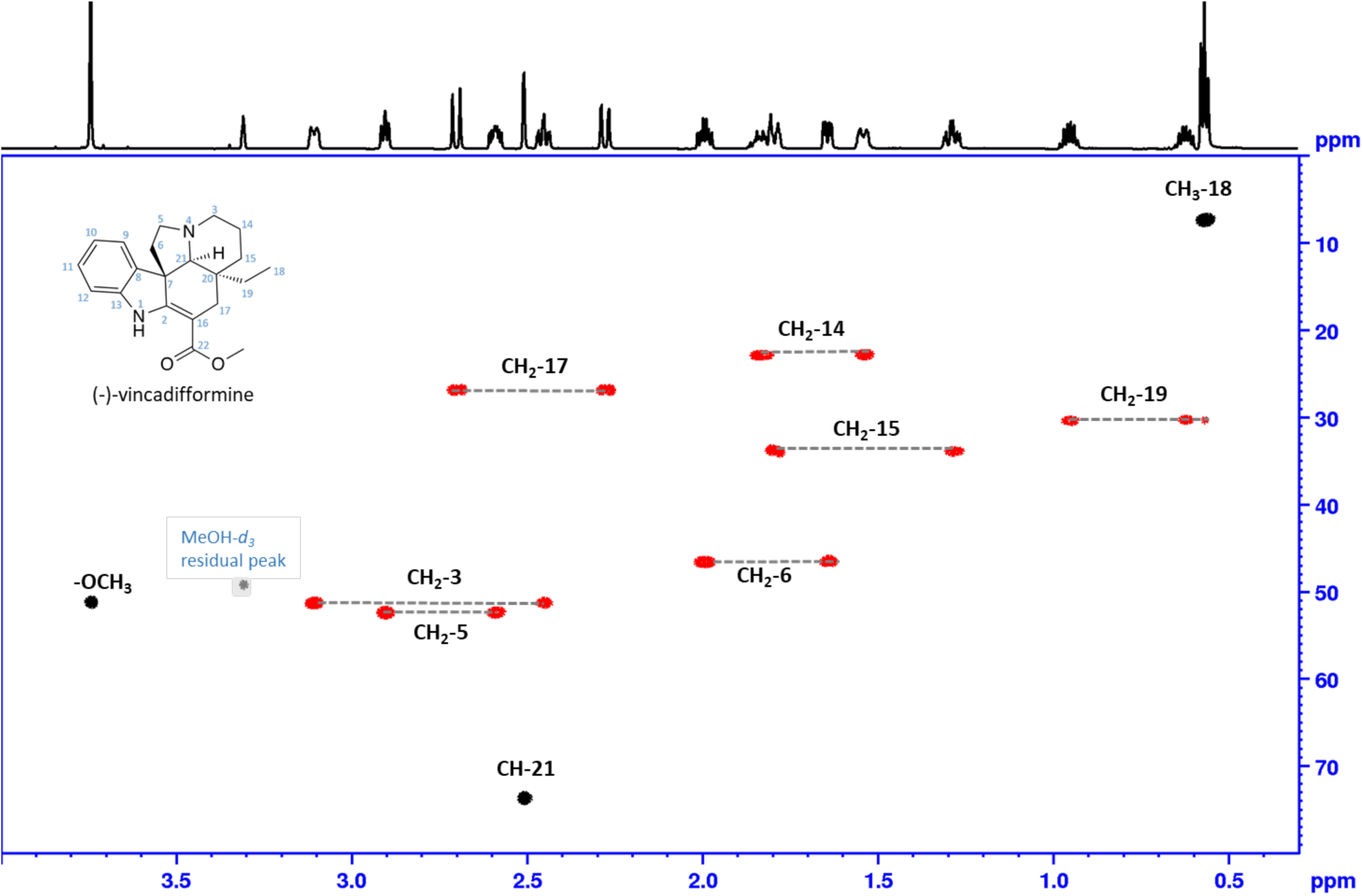
^1^H NMR data for *m/z* 339, (–)-vincadifformine (standard). Phase sensitive HSQC, aliphatic range in MeOH-d3. Shaded areas mark impurity and solvent, red: CH2, black: CH, CH3

**Figure S7.**
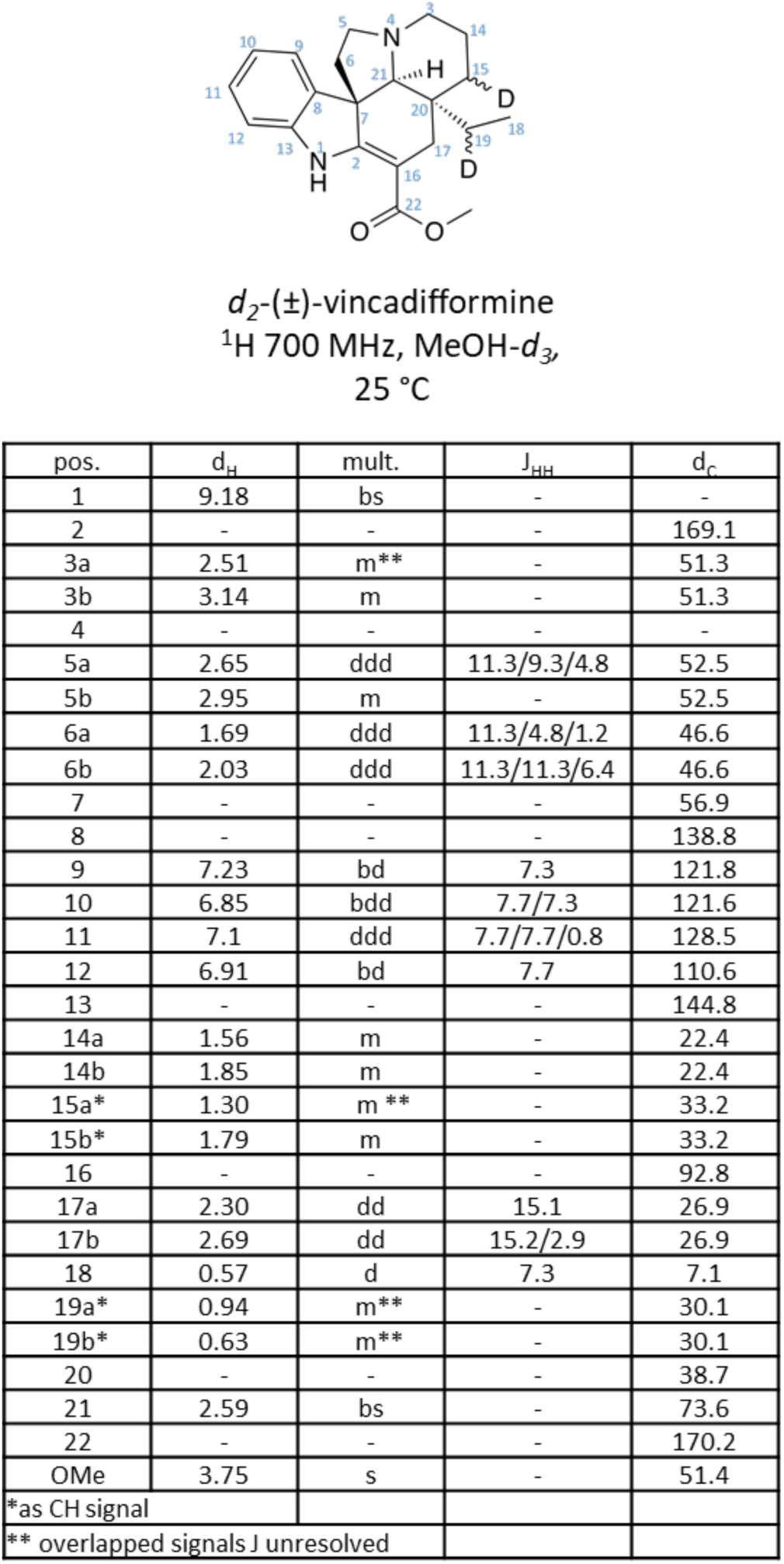
^1^H NMR data for *d_2_*-(±)-vincadifformine in MeOH-*d_3_*.

**Figure S8.**
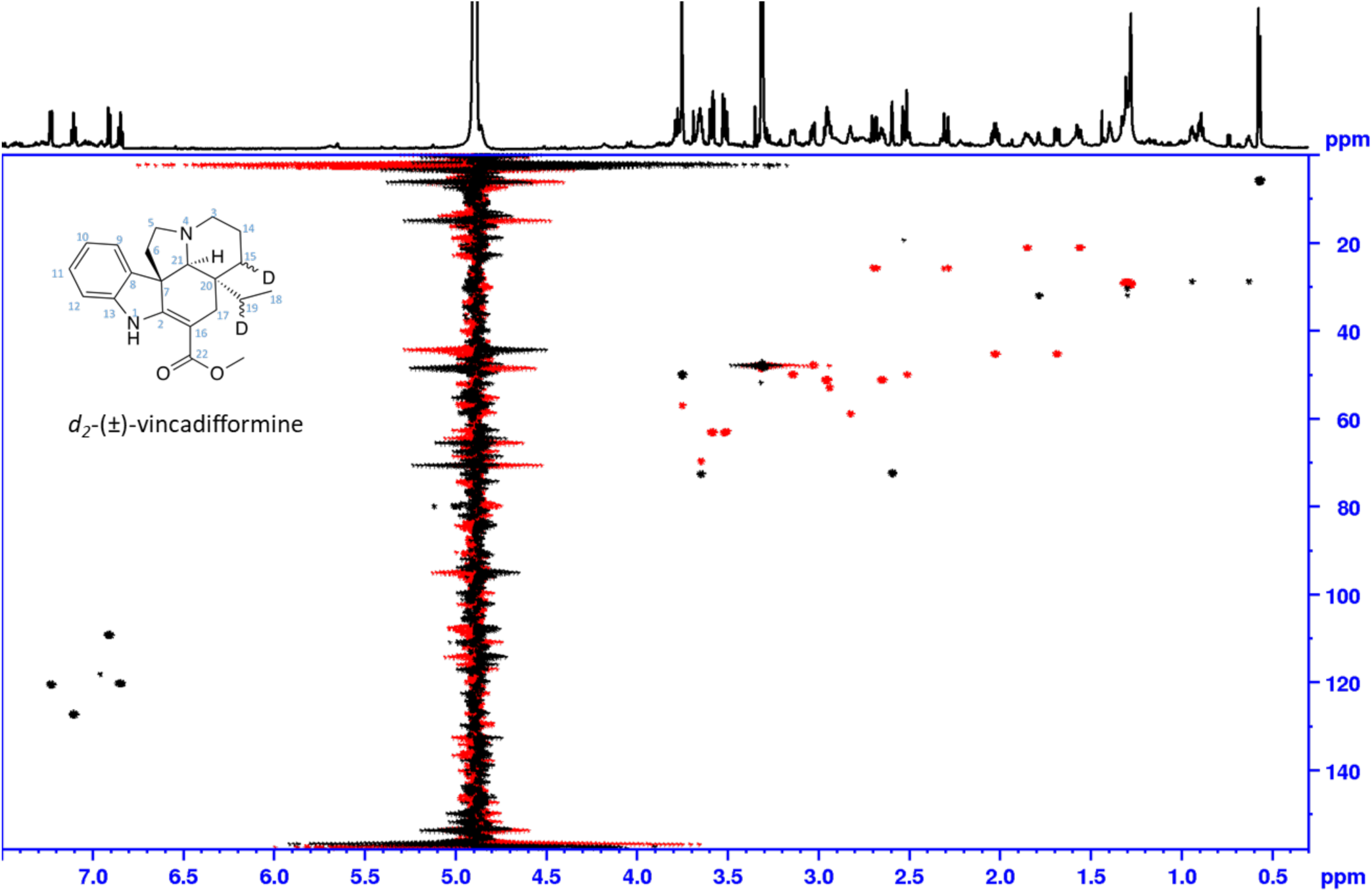
Phase sensitive HSQC NMR data for *m/z* 341, *d_2_*-(±)-vincadifformine full range in MeOH-*d_3_*. Shaded areas mark impurity and solvent, red: CH2, black: CH, CH3

**Figure S9.**
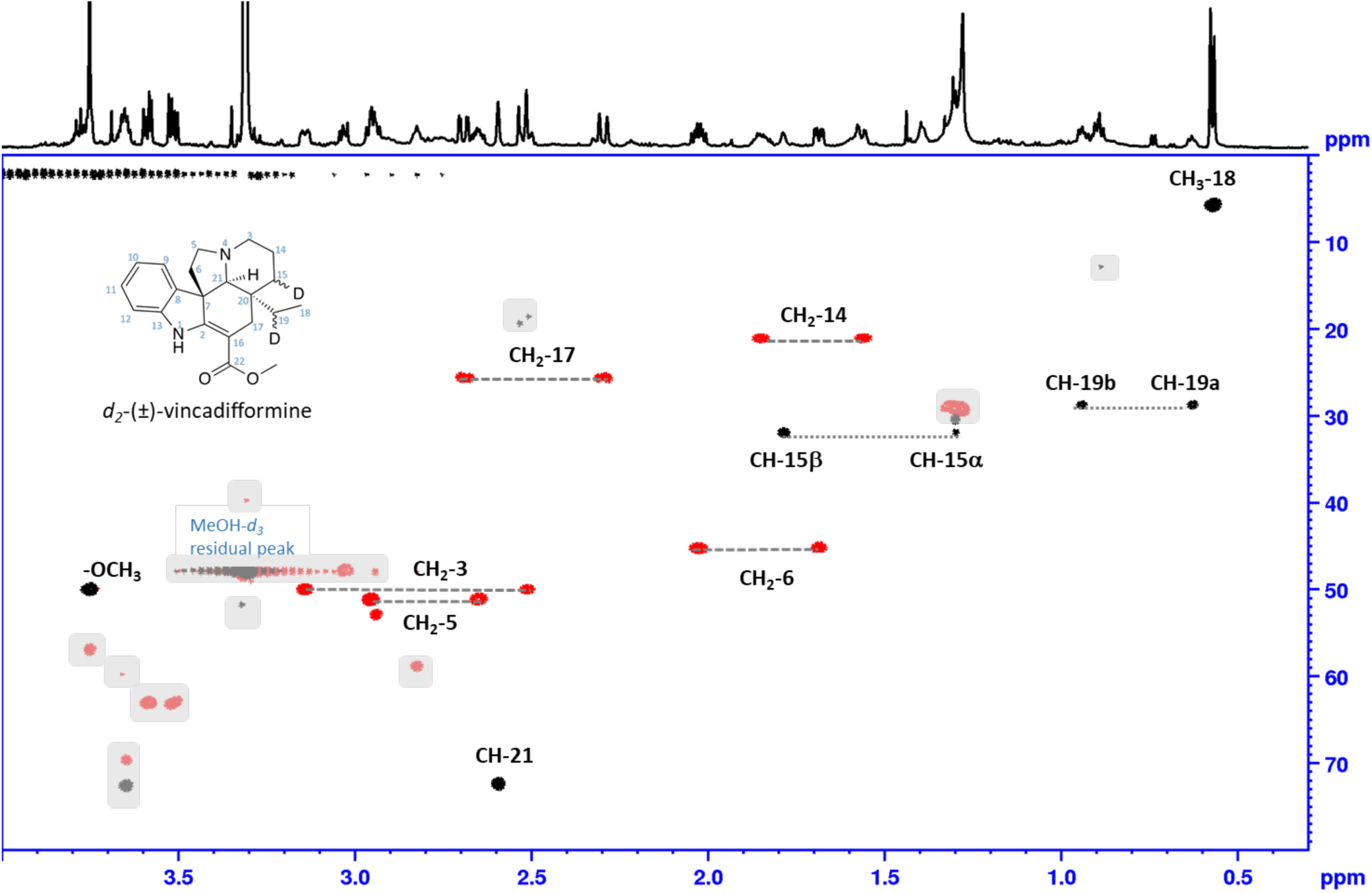
Phase sensitive HSQC NMR data for *m/z* 341, *d_2_*-(±)-vincadifformine, aliphatic range in MeOH-*d_3_*. Shaded areas mark impurity and solvent, red: CH2, black: CH, CH3

**Figure S10.**
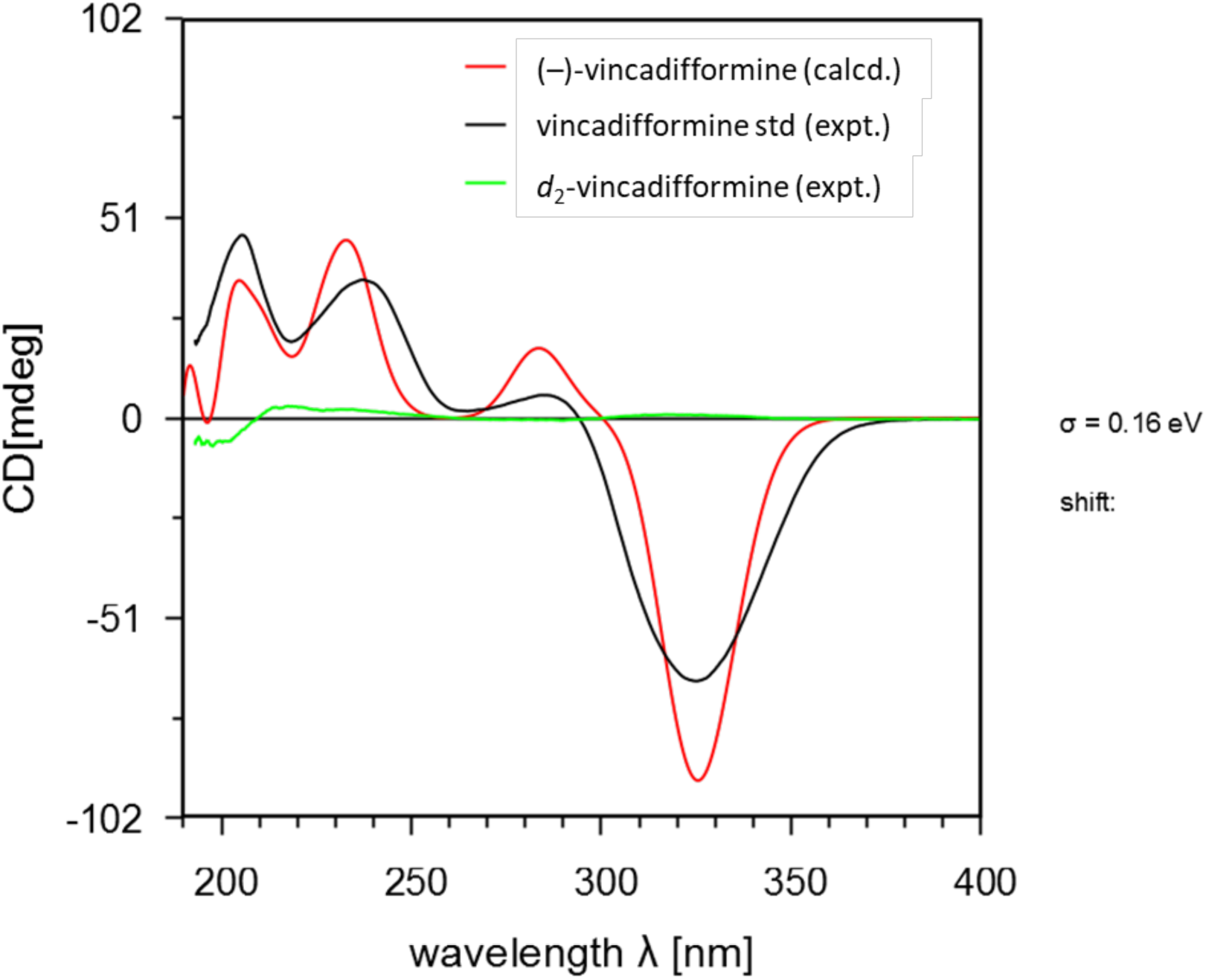
Comparison of the experimental ECD spectra of *d_2_*-vincadifformine (green), vincadifformine standard (black) and calculated ECD spectra of (-)-vincadifformine (red).

**Figure S11.**
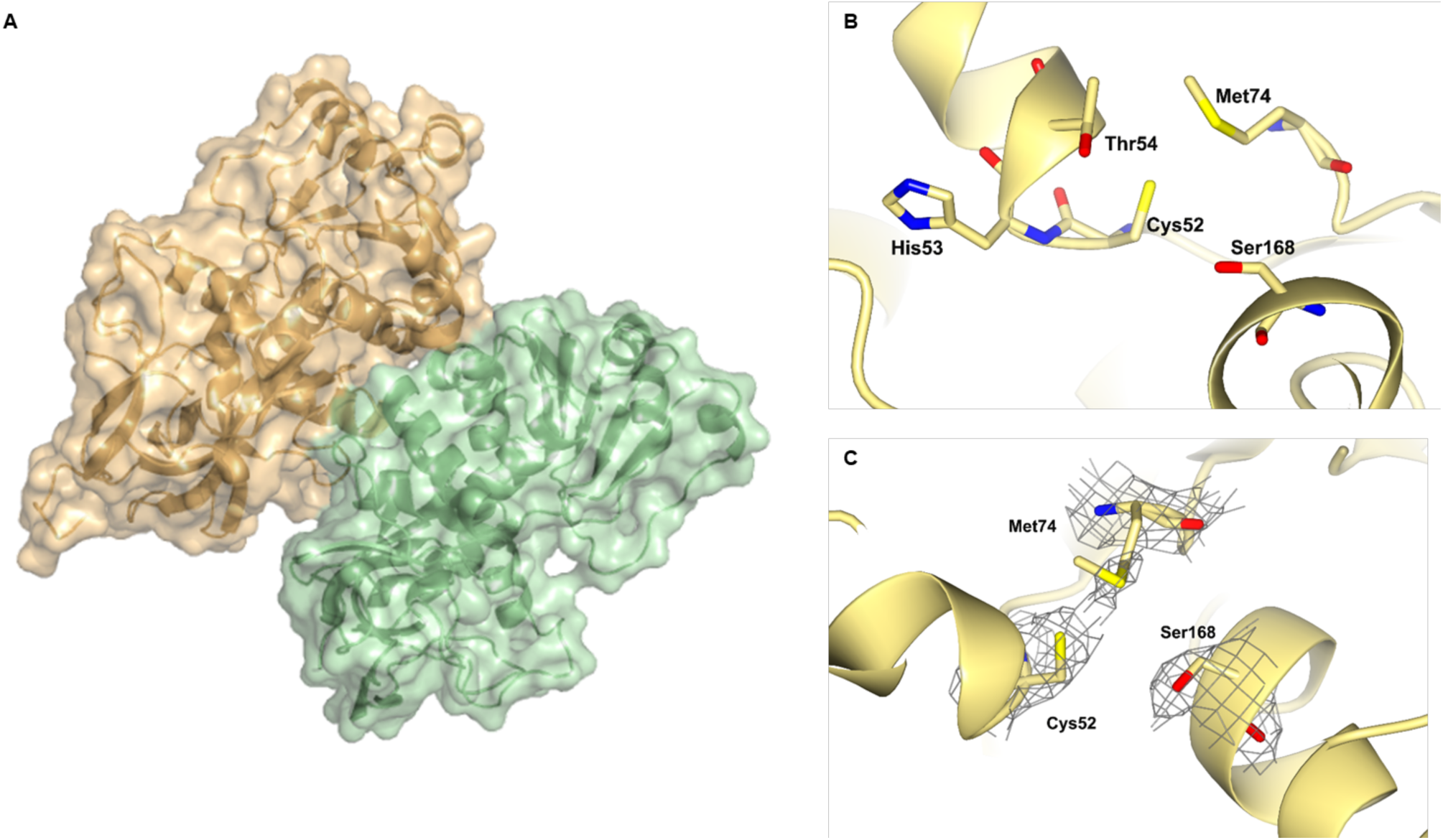
Structure of *Catharanthus roseus* DPAS (*Cr*DPAS) crystallised as a homodimer at 2.45 Å resolution. **A.** Structure coloured by chains. Structure lacked electron density for residues Gly102-Thr134. **B.** Active site of *Cr*DPAS showing atypical residues canonically involved in coordination of the catalytic zinc. **C.** Electron density of residues of *Cr*DPAS canonically involved in coordinating the catalytic zinc.

**Figure S12.**
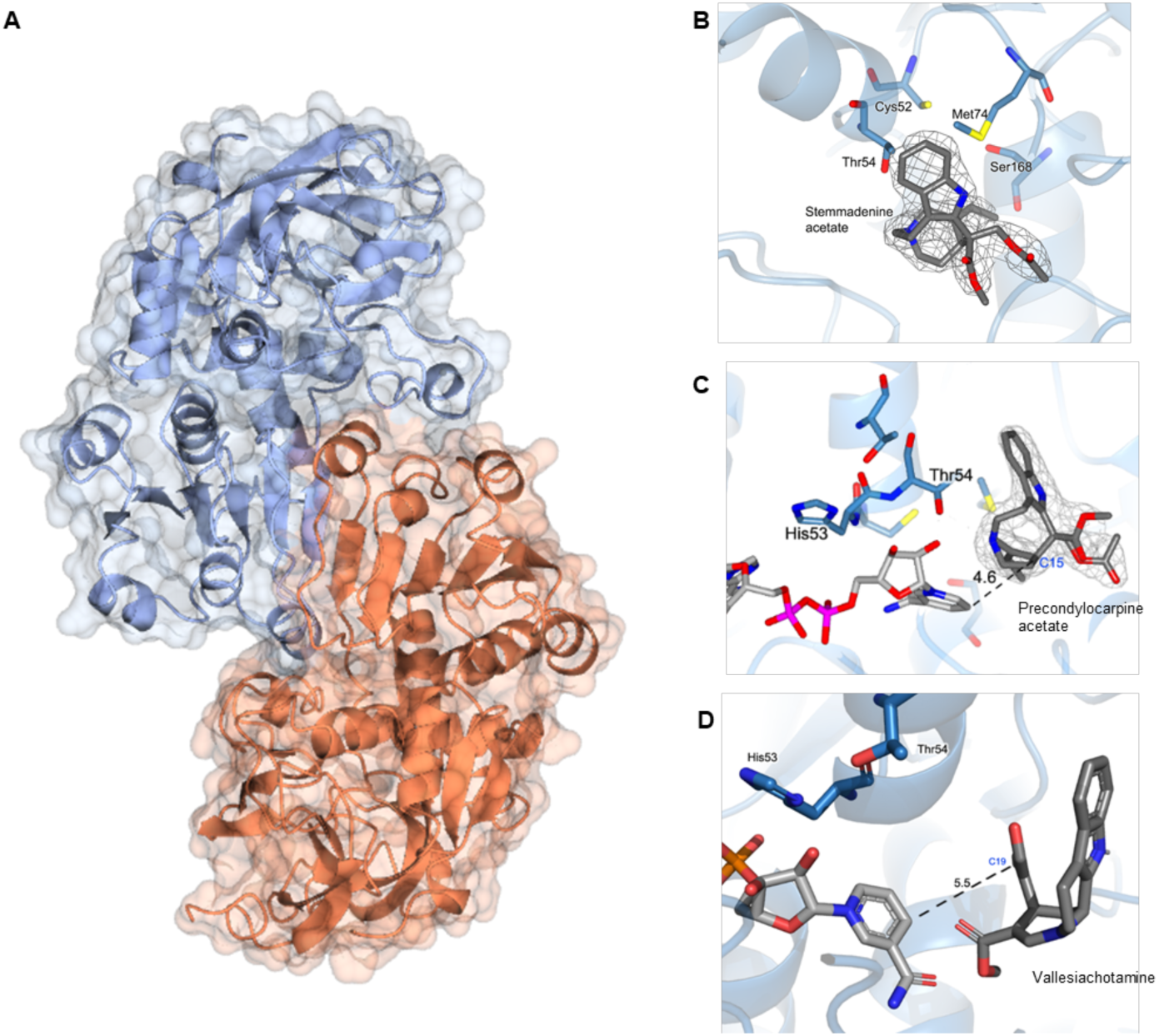
Structure of *Tabernanthe iboga* DPAS2 (*Ti*DPAS2). **A.** Apo-*Ti*DPAS2 crystallised as a homodimer at 2.42 Å coloured by chains. Electron density for NADP^+^ cofactor was not observed. **B.** Active site of *Ti*DPAS2 bound to stemmadenine acetate. **C.** *Ti*DPAS2 bound to precondylocarpine acetate and NADPH cofactor showing distance between the 4-pro-*S*-hydride of NADPH and position of reduction. **D.** *Ti*DPAS2 modelled with vallesiachotamine and NADPH cofactor showing distance between the 4-pro-*S*-hydride of NADPH and position of reduction.

**Figure S13.**
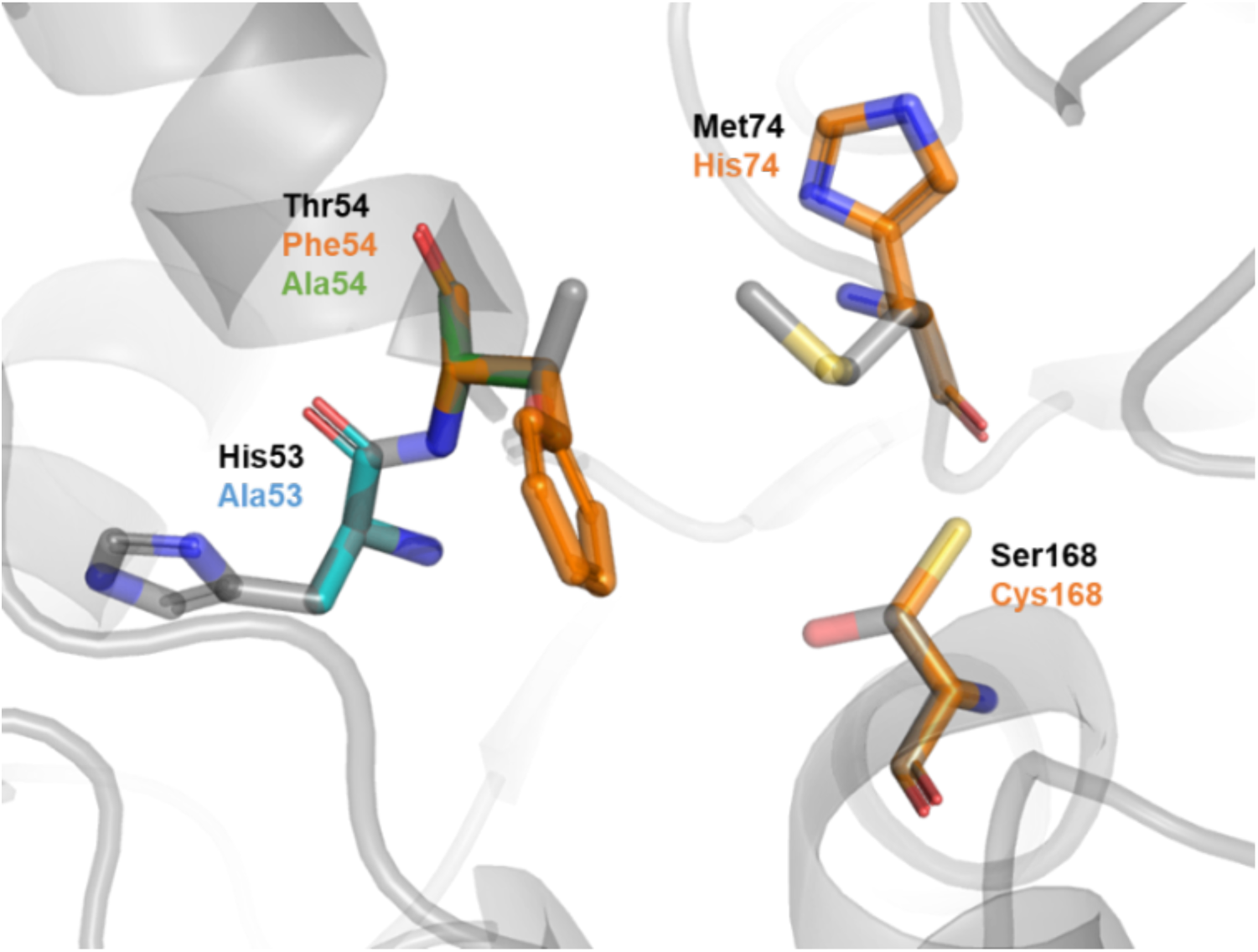
Catharanthus DPAS (*Cr*DPAS) active site with highly conserved residues involved in canonical ADH enzymes with the coordination of the catalytic zinc (Met74 Ser168) and the proton relay (His53, Thr54) that were mutated in this study.

**Figure S14.**
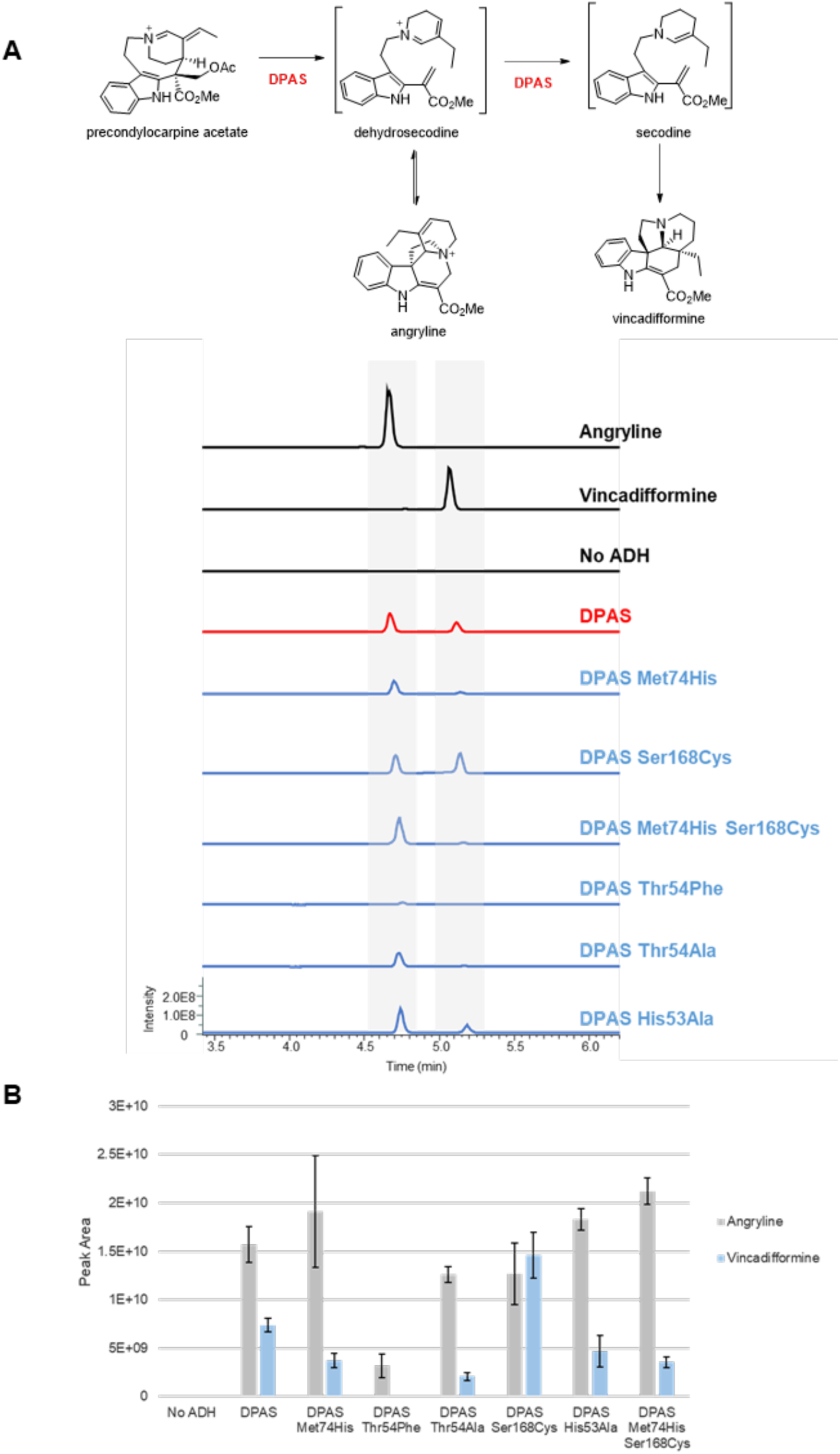
Wild-type *Cr*DPAS and mutants incubated with precondylocarpine acetate showing production of angryline and vincadifformine. **A.** Representative LC-MS chromatogram for wild-type DPAS and mutants. EIC *m/z* 337.05-340.05 **B.** Peak area of angryline and vincadifformine products of DPAS and mutants resulting from an endpoint assay. n=3, bars represent standard deviation.

**Figure S15.**
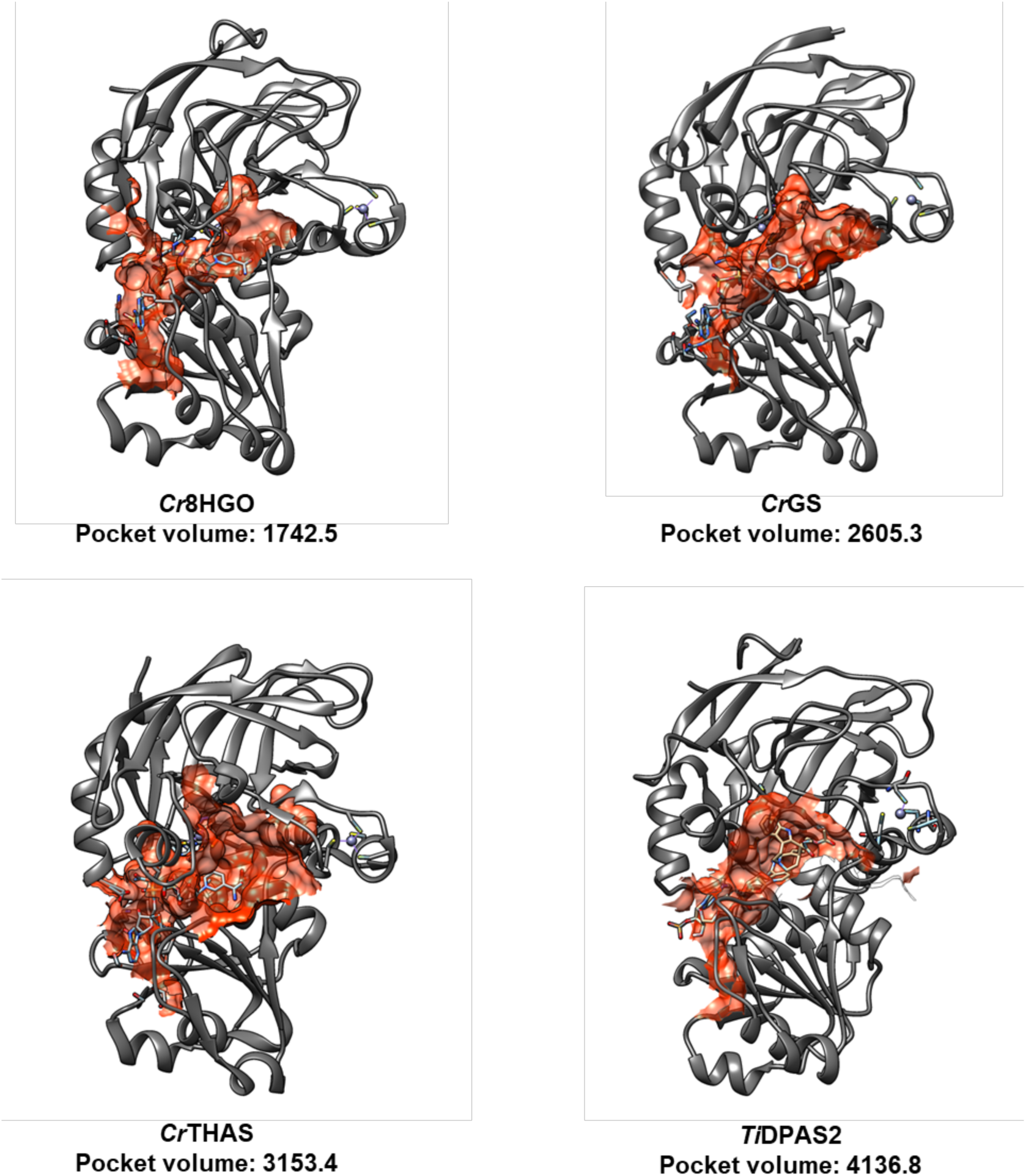
Substrate cavity volume and area of *Cr*8HGO, *Cr*GS, *Cr*THAS and *Ti*DPAS2. Cavity coloured in red, in co-crystallised (8HGO, GS and THAS) or modelled (DPAS2) cofactor NADP^+^ coloured in grey, and bound precondylocarpine acetate in DPAS2 coloured in yellow. Pocket volumes computed by CASTp 3.0 ^[21]^.

**Figure S16.**
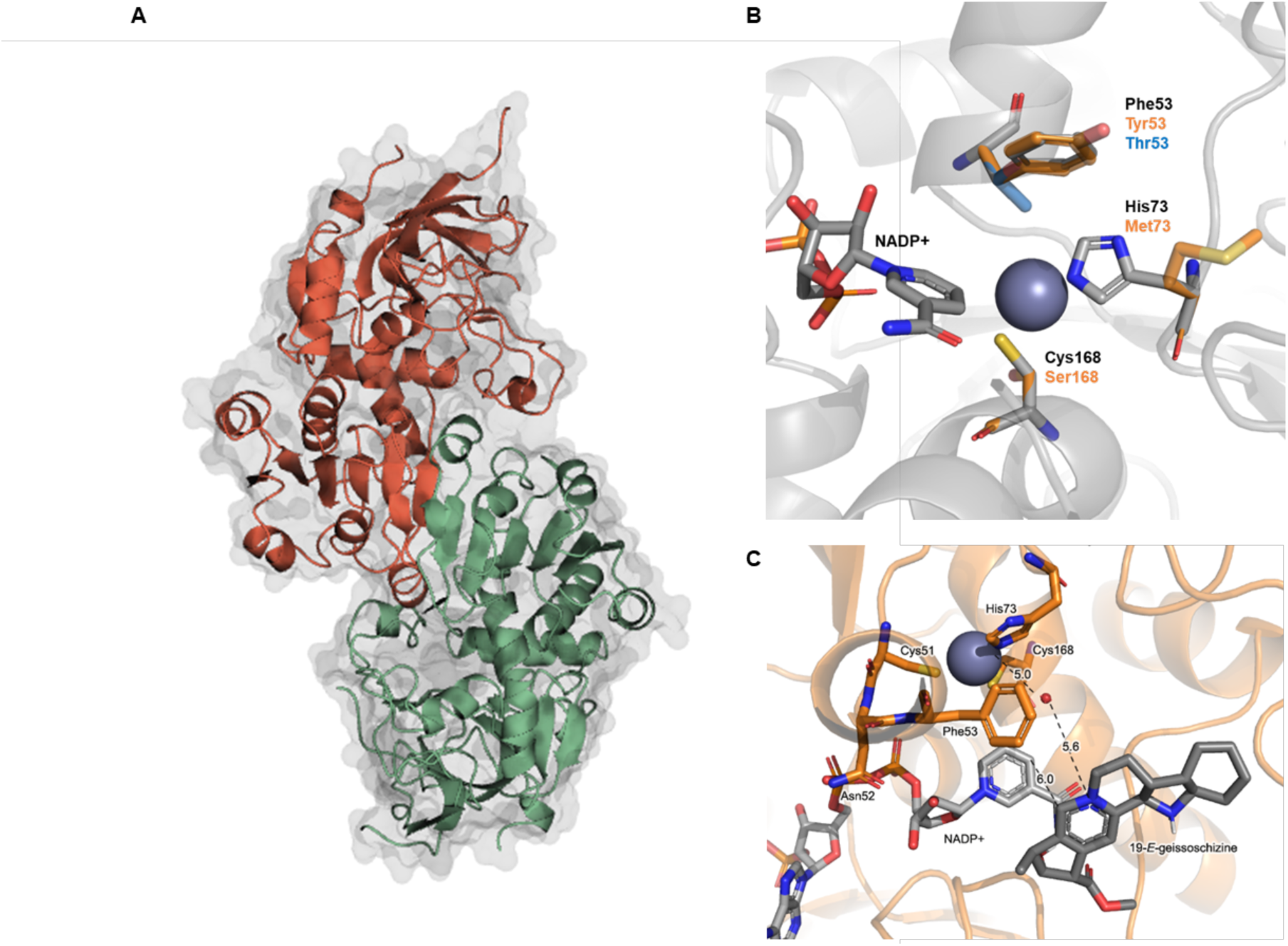
**A.** *Catharanthus roseus* GS (*Cr*GS) crystallised as a homodimer bound to cofactor NADP^+^ at 2.00 Å. Structure coloured by chains. **B.** GS active site with residues involved in coordination of the catalytic zinc and proton relay mutated in this study. **C.** NADP^+^-bound GS active site docked with 19-*E-*geissoschizine. Distance between the 4-pro-*S*-hydride of NADPH and position of reduction, and the distance between the catalytic zinc, bound water molecule and the substrate iminium as in proposed enzyme mechanism are shown.

**Figure S17.**
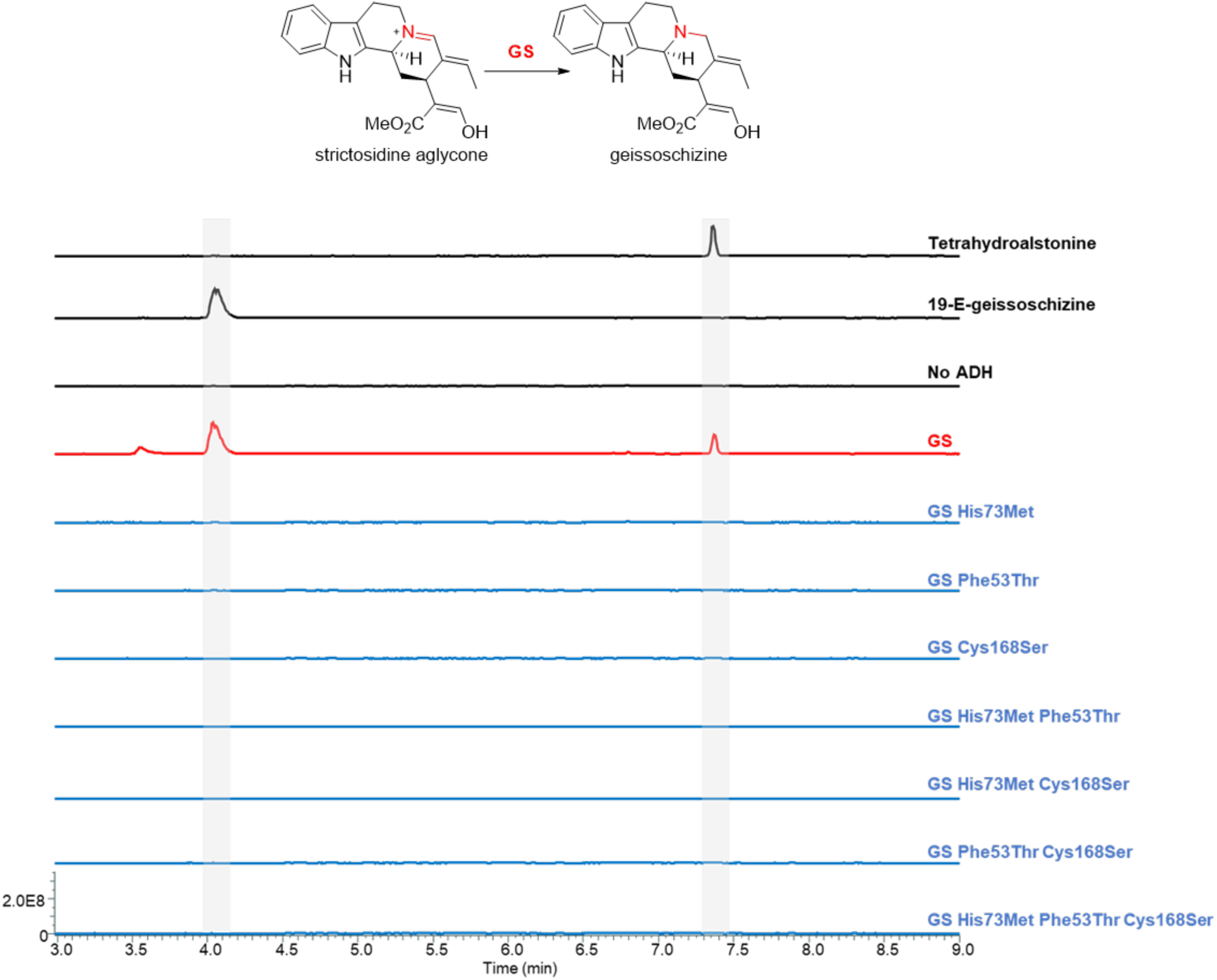
LC-MS chromatograms of *Cr*GS and mutants reacted with strictosidine aglycone. These mutants probe the role of residues involved in coordination of the catalytic zinc and involved in the proton relay. EIC *m/z* 353.185-353.225.

**Figure S18.**
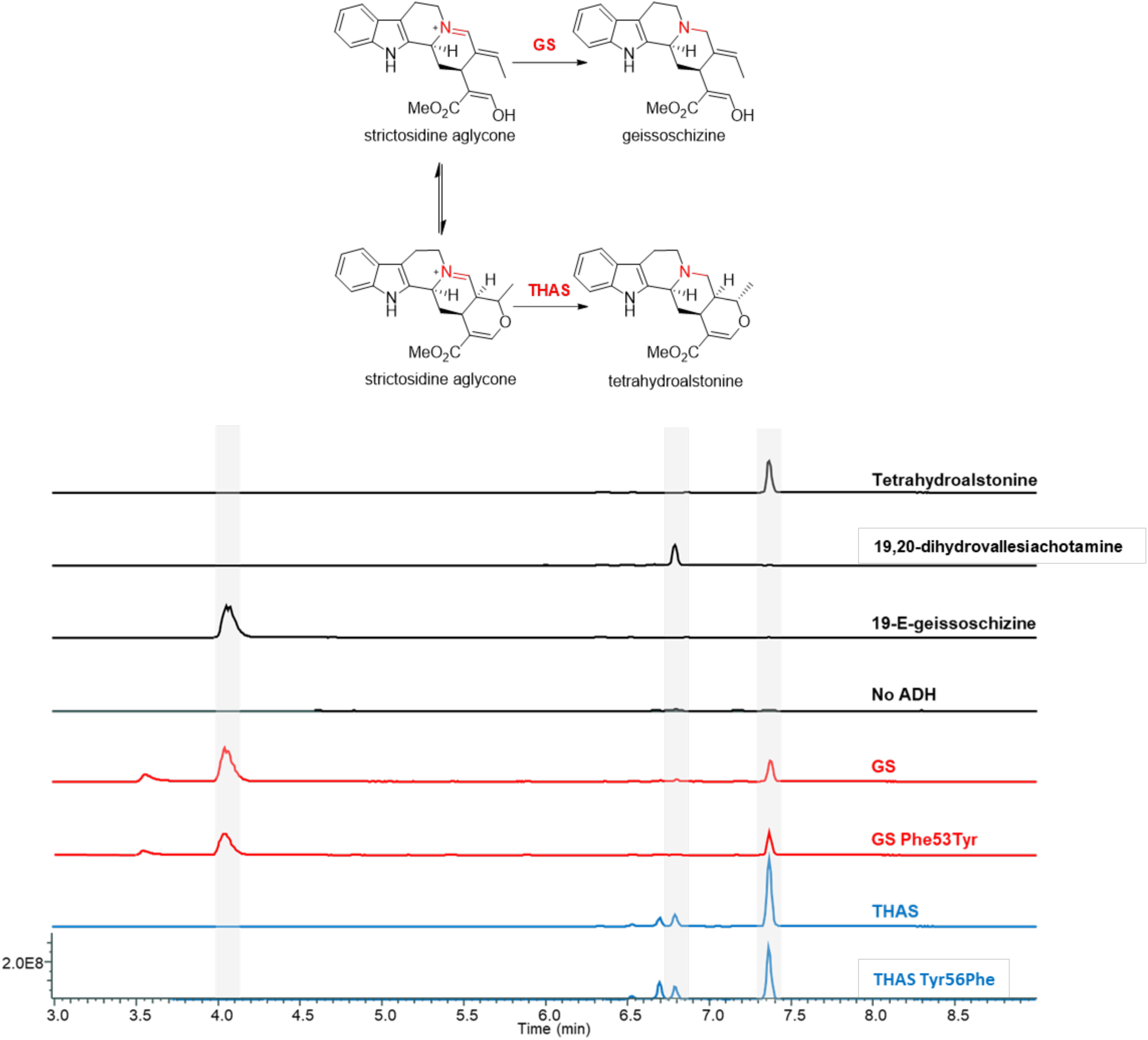
LC-MS chromatograms of GS and GS Phe53Tyr mutant and the corresponding THAS and THAS Tyr56Phe mutant reacted with strictosidine aglycone. These mutations probe the role of the hydroxyl group in the proton relay. EIC *m/z* 353.185-353.225.

**Figure S19.**
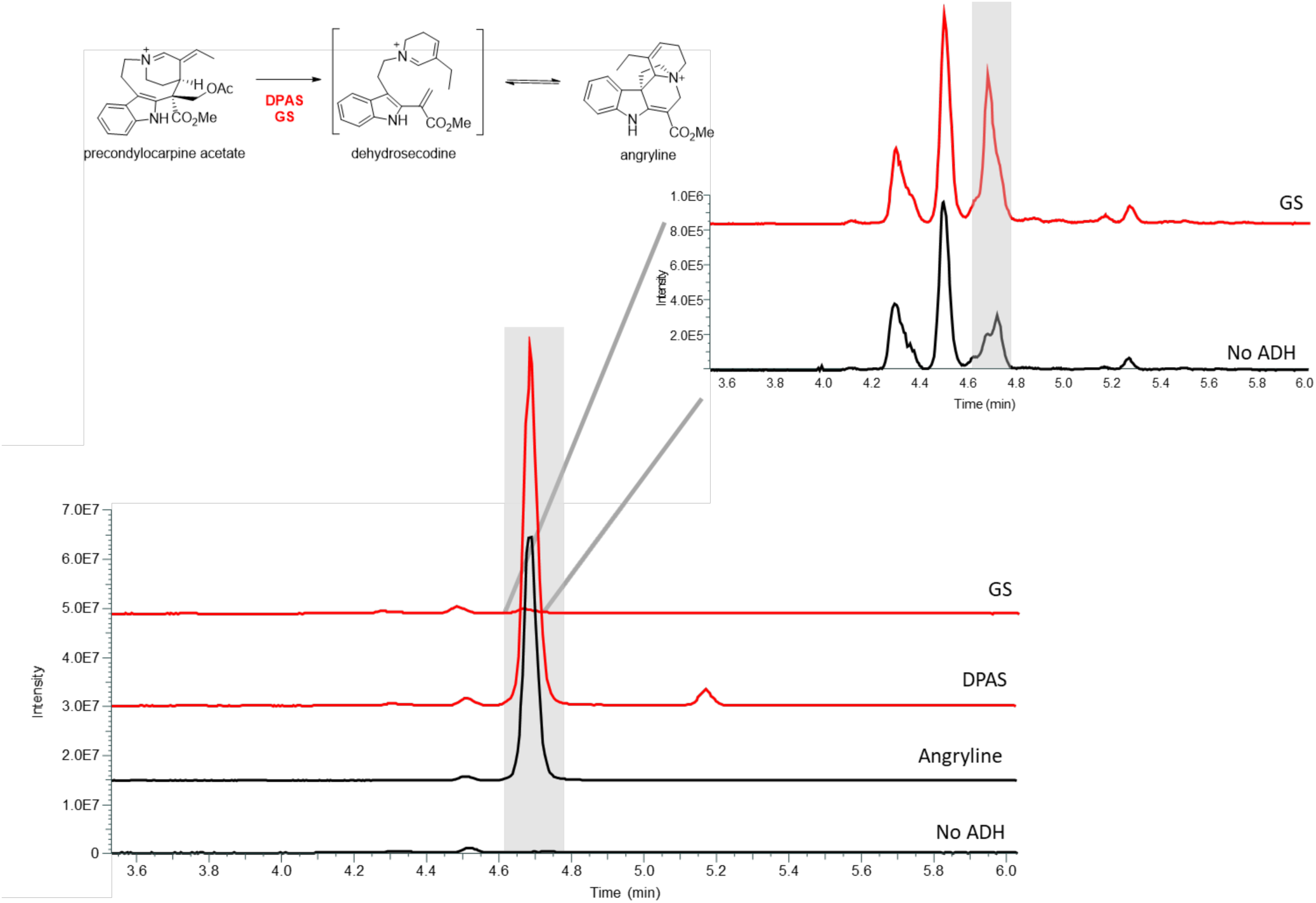
LC-MS chromatograms of *Cr*DPAS and CrGS reacted with precondylocarpine acetate incubated for 1 hour at 30 °C. EIC *m/z* 337.180-337.200. Inset of GS reaction to show small amount of peak with same mass and elution time as angryline standard.

**Figure S20.**
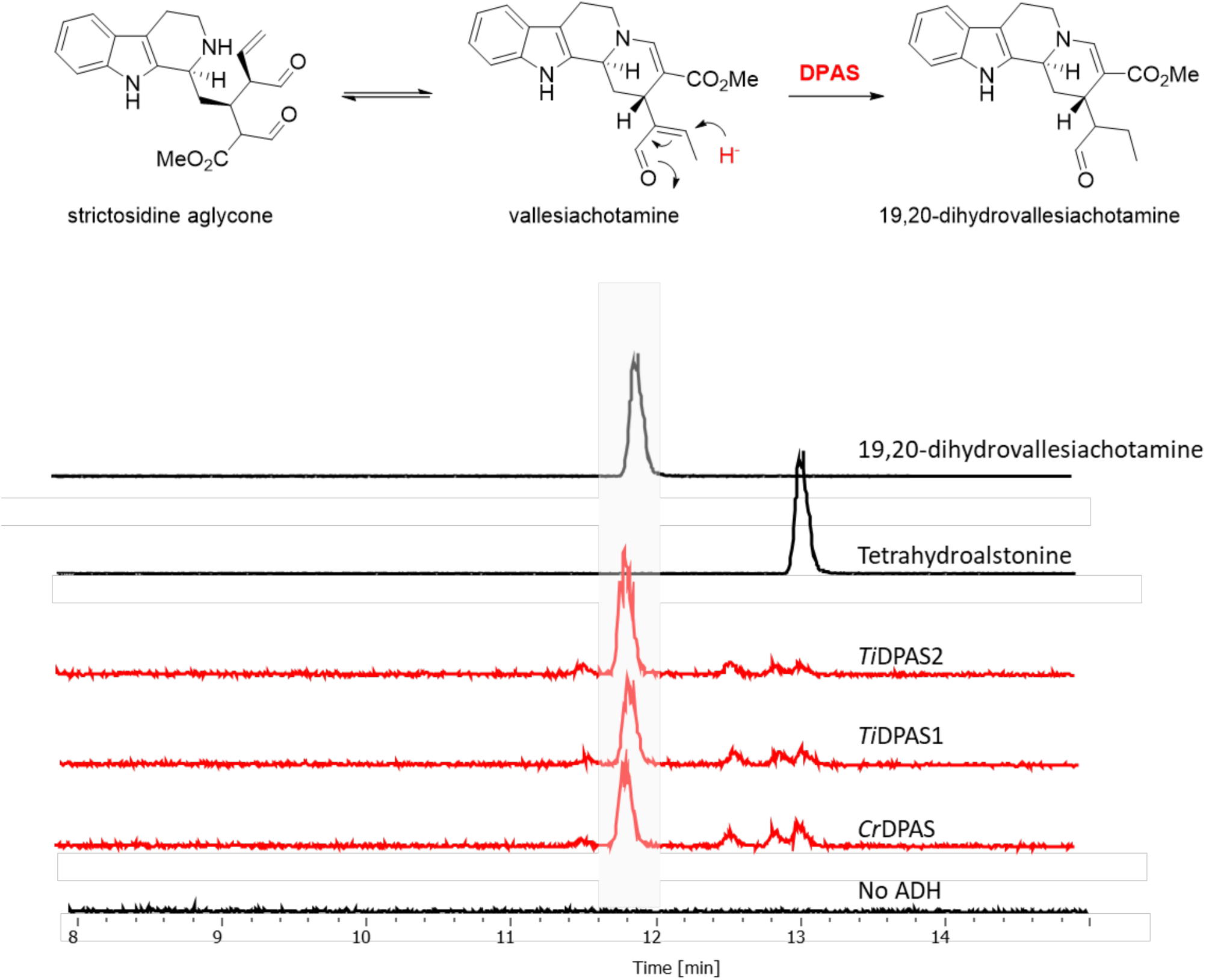
LC-MS chromatograms of *Cr*DPAS, *Ti*DPAS1 and *Ti*DPAS2 reacted with strictosidine aglycone. EIC *m/z* 353.185-353.225.

**Figure S21.**
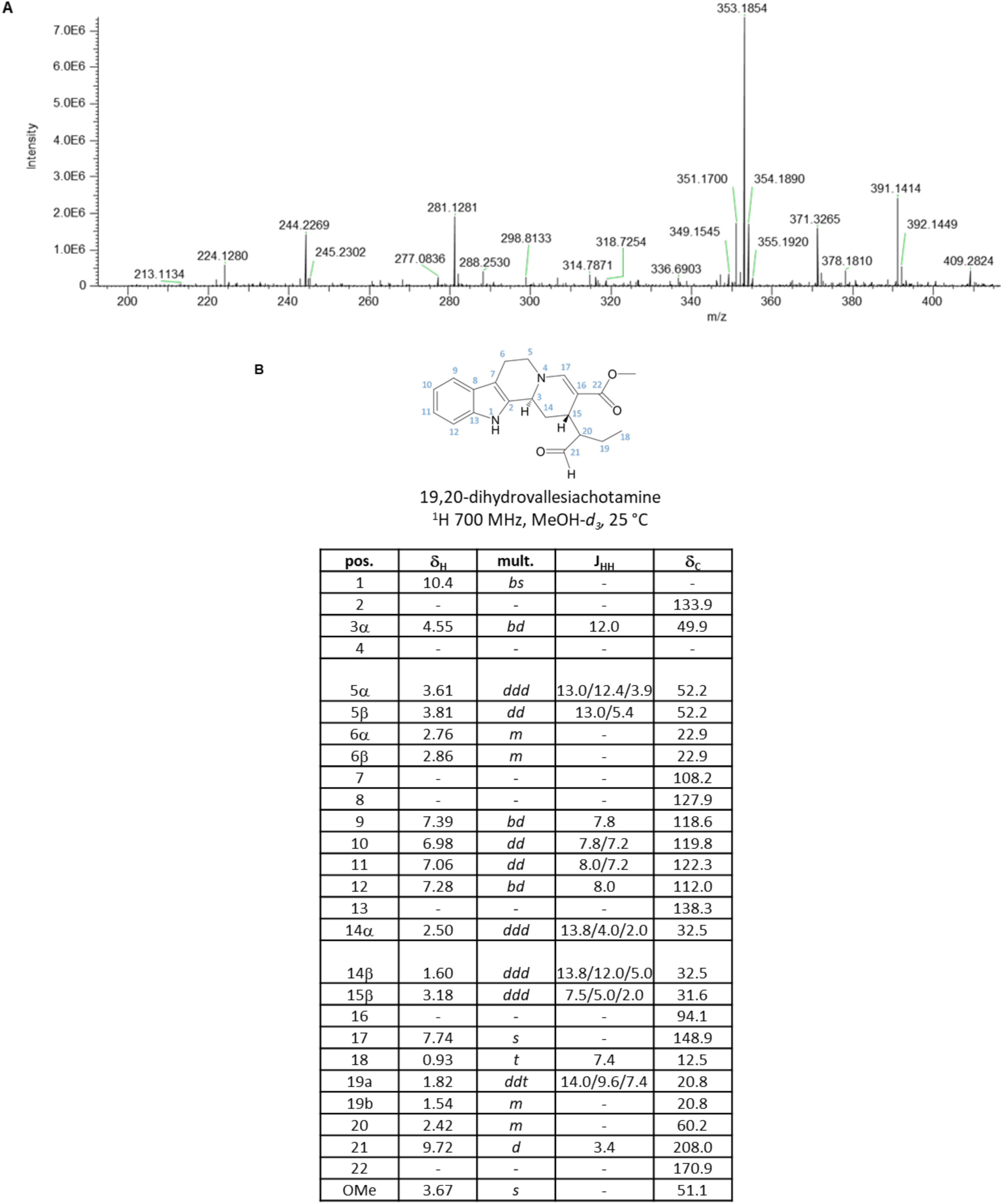
MS/MS and NMR data of 19,20-dihydrovallesiachotamine. **A.** MS/MS spectra of 19,20-dehydrovallesiachotamine. Formula: C_21_H_24_N_2_O_3_; observed mass: 353.1854; theoretical mass: 353.1860; error 1.6988 p.p.m. **B.** ^1^H NMR spectra for 19,20-dihydrovallesiachotamine in MeOH-*d_3_*.

**Figure S22.**
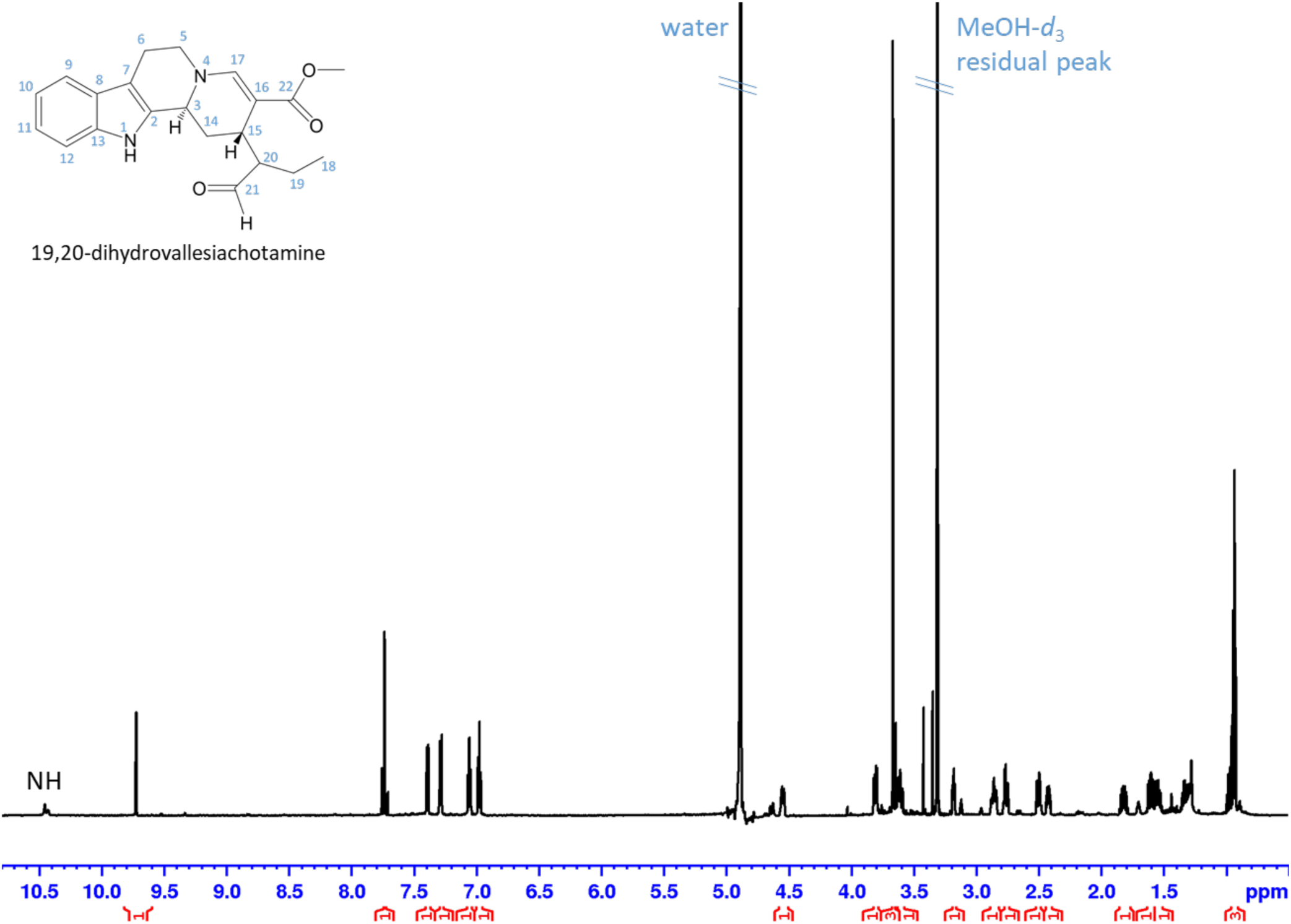
^1^H NMR data of 19,20-dihydrovallesiachotamine with water suppression, full range in MeOH-*d_3_*

**Figure S23.**
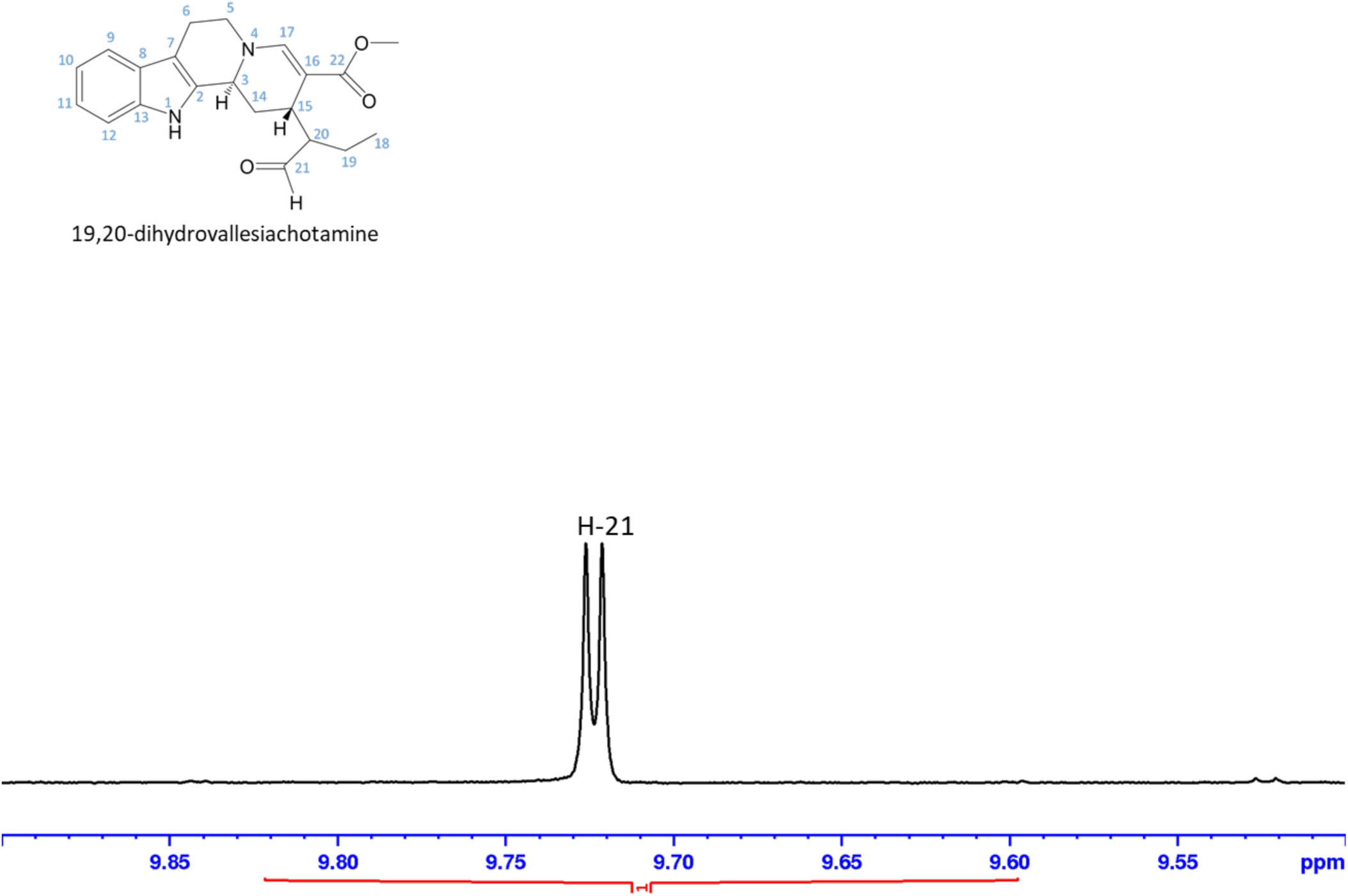
^1^H NMR data of 19,20-dihydrovallesiachotamine with water suppression, aldehyde range in MeOH-*d_3_*

**Figure S24.**
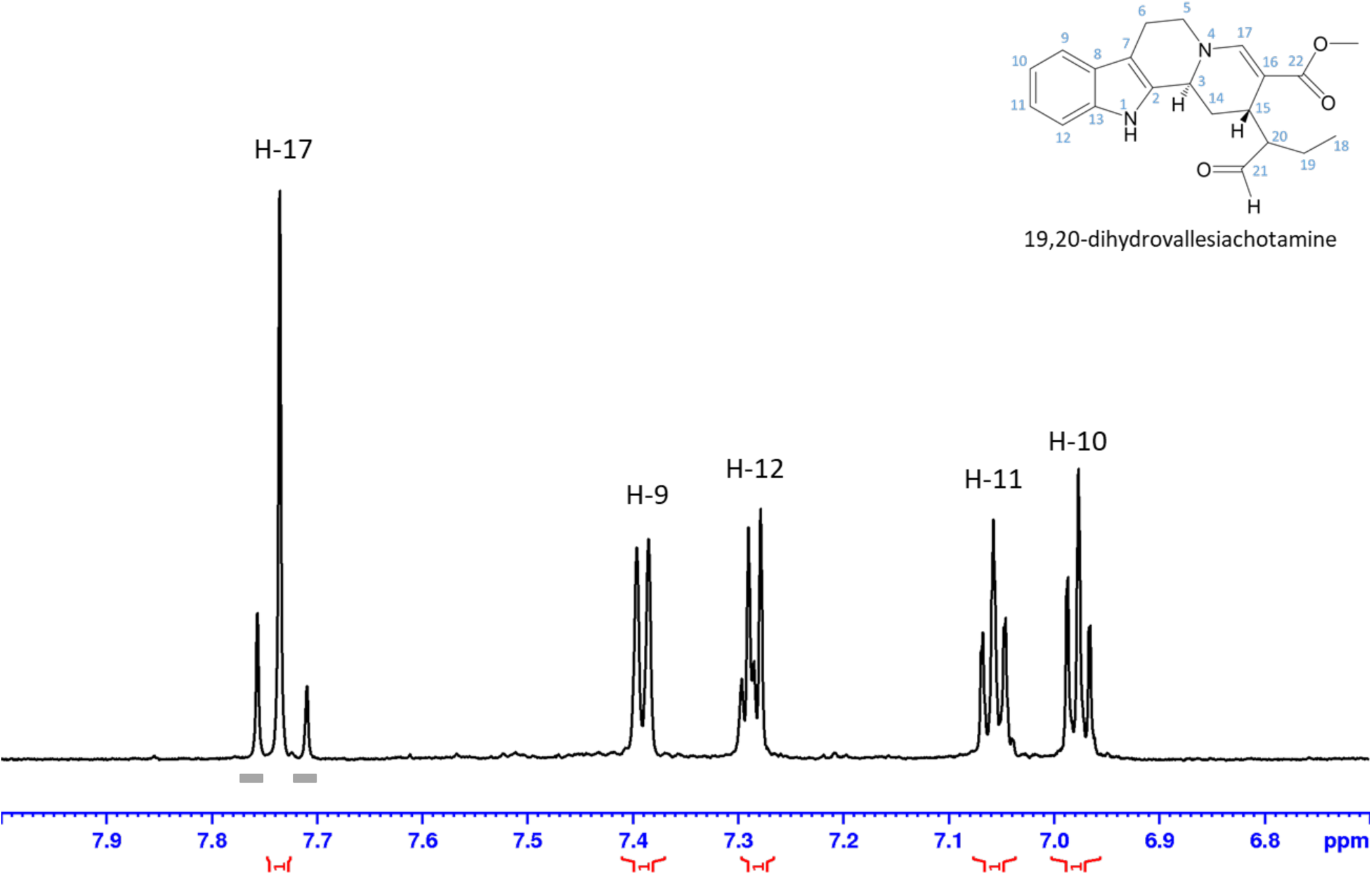
^1^H NMR data of 19,20-dihydrovallesiachotamine with water suppression, aromatic range in MeOH-*d_3_.* Grey bars indicate impurities.

**Figure S25.**
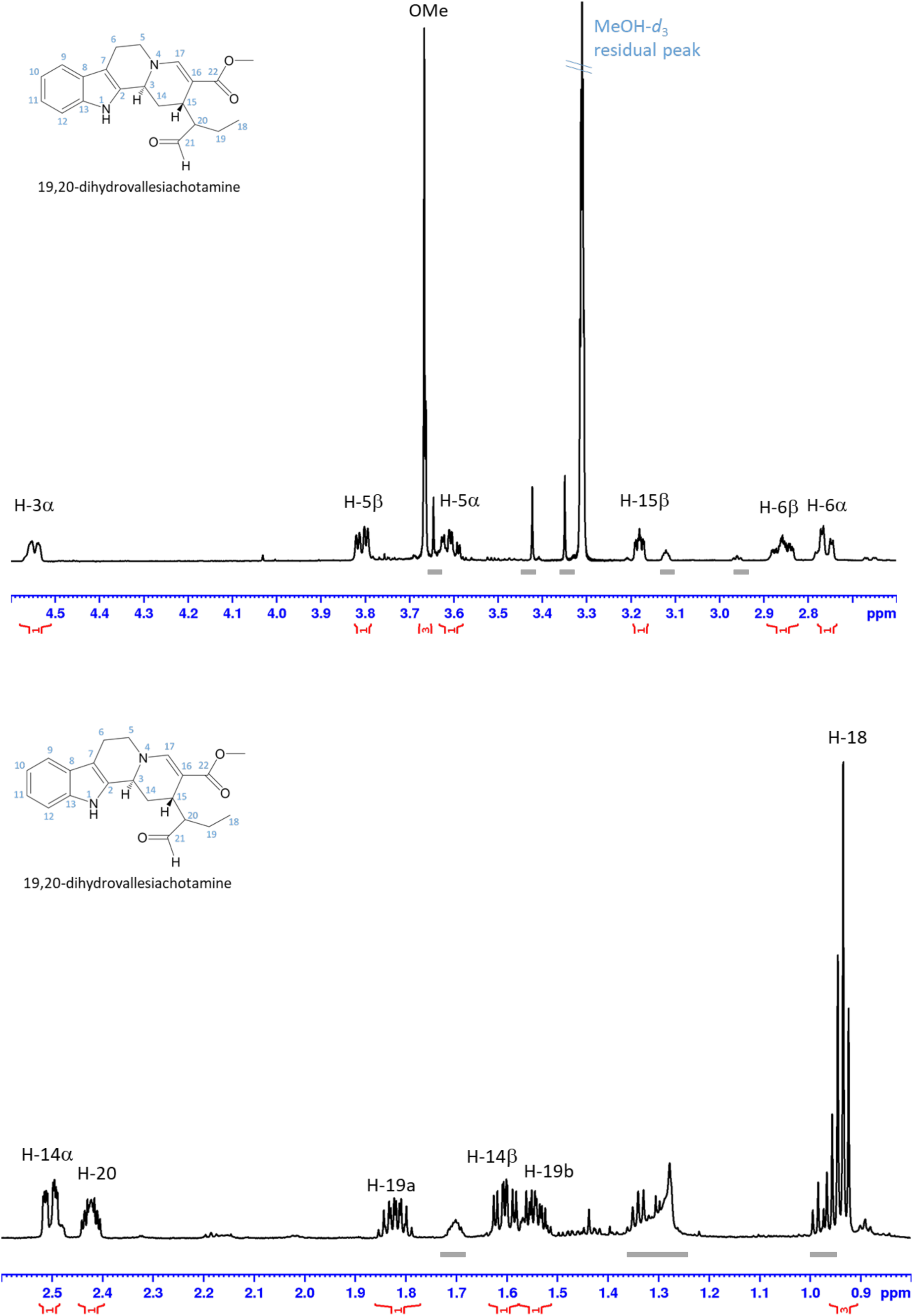
^1^H NMR data of 19,20-dihydrovallesiachotamine with water suppression, aliphatic range in MeOH-*d_3_.* Grey bars indicate impurities.

**Figure S26.**
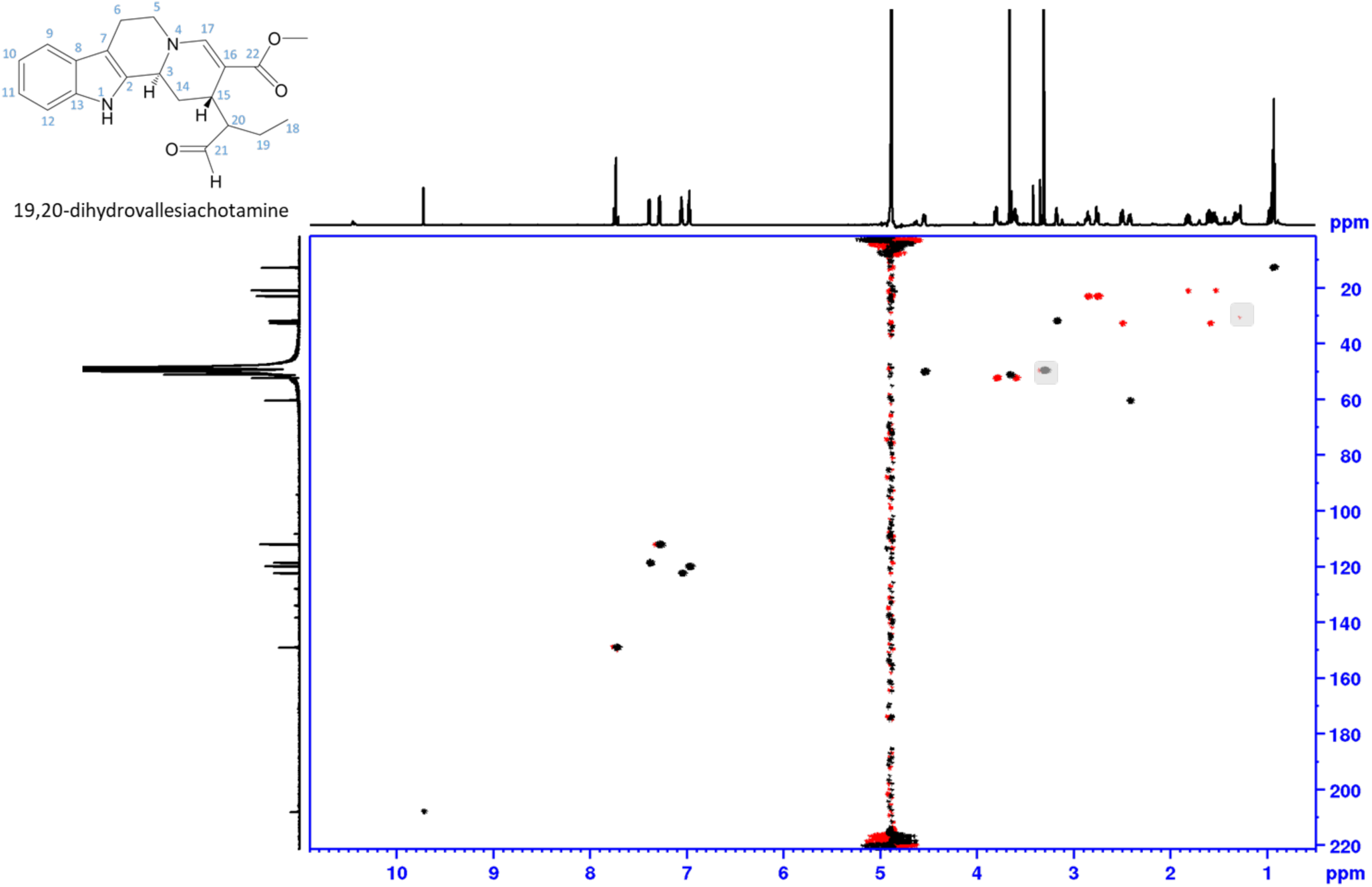
NMR data of 19,20-dihydrovallesiachotamine, phase sensitive HSQC, full range in MeOH-*d_3_*. Shaded areas mark impurity and solvent, red: CH2, black: CH, CH3

**Figure S27.**
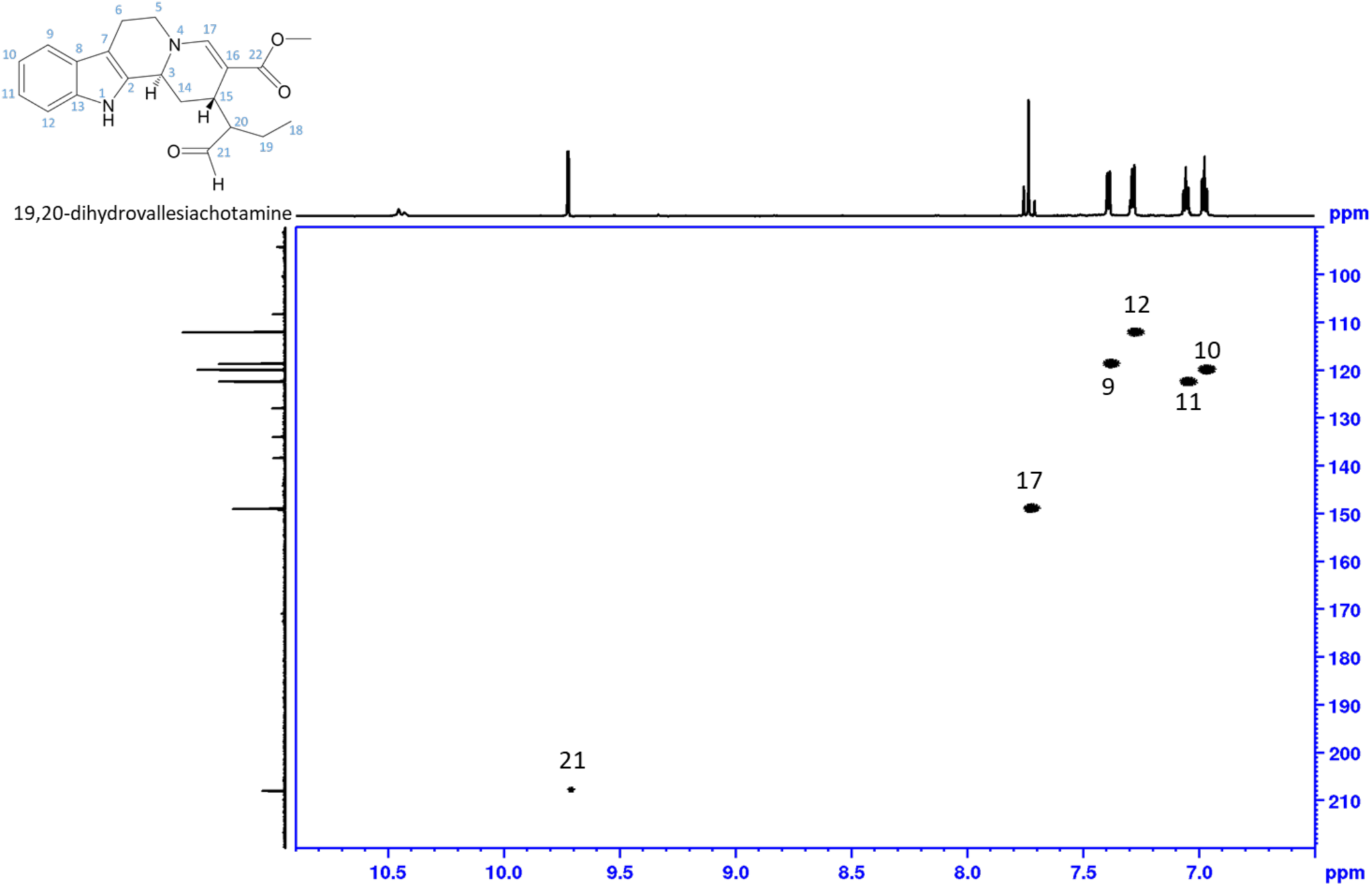
NMR data of 19,20-dihydrovallesiachotamine, phase sensitive HSQC, aldehyde and aromatic range in MeOH-*d_3_*. Shaded areas mark impurity and solvent, red: CH2, black: CH

**Figure S27.**
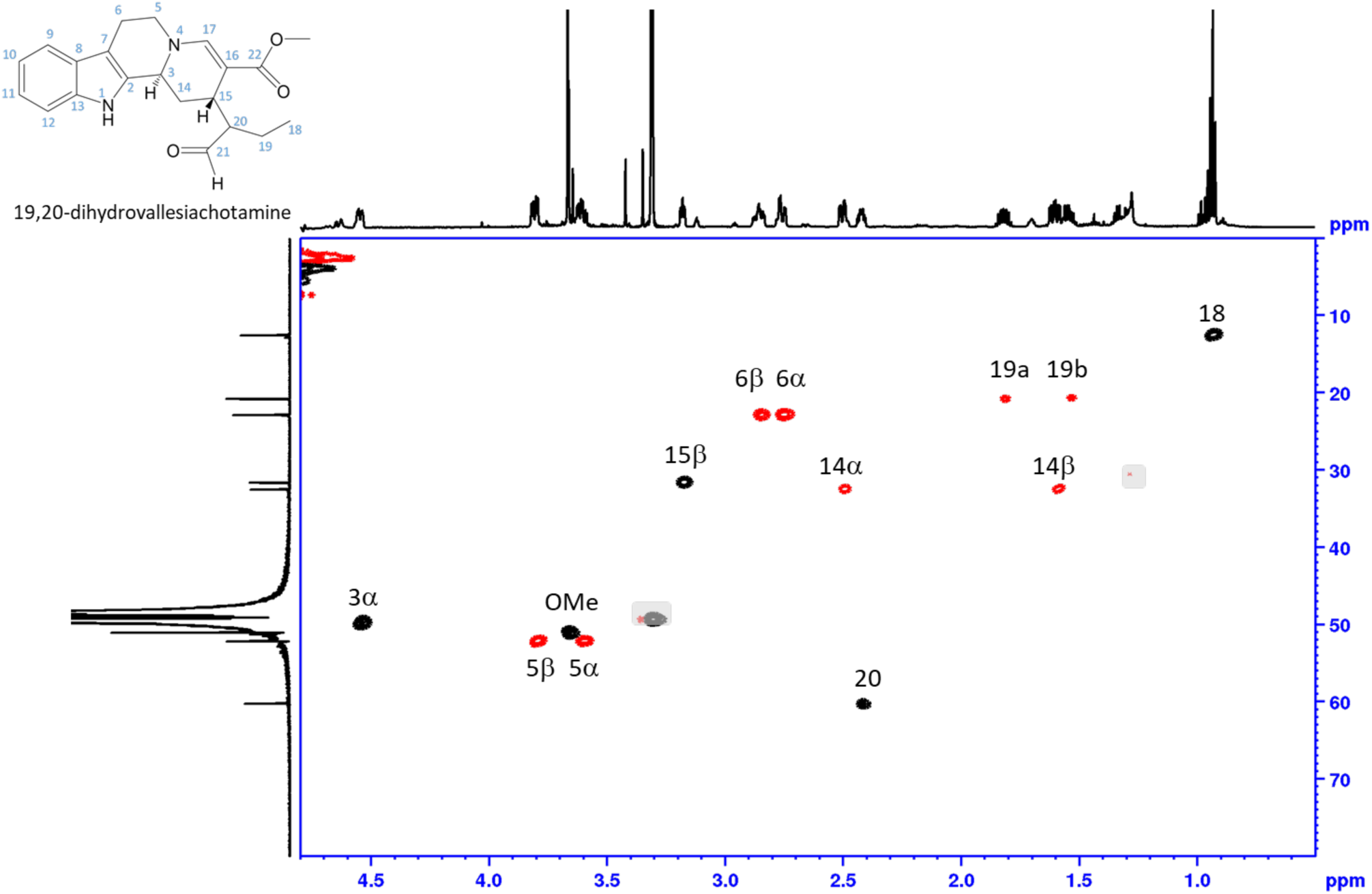
NMR data of 19,20-dihydrovallesiachotamine, phase sensitive HSQC, aliphatic range in MeOH-*d_3_*. Shaded areas mark impurity and solvent, red: CH2, black: CH

**Figure S28.**
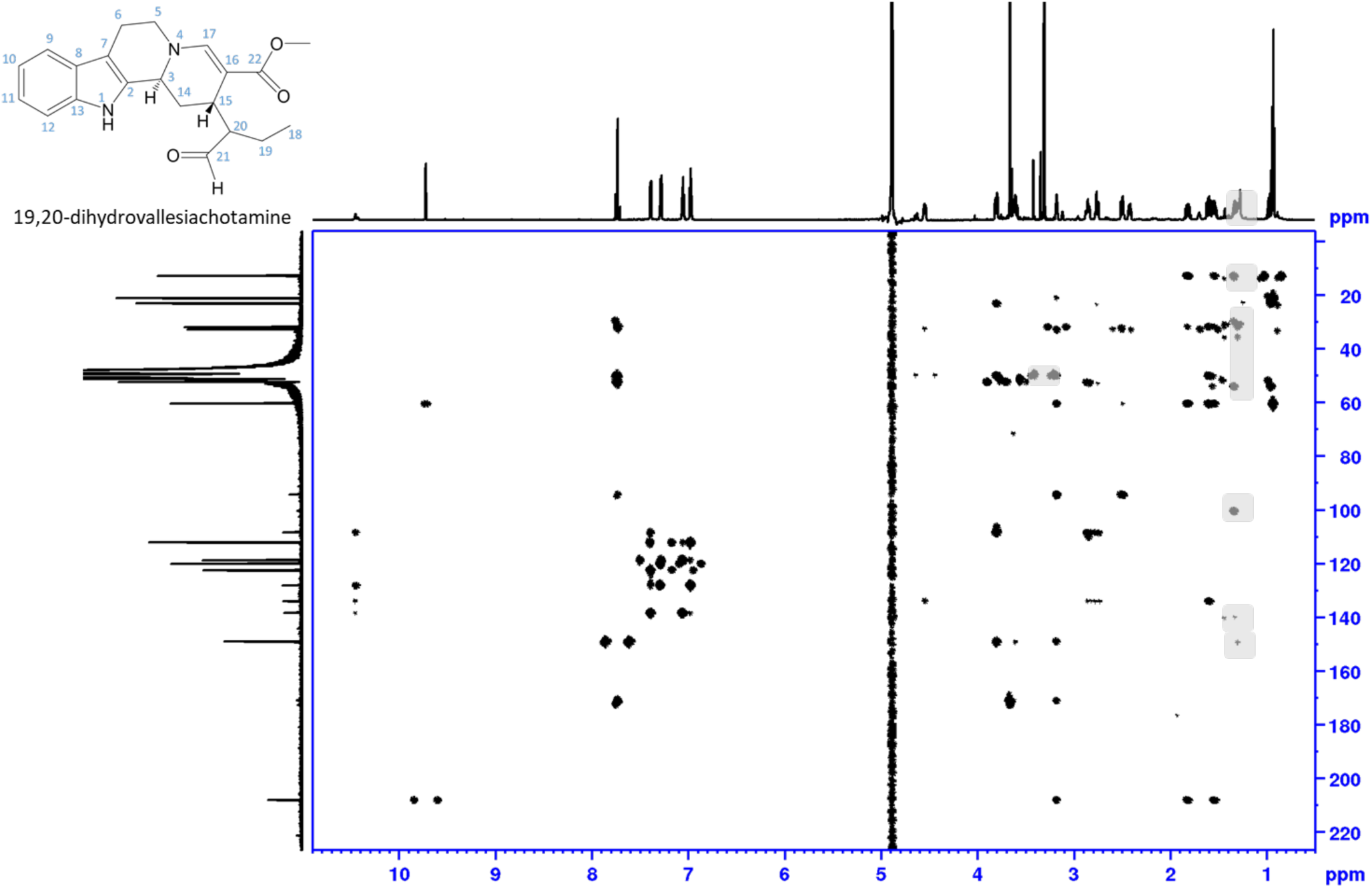
NMR data of 19,20-dihydrovallesiachotamine, HMBC, full range in MeOH-*d_3_*. Shaded areas mark impurity and solvent.

**Figure S29.**
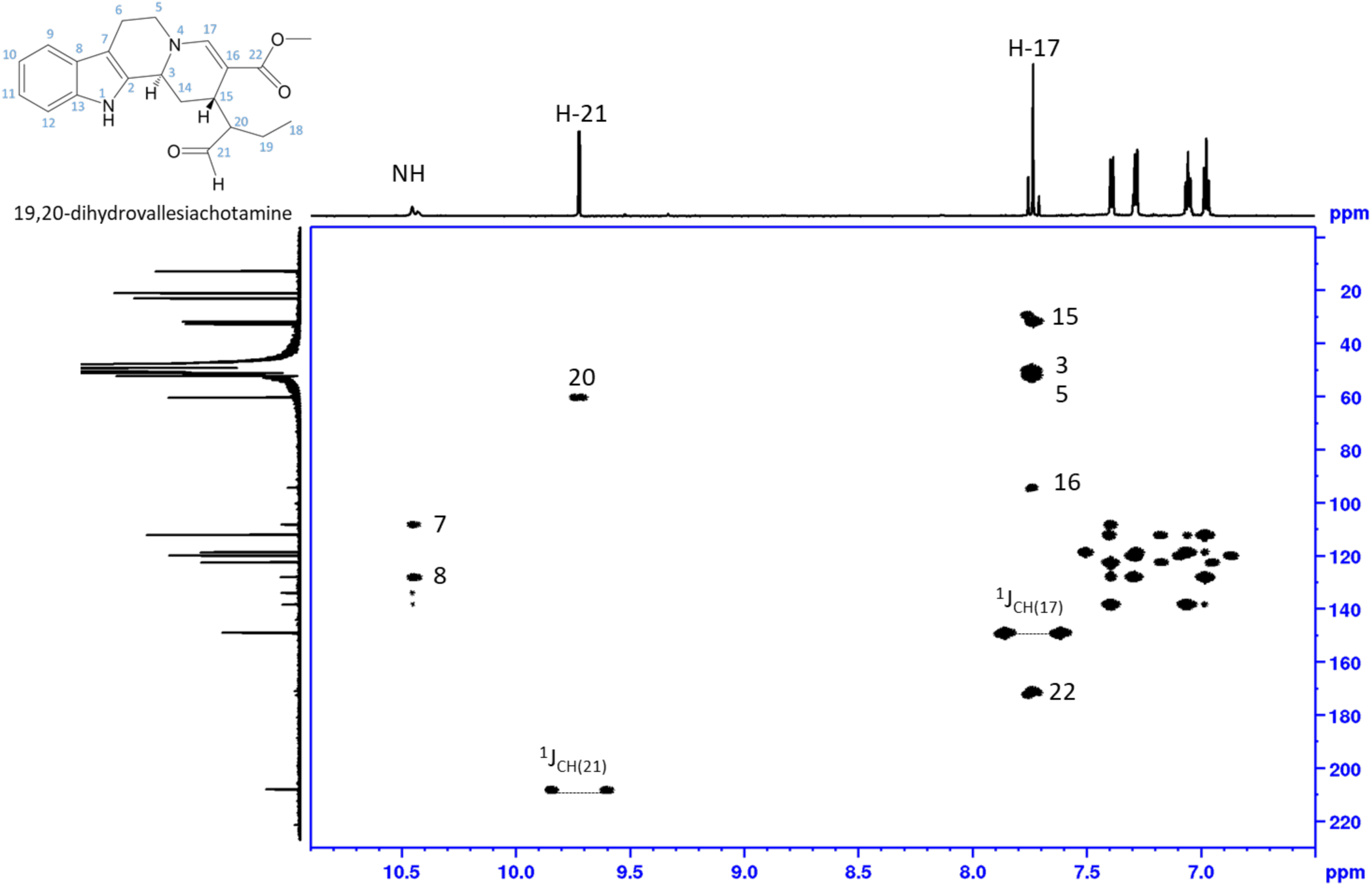
NMR data of 19,20-dihydrovallesiachotamine, HMBC, aldehyde and aromatic range in MeOH-*d_3_*. Shaded areas mark impurity and solvent.

**Figure S30.**
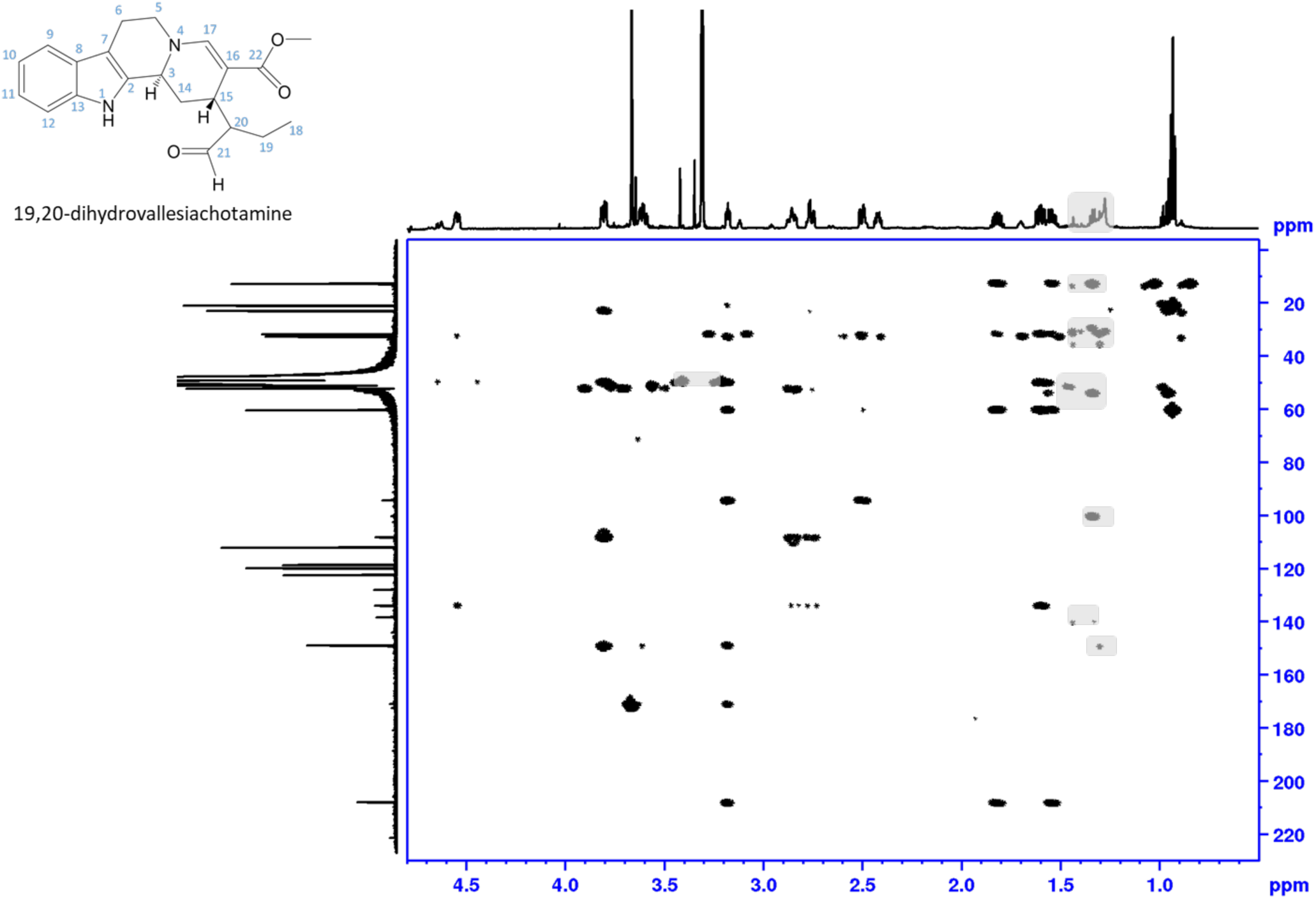
NMR data of 19,20-dihydrovallesiachotamine, HMBC, aliphatic range in MeOH-*d_3_*. Shaded areas mark impurity and solvent.

**Figure S31.**
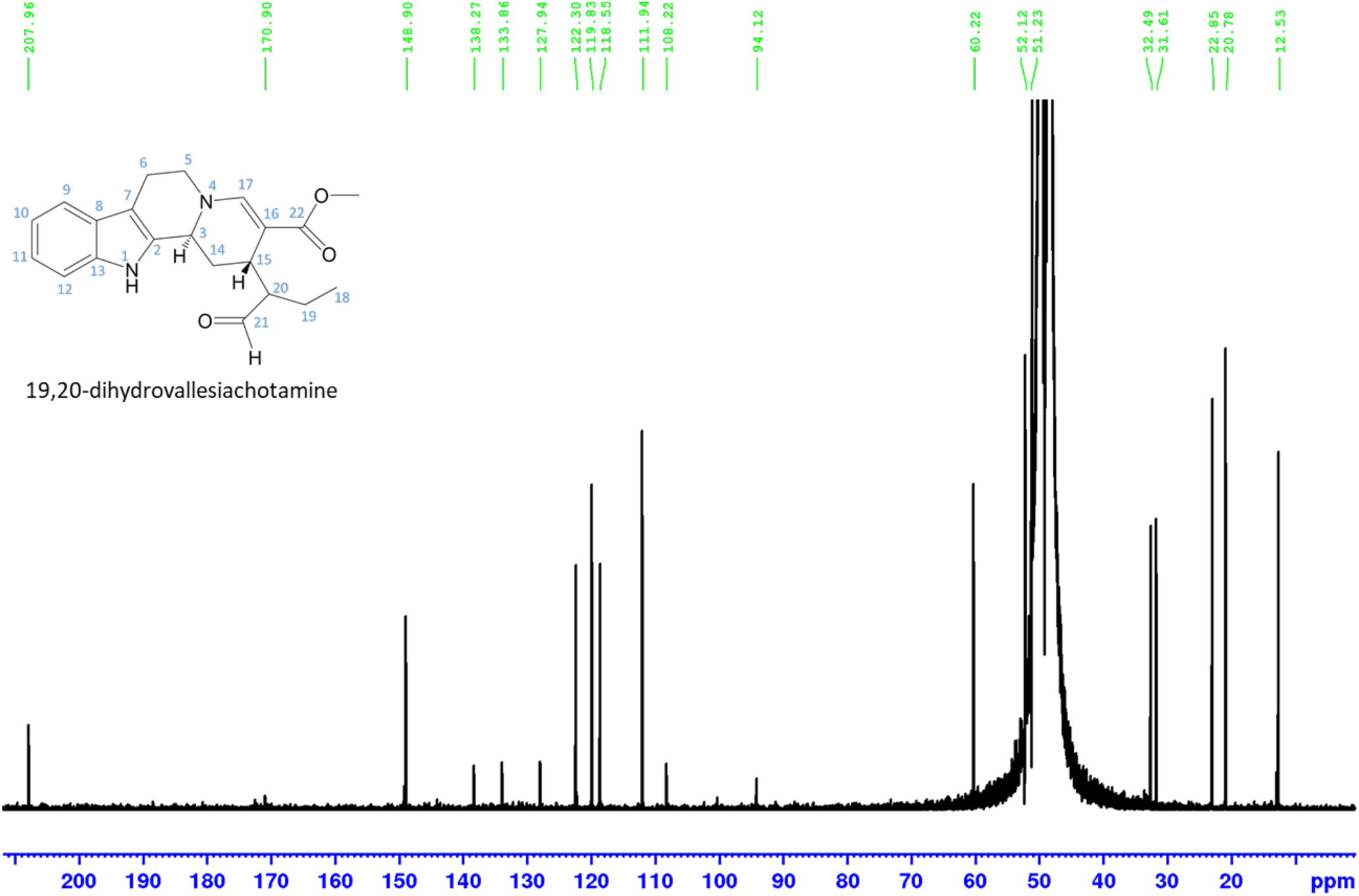
NMR data of 19,20-dihydrovallesiachotamine, DEPTQ, power spectrum, full range in MeOH-*d_3_*.

**Figure S32.**
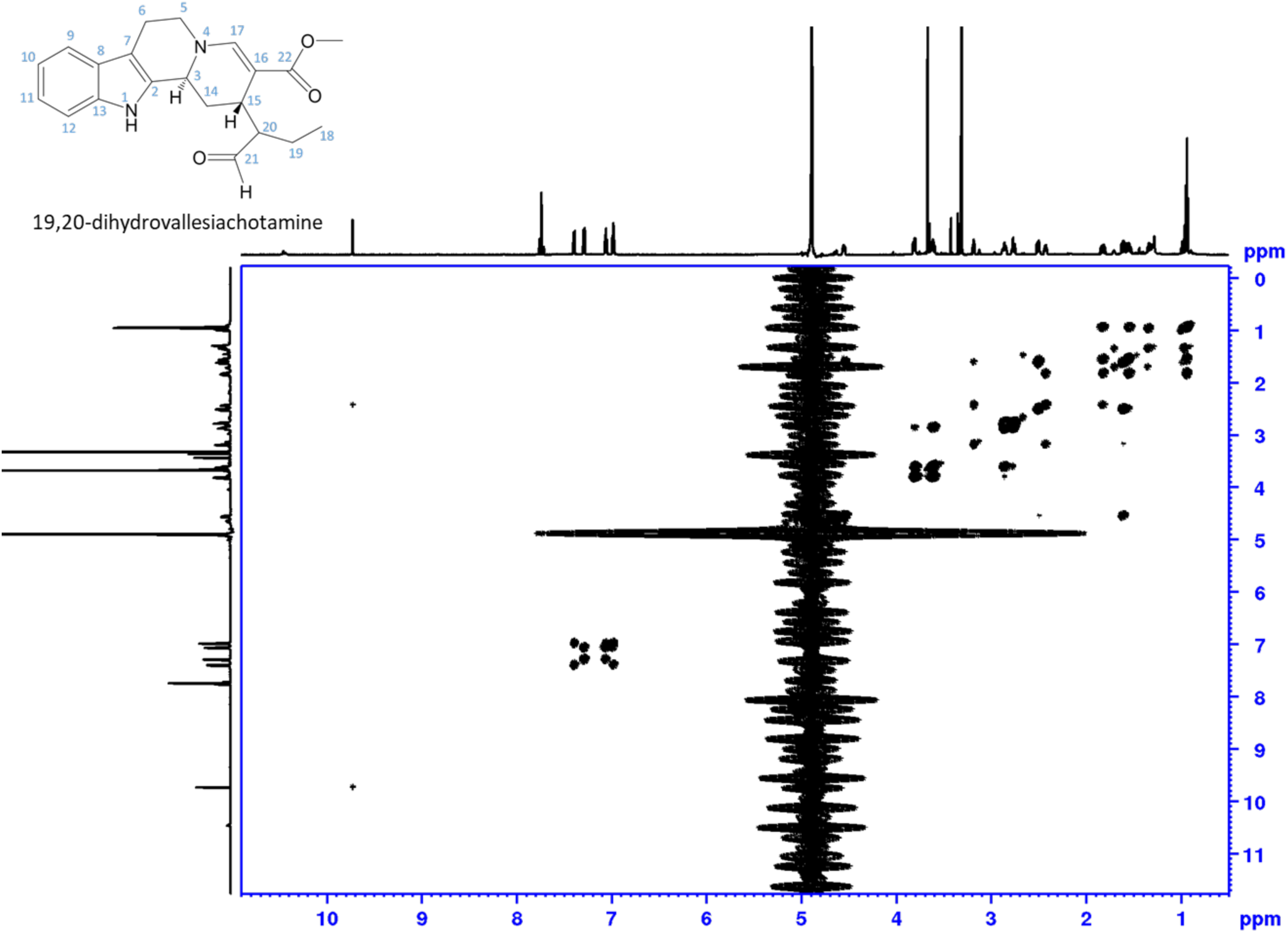
NMR data of 19,20-dihydrovallesiachotamine, 1H-1H DQF COSY with water suppression, magnitude mode processed, full range in MeOH-*d_3_*.

**Figure S33.**
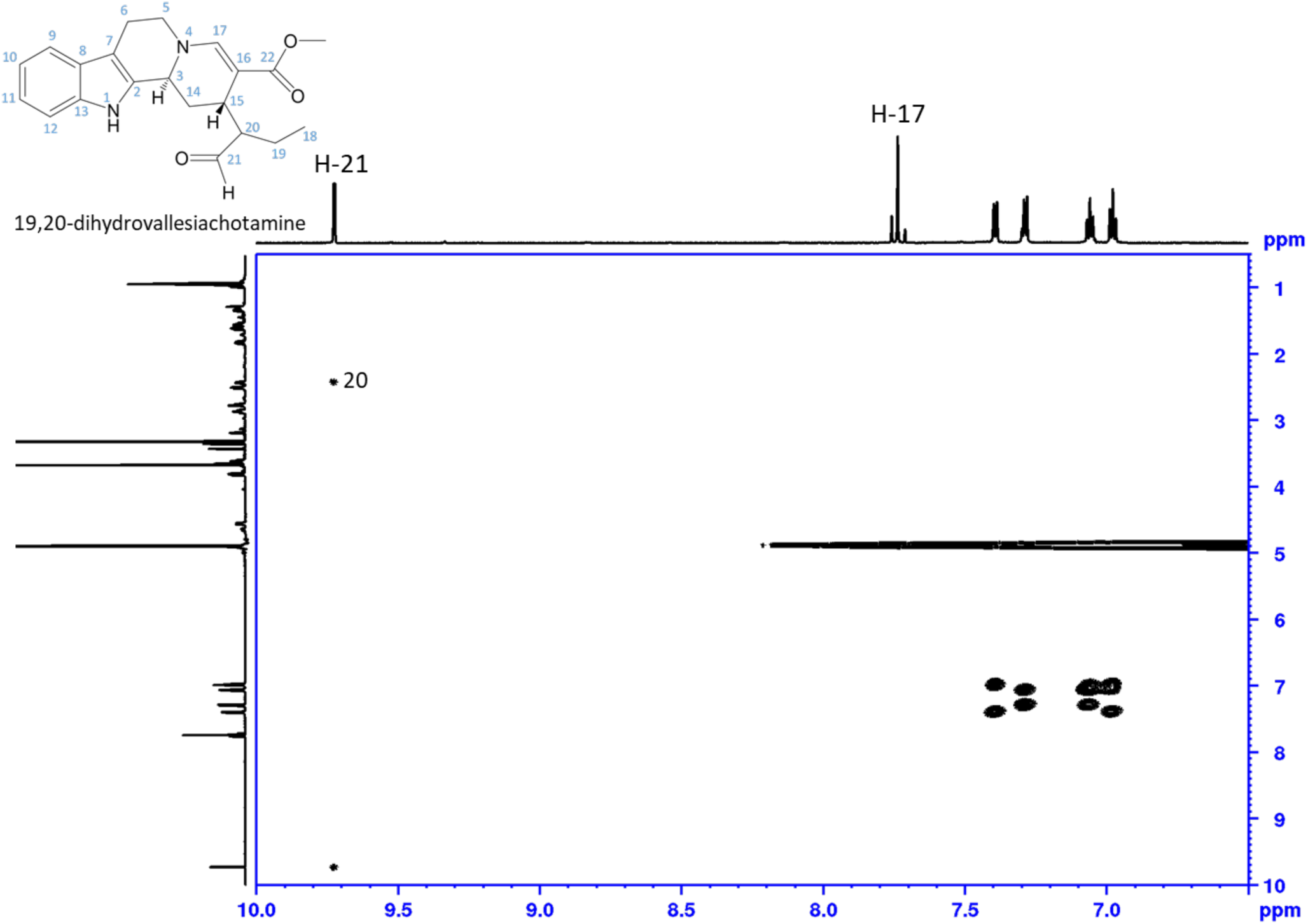
NMR data of 19,20-dihydrovallesiachotamine, 1H-1H DQF COSY with water suppression, magnitude mode processed, aldehyde and aromatic range in MeOH-*d_3_*.

**Figure S34.**
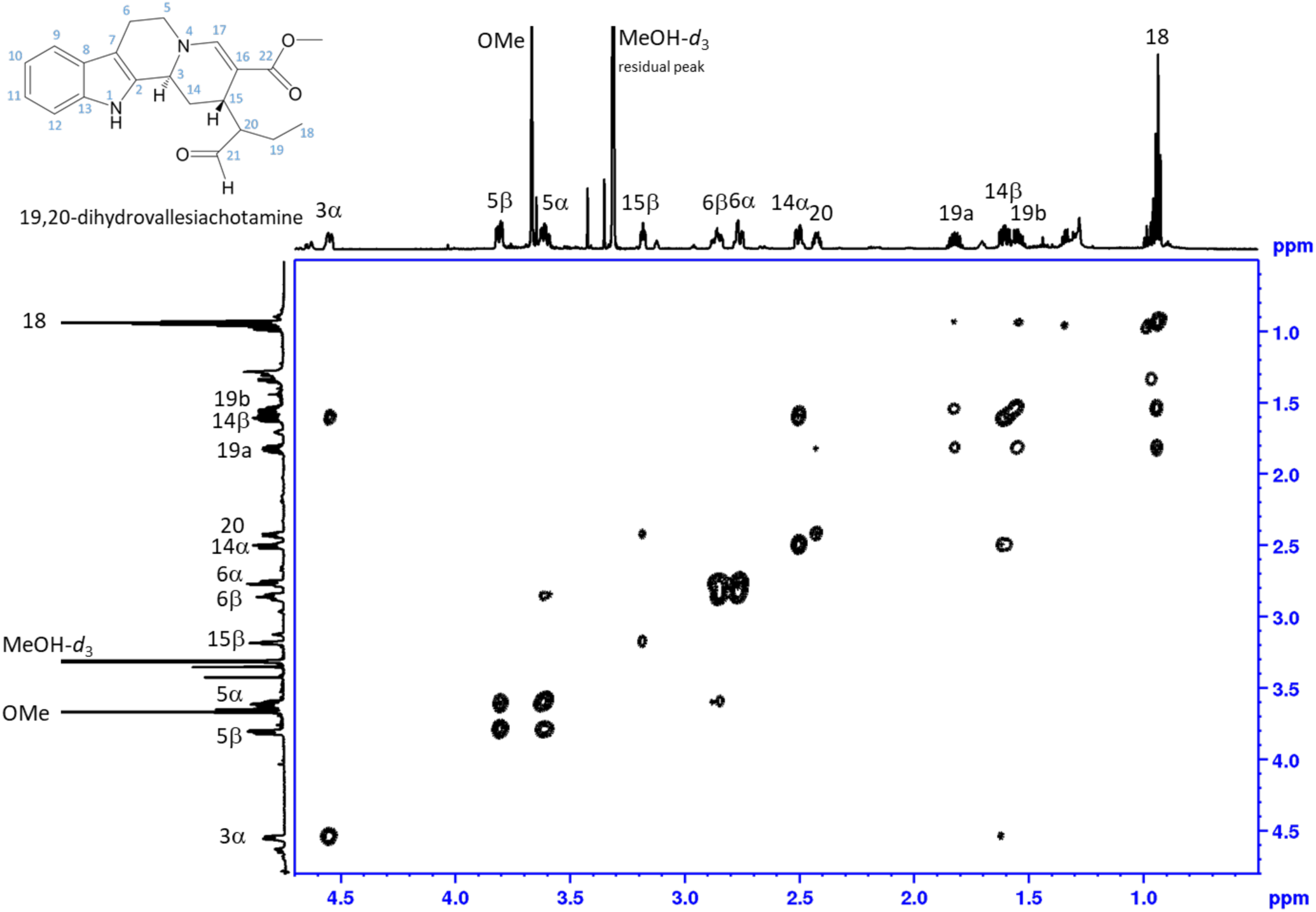
NMR data of 19,20-dihydrovallesiachotamine, 1H-1H DQF COSY with water suppression, magnitude mode processed, aliphatic range in MeOH-*d_3_*.

**Figure S35.**
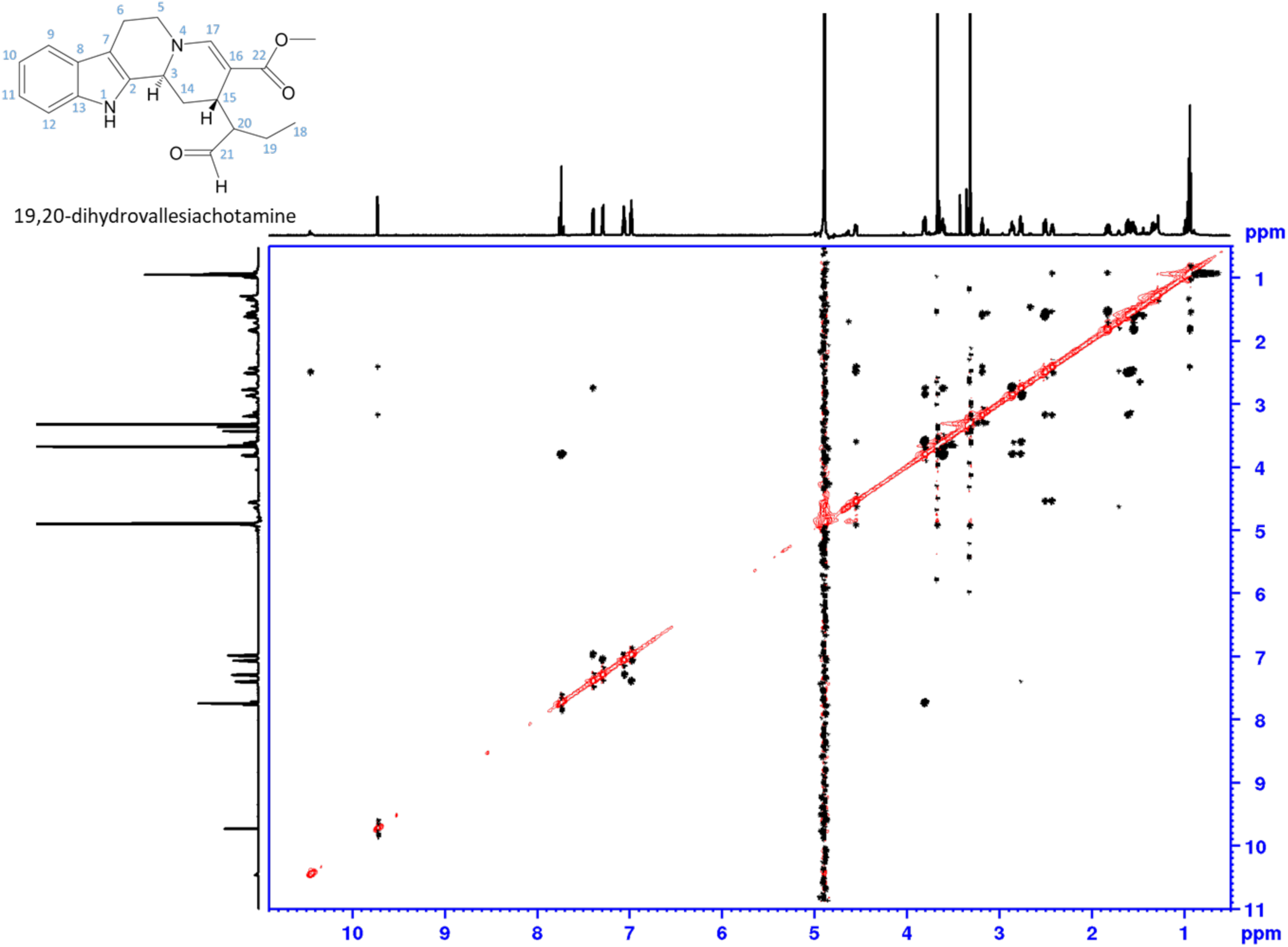
NMR data of 19,20-dihydrovallesiachotamine, ^1^H-^1^H ROESY with water suppression, full range in MeOH-*d*_3_

**Figure S36.**
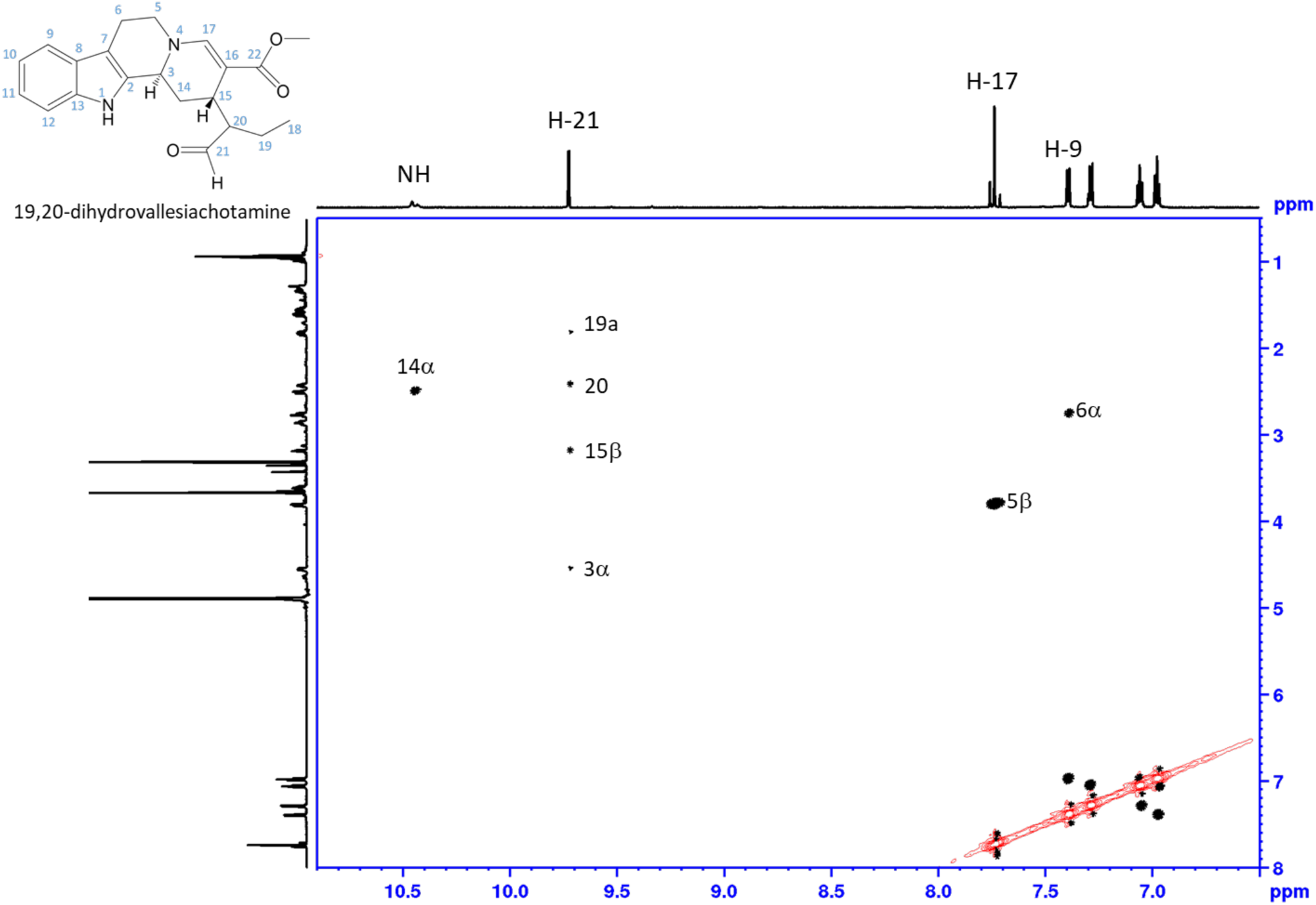
NMR data of 19,20-dihydrovallesiachotamine, ^1^H-^1^H ROESY with water suppression, aldehyde and aromatic range in MeOH-*d*_3_

**Figure S37.**
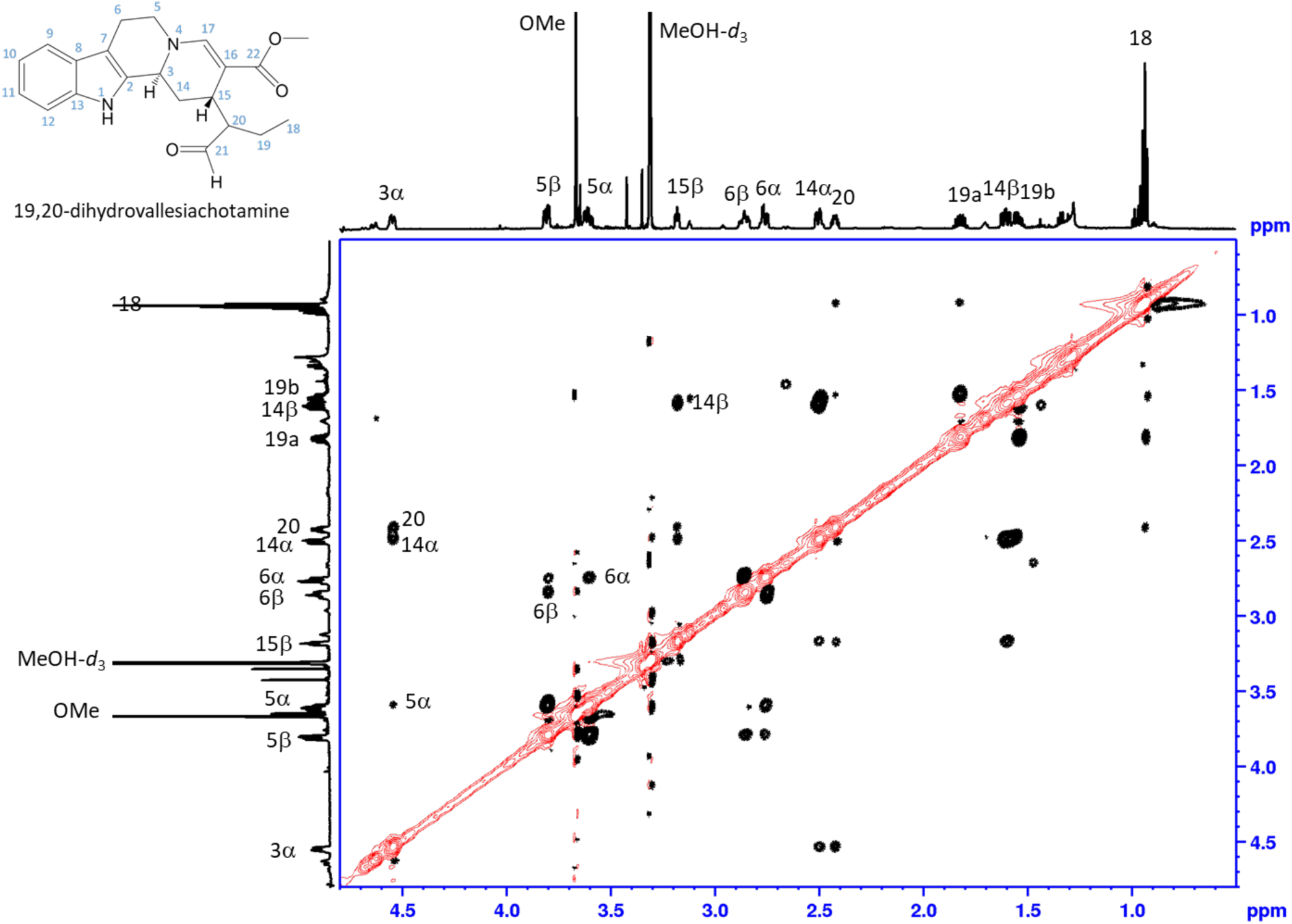
NMR data of 19,20-dihydrovallesiachotamine, ^1^H-^1^H ROESY with water suppression, aliphatic range in MeOH-*d*_3_

**Figure S38.**
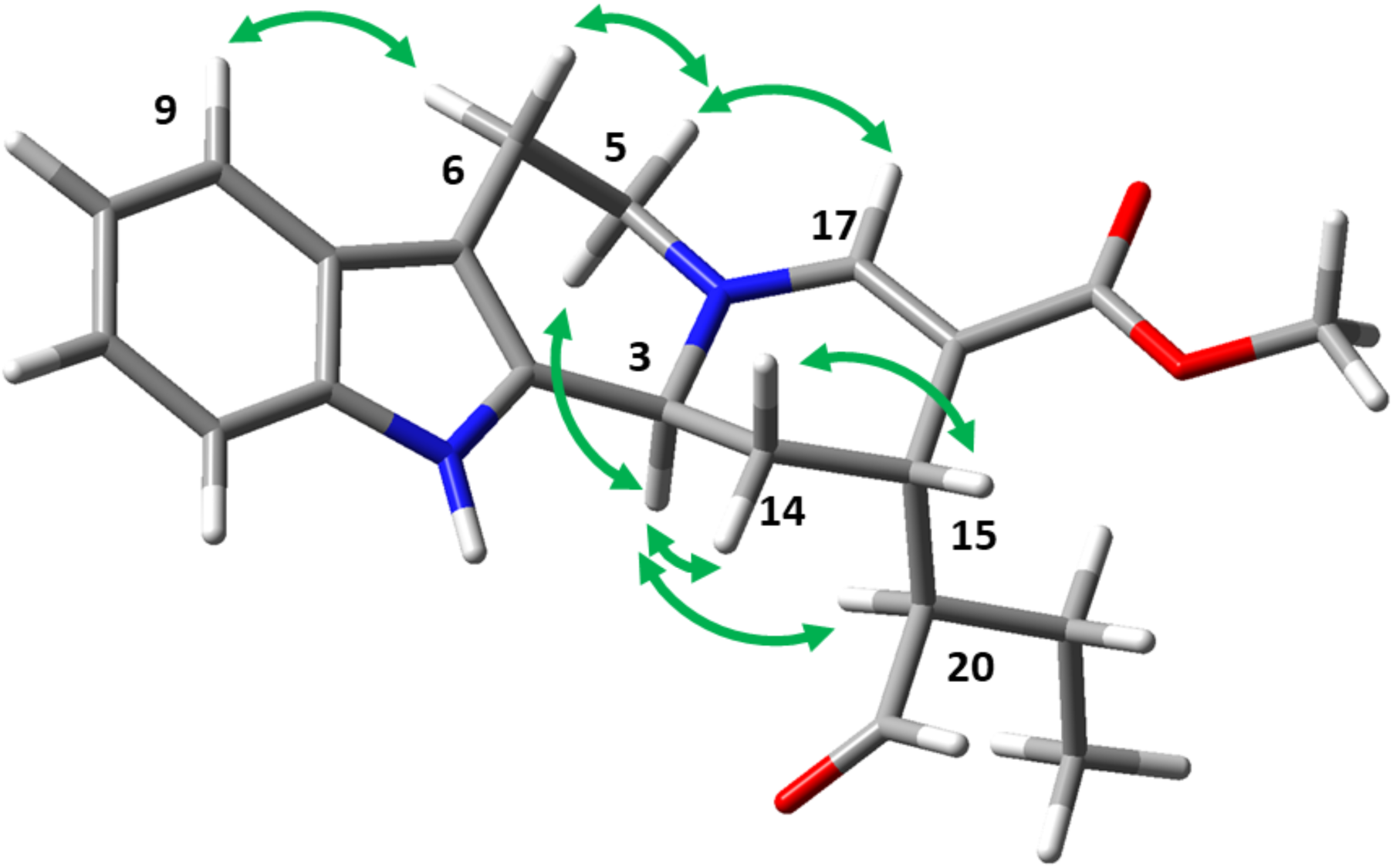
Structure of 19,20-dihydrovallesiachotamine optimized using Gaussian 16 (DFT APFD/6-311G++(2d,p), solvent MeOH). Important ROESY correlations extracted from NMR data are depicted in green.

**Figure S39.**
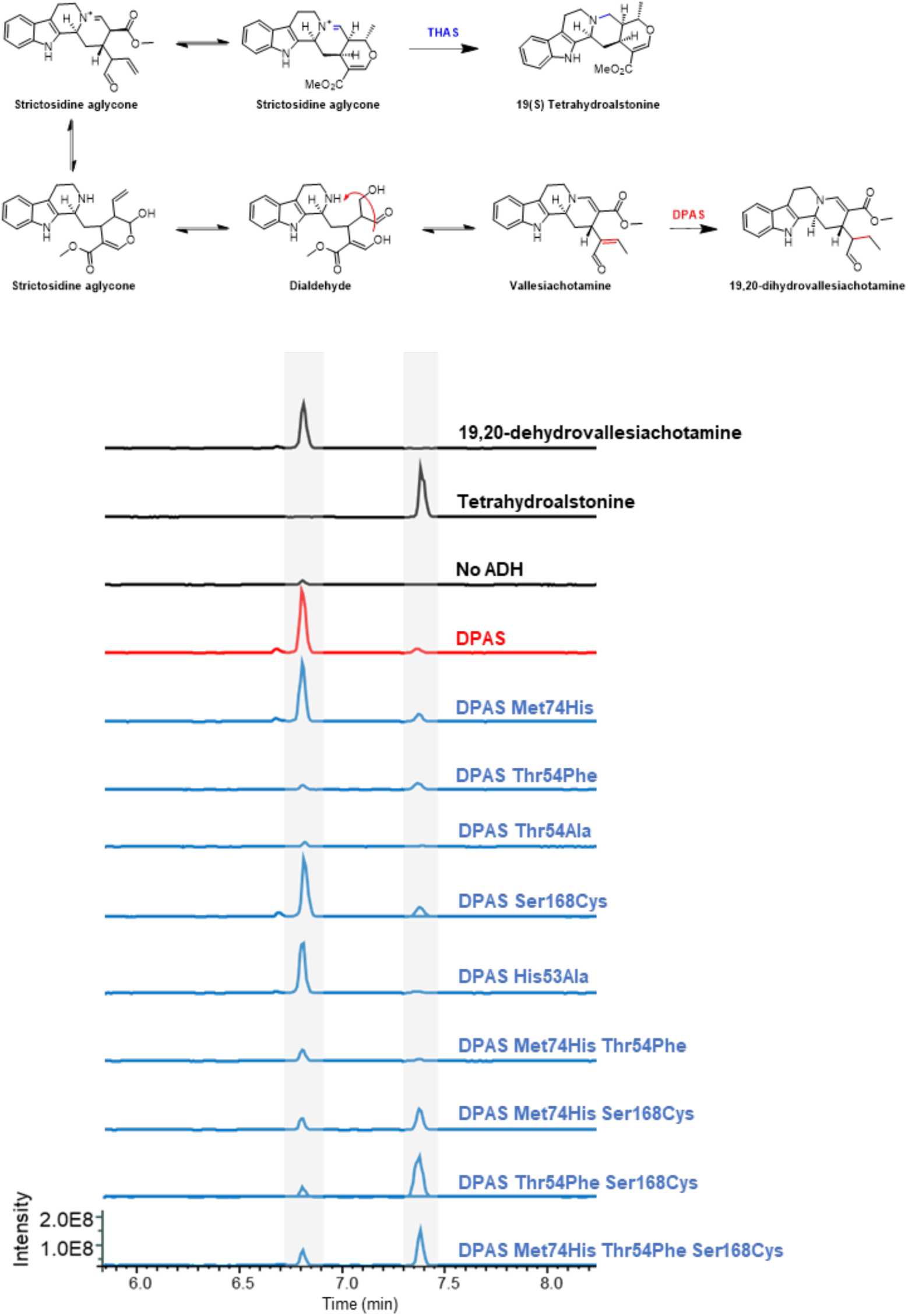
LC-MS chromatograms of *Cr*DPAS and mutants reacted with strictosidine aglycone. EIC *m/z* 353.185-353.225.

**Figure S40.**
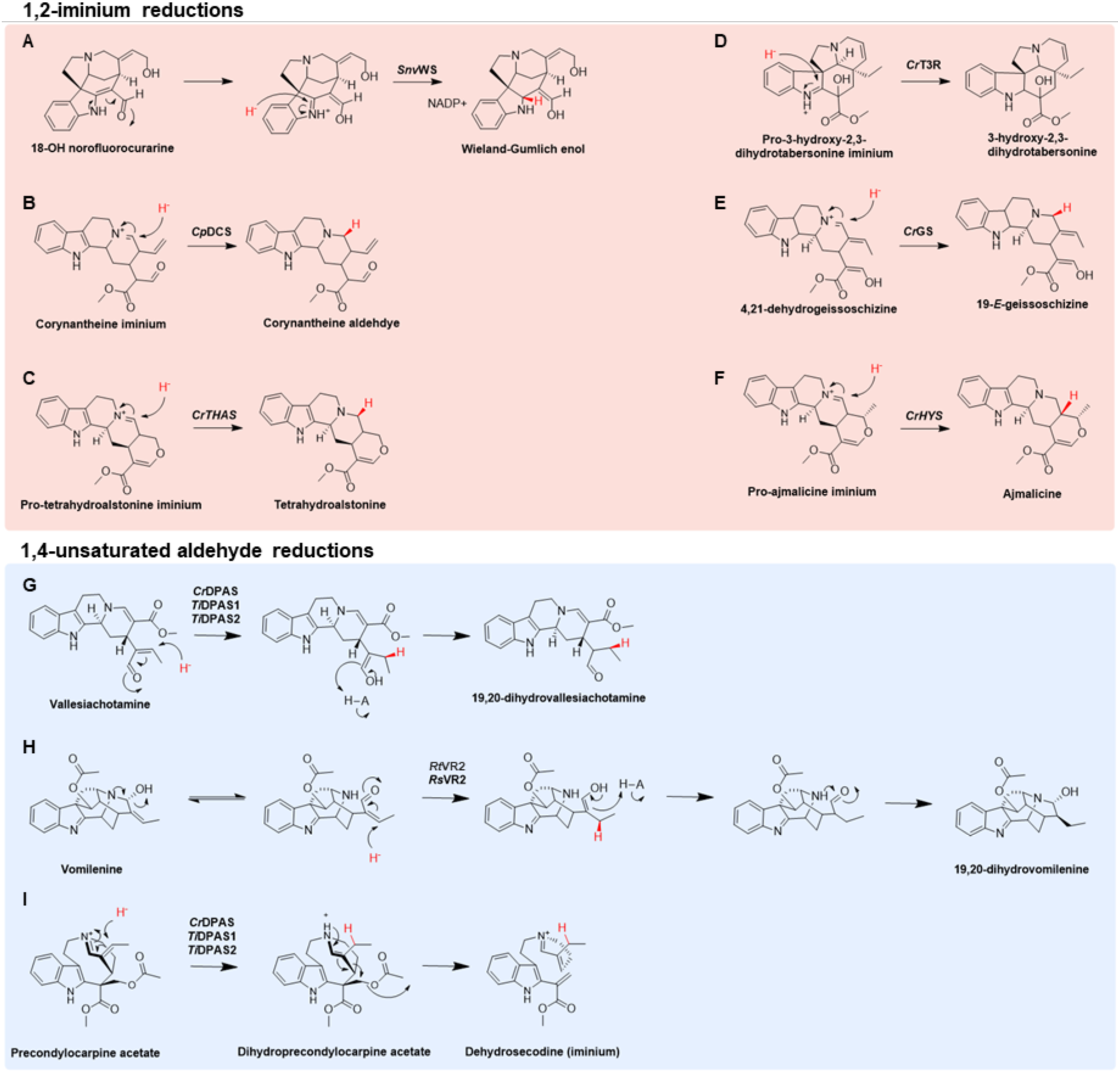
Previously characterised reactions in monoterpene indole alkaloid biosynthesis catalysed by ADHs. Reactions A-F are 1,2-iminium reductions, reactions G-H are 1,4 unsaturated aldehyde reductions and reaction I is 1,4-iminium reduction. **A.** *Strychnos nux-vomica* Wieland-Gumlich synthase (WS) ^[27]^; **B.** *Chinchona pubescens* dihydrocorynantheine aldehyde synthase (DCS) ^[28]^; **C.** *Catharanthus roseus* tetrahydroalstonine synthase (THAS) ^[4]^; **D.** *Catharanthus roseus* tabersonine-3-reductase (T3R) ^[29]^; **E.** *Catharanthus roseus* geissoschizine synthase (GS) ^[5]^; **F.** *Catharanthus roseus* heteroyohimbine synthase (HYS) ^[30]^; **G.** *Catharanthus roseus* and Tabernanthe iboga dihydroprecondylocarpine acetate synthase (DPAS), this study; **H.** *Rauwolfia serpentina* vomilenine reductase 2 (VR2) ^[31]^; **I.** *Catharanthus roseus* and *Tabernanthe iboga* dihydroprecondylocarpine acetate synthase (DPAS) ^[1,2]^.

## Supporting Tables

**Table S1.**
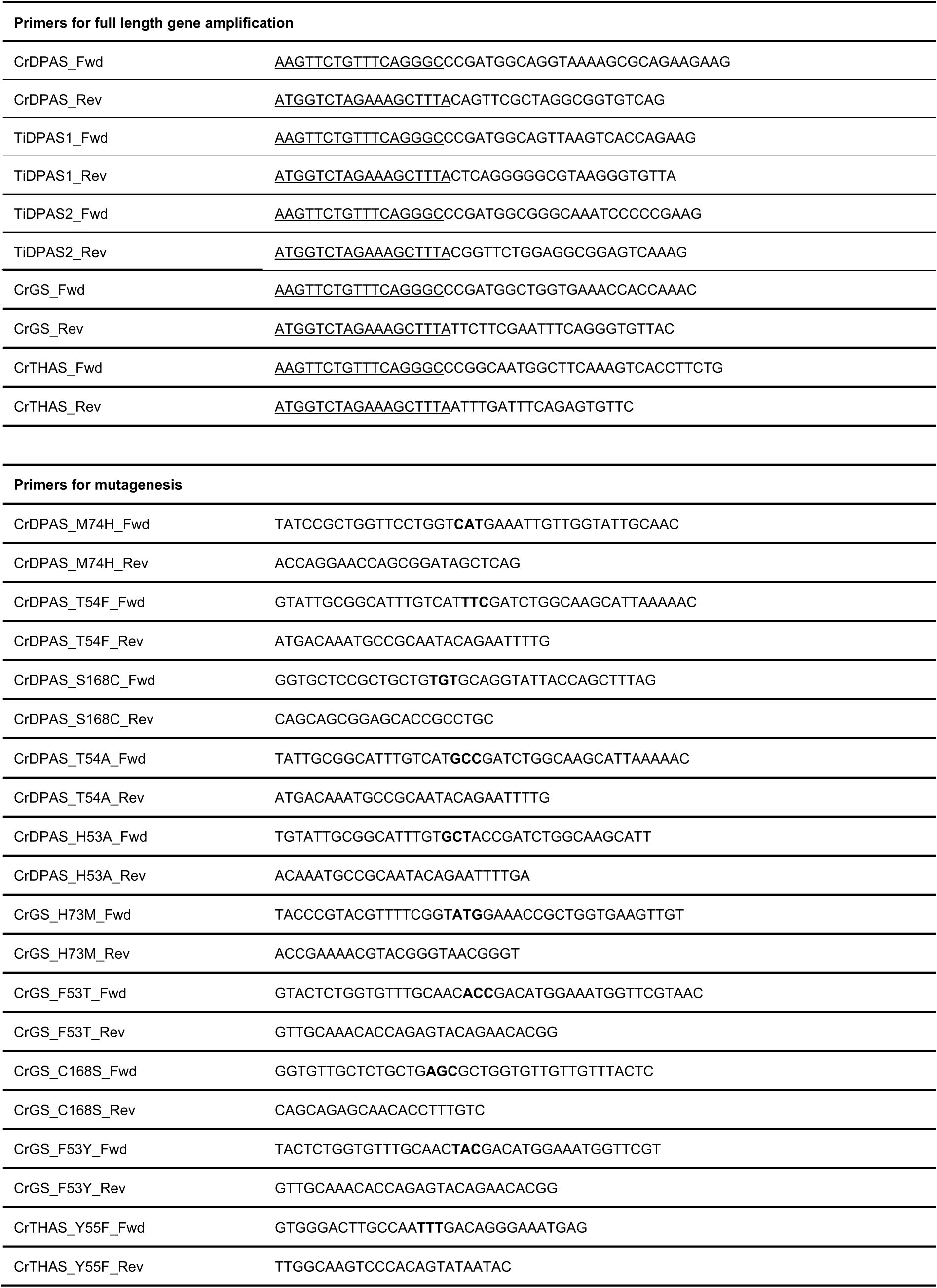
Primer sequences used in this study. Cloning overhangs are underlined. Mutated codons are in bold.

**Table S2.**
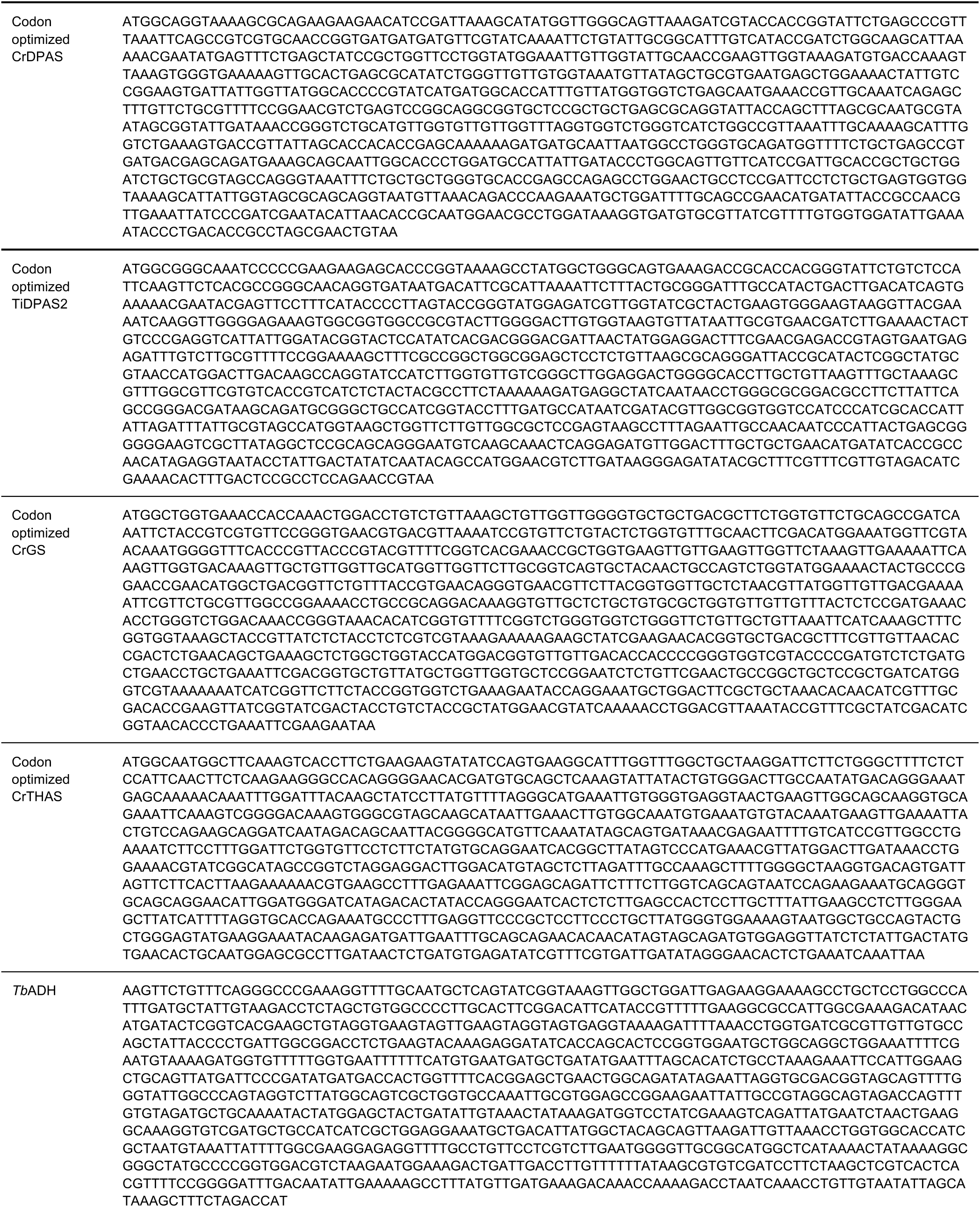
Full length sequences of codon optimized genes used in this study.

**Table S3.** Summary of X-ray data and model parameters for *Cr*DPAS

|  |  |
| --- | --- |
| Data collection |  |
| Paul Scherrer Institute | 10SA (PX II) |
| Wavelength (Å) | 1 |
| Resolution range (Å) | 44.62 - 2.45 (2.548 - 2.45) |
| Space Group | *P* 21 21 21 |
| Cell parameters (Å) | *a* = 61.019, *b* = 114.015, *c* = 143.357, β = 90° |
| Total no. of measured reflections | 494135 (51201) |
| Unique reflections | 37564 (3719) |
| Multiplicity | 13.2 (13.8) |
| Mean *I*/σ(*I*) | 21.46 (3.10) |
| Completeness (%) | 98.7 (96.8) |
| *R*<sub>merge</sub><sup>a</sup> | 0.2154 (1.406) |
| *R*<sub>meas</sub><sup>b</sup> | 0.2242 (1.46) |
| *CC*<sub>1/2</sub><sup>c</sup> | 0.999 (0.879) |
| Wilson *B* value (Å<sup>2</sup>) | 53.18 |
| Refinement |  |
| Reflections used in refinement | 37560 (3719) |
| Reflections used for R-free | 1877 (186) |
| *R*<sub>work</sub> | 0.2217 (0.2772) |
| *R*<sub>free</sub> | 0.2501 (0.3143) |
| *CC*<sub>work</sub> | 0.942 (0.845) |
| *CC*<sub>free</sub> | 0.958 (0.704) |
| Protein residues | 640 |
| Number of non-hydrogen atoms | 4668 |
| macromolecules | 4566 |
| ligands | 43 |
| solvent | 69 |
| Ramachandran plot: favoured/allowed/disallowed<sup>f</sup> (%) | 98.1/1.58/0.32 |
| Rotamer outliers (%) | 3.97 |
| R.m.s. bond distance deviation (Å) | 0.007 |
| R.m.s. bond angle deviation (°) | 0.98 |
| Clashscore | 19.95 |
| Mean *B* factors: protein/waters/ligands/overall (Å<sup>2</sup>) | 62.93/54.82/76.91/62.91 |
| PDB accession code | --- |

Statistics for the highest-resolution shell are shown in parentheses.
<sup>a</sup> $R_{\text{merge}} = \frac{\sum_{hkl} \sum_i |I_i(hkl) - \langle I(hkl) \rangle|}{\sum_{hkl} \sum_i I_i(hkl)}$ .
<sup>b</sup> $R_{\text{meas}} = \frac{\sum_{hkl} [N/(N-1)]^{1/2} \times \sum_i |I_i(hkl) - \langle I(hkl) \rangle|}{\sum_{hkl} \sum_i I_i(hkl)}$ , where $I_i(hkl)$ is the *i*th observation of reflection *hkl*, $\langle I(hkl) \rangle$ is the weighted average intensity for all observations *i* of reflection *hkl* and *N* is the number of observations of reflection *hkl*.
<sup>c</sup> *CC*<sub>1/2</sub> is the correlation coefficient between symmetry equivalent intensities from random halves of the dataset.
<sup>d</sup> The data set was split into "working" and "free" sets consisting of 95 and 5% of the data respectively. The free set was not used for refinement.
<sup>e</sup> The R-factors *R*<sub>work</sub> and *R*<sub>free</sub> are calculated as follows: $R = \frac{\sum (|F_{\text{obs}} - F_{\text{calc}}|)}{\sum |F_{\text{obs}}|}$ , where *F*<sub>obs</sub> and *F*<sub>calc</sub> are the observed and calculated structure factor amplitudes, respectively.
<sup>f</sup> As calculated using MolProbity [32].

**Table S4.**
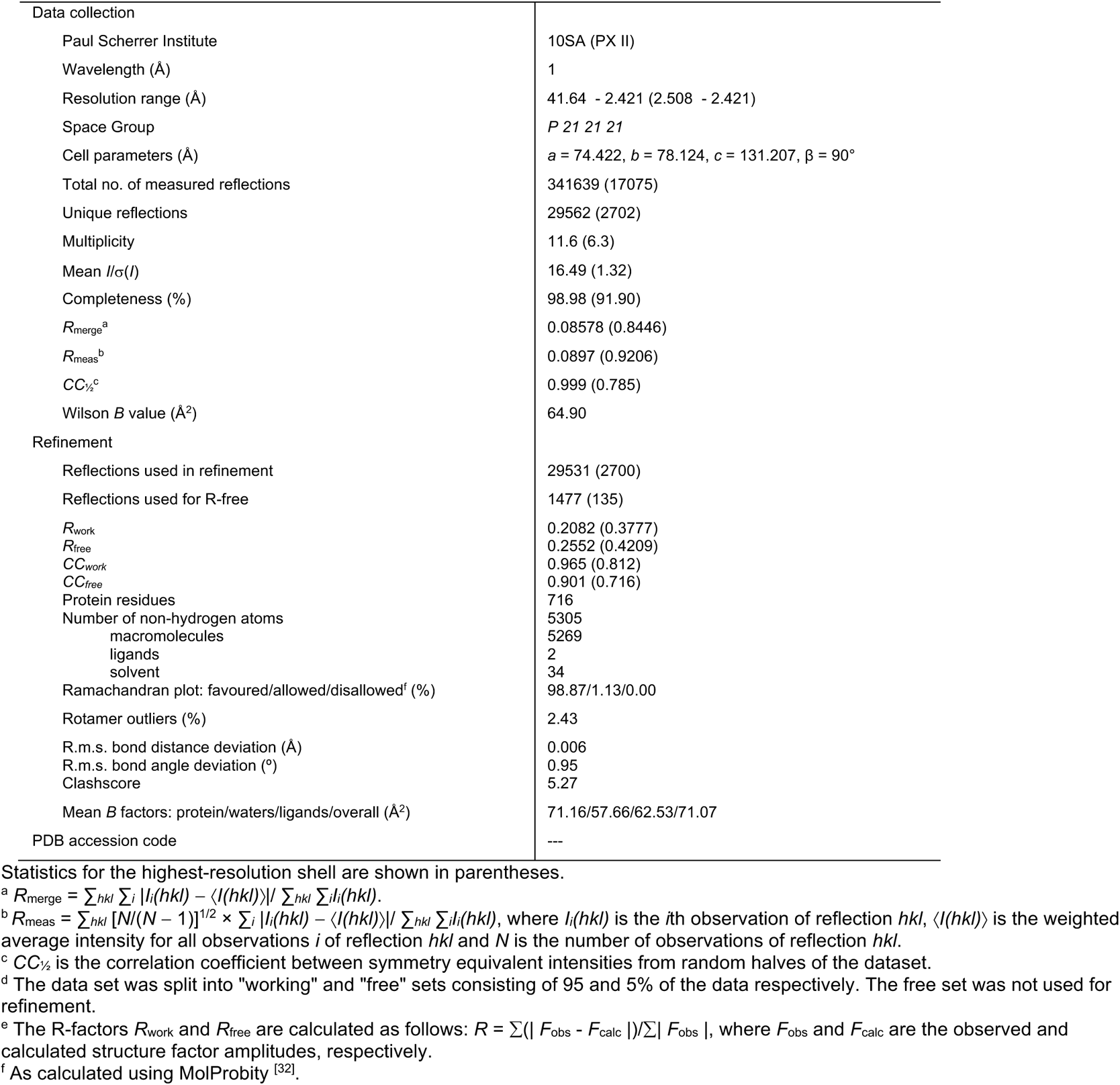
Summary of X-ray data and model parameters for apo-*Ti*DPAS2

**Table S5.** Summary of X-ray data and model parameters for precondylocarpine acetate-bound *Ti*DPAS2

|  |  |
| --- | --- |
| Data collection |  |
| Paul Scherrer Institute | 10SA (PX II) |
| Wavelength (Å) | 1 |
| Resolution range (Å) | 39.81 – 1.882 (1.949 – 1.882) |
| Space Group | *P* 21 21 21 |
| Cell parameters (Å) | *a* = 72.888, *b* = 79.624, *c* = 130.801, β = 90° |
| Total no. of measured reflections | 809479 (78567) |
| Unique reflections | 62174 (5895) |
| Multiplicity | 13.0 (13.3) |
| Mean *I*/σ(*I*) | 14.05 (0.88) |
| Completeness (%) | 99.49 (95.74) |
| *R*<sub>merge</sub><sup>a</sup> | 0.1082 (3.23) |
| *R*<sub>meas</sub><sup>b</sup> | 0.1128 (3.357) |
| *CC*<sub>1/2</sub><sup>c</sup> | 0.999 (0.463) |
| Wilson *B* value (Å<sup>2</sup>) | 40.94 |
| Refinement |  |
| Reflections used in refinement | 62152 (5895) |
| Reflections used for R-free | 3104 (295) |
| *R*<sub>work</sub> | 0.1927 (0.4735) |
| *R*<sub>free</sub> | 0.2216 (0.5240) |
| *CC*<sub>work</sub> | 0.972 (0.696) |
| *CC*<sub>free</sub> | 0.966 (0.671) |
| Protein residues | 716 |
| Number of non-hydrogen atoms | 5601 |
| macromolecules | 5272 |
| ligands | 97 |
| solvent | 242 |
| Ramachandran plot: favoured/allowed/disallowed<sup>f</sup> (%) | 97.33/2.67/0.00 |
| Rotamer outliers (%) | 0.93 |
| R.m.s. bond distance deviation (Å) | 0.004 |
| R.m.s. bond angle deviation (°) | 0.71 |
| Clashscore | 3.89 |
| Mean *B* factors: protein/waters/ligands/overall (Å<sup>2</sup>) | 44.88/47.01/46.74/45.00 |
| PDB accession code | --- |

Statistics for the highest-resolution shell are shown in parentheses.
<sup>a</sup> $R_{\text{merge}} = \frac{\sum_{hkl} \sum_i |I_i(hkl) - \langle I(hkl) \rangle|}{\sum_{hkl} \sum_i I_i(hkl)}$ .
<sup>b</sup> $R_{\text{meas}} = \frac{\sum_{hkl} [N/(N-1)]^{1/2} \times \sum_i |I_i(hkl) - \langle I(hkl) \rangle|}{\sum_{hkl} \sum_i I_i(hkl)}$ , where $I_i(hkl)$ is the *i*th observation of reflection *hkl*, $\langle I(hkl) \rangle$ is the weighted average intensity for all observations *i* of reflection *hkl* and *N* is the number of observations of reflection *hkl*.
<sup>c</sup> *CC*<sub>1/2</sub> is the correlation coefficient between symmetry equivalent intensities from random halves of the dataset.
<sup>d</sup> The data set was split into "working" and "free" sets consisting of 95 and 5% of the data respectively. The free set was not used for refinement.
<sup>e</sup> The R-factors *R*<sub>work</sub> and *R*<sub>free</sub> are calculated as follows: $R = \frac{\sum (|F_{\text{obs}} - F_{\text{calc}}|)}{\sum |F_{\text{obs}}|}$ , where *F*<sub>obs</sub> and *F*<sub>calc</sub> are the observed and calculated structure factor amplitudes, respectively.
<sup>f</sup> As calculated using MolProbity [32].

**Table S6.** Summary of X-ray data and model parameters for stemmadenine acetate-bound *Ti*DPAS2

|  |  |
| --- | --- |
| Data collection |  |
| Paul Scherrer Institute | 10SA (PX II) |
| Wavelength (Å) | 1 |
| Resolution range (Å) | 39.92 – 2.24 (2.32 – 2.24) |
| Space Group | *P* 21 21 21 |
| Cell parameters (Å) | *a* = 73.186, *b* = 79.845, *c* = 130.922, β = 90° |
| Total no. of measured reflections | 432608 (21387) |
| Unique reflections | 35719 (2561) |
| Multiplicity | 12.1 (8.4) |
| Mean *I*/σ(*I*) | 17.69 (1.77) |
| Completeness (%) | 94.79 (68.96) |
| *R*<sub>merge</sub><sup>a</sup> | 0.1239 (1.273) |
| *R*<sub>meas</sub><sup>b</sup> | 0.1294 (1.358) |
| *CC*<sub>1/2</sub><sup>c</sup> | 0.999 (0.586) |
| Wilson *B* value (Å<sup>2</sup>) | 44.54 |
| Refinement |  |
| Reflections used in refinement | 35691 (2561) |
| Reflections used for R-free | 1786 (128) |
| *R*<sub>work</sub> | 0.1737 (0.3245) |
| *R*<sub>free</sub> | 0.2199 (0.3957) |
| *CC*<sub>work</sub> | 0.972 (0.790) |
| *CC*<sub>free</sub> | 0.957 (0.700) |
| Protein residues | 717 |
| Number of non-hydrogen atoms | 5530 |
| macromolecules | 5272 |
| ligands | 114 |
| solvent | 168 |
| Ramachandran plot: favoured/allowed/disallowed<sup>f</sup> (%) | 96.49/3.51/0.00 |
| Rotamer outliers (%) | 2.79 |
| R.m.s. bond distance deviation (Å) | 0.148 |
| R.m.s. bond angle deviation (°) | 4.02 |
| Clashscore | 5.86 |
| Mean *B* factors: protein/waters/ligands/overall (Å<sup>2</sup>) | 45.79/46.44/45.13/45.78 |
| PDB accession code | --- |

Statistics for the highest-resolution shell are shown in parentheses.
<sup>a</sup> $R_{\text{merge}} = \sum_{hkl} \sum_i |I_i(hkl) - \langle I(hkl) \rangle| / \sum_{hkl} \sum_i I_i(hkl)$ .
<sup>b</sup> $R_{\text{meas}} = \sum_{hkl} [N/(N-1)]^{1/2} \times \sum_i |I_i(hkl) - \langle I(hkl) \rangle| / \sum_{hkl} \sum_i I_i(hkl)$ , where $I_i(hkl)$ is the *i*th observation of reflection *hkl*, $\langle I(hkl) \rangle$ is the weighted average intensity for all observations *i* of reflection *hkl* and *N* is the number of observations of reflection *hkl*.
<sup>c</sup> *CC*<sub>1/2</sub> is the correlation coefficient between symmetry equivalent intensities from random halves of the dataset.
<sup>d</sup> The data set was split into "working" and "free" sets consisting of 95 and 5% of the data respectively. The free set was not used for refinement.
<sup>e</sup> The R-factors *R*<sub>work</sub> and *R*<sub>free</sub> are calculated as follows: $R = \sum (|F_{\text{obs}} - F_{\text{calc}}|) / \sum |F_{\text{obs}}|$ , where *F*<sub>obs</sub> and *F*<sub>calc</sub> are the observed and calculated structure factor amplitudes, respectively.
<sup>f</sup> As calculated using MolProbity [32].

**Table S7.** Summary of X-ray data and model parameters for *Cr*GS

|  |  |
| --- | --- |
| Data collection |  |
| Diamond Light Source beamline | I03 |
| Wavelength (Å) | 0.9762 |
| Detector | Pilatus3 6M |
| Resolution range (Å) | 63.84 – 2.00 (2.05 – 2.0) |
| Space Group | *P*2<sub>1</sub> |
| Cell parameters (Å) | *a* = 55.4, *b* = 101.0, *c* = 66.7, $\beta$ = 106.8° |
| Total no. of measured intensities | 299680 (22122) |
| Unique reflections | 46753 (3406) |
| Multiplicity | 6.4 (6.5) |
| Mean *I*/$\sigma$(*I*) | 9.8 (3.2) |
| Completeness (%) | 98.7 (96.8) |
| *R*<sub>merge</sub><sup>a</sup> | 0.150 (0.740) |
| *R*<sub>meas</sub><sup>b</sup> | 0.180 (0.886) |
| *CC*<sub>1/2</sub><sup>c</sup> | 0.986 (0.648) |
| Wilson *B* value (Å<sup>2</sup>) | 19.2 |
| Refinement |  |
| Resolution range (Å) | 63.84 – 2.00 (2.05 – 2.00) |
| Reflections: working/free<sup>d</sup> | 44395/2334 |
| *R*<sub>work</sub>/*R*<sub>free</sub><sup>e</sup> | 0.205/0.232 (0.401/0.413) |
| Ramachandran plot: favoured/allowed/disallowed<sup>f</sup> (%) | 97.1/2.8/0.1 |
| R.m.s. bond distance deviation (Å) | 0.004 |
| R.m.s. bond angle deviation (°) | 1.65 |
| No. of protein residues: (ranges) | A:351 (9-129, 132-361)<br>B:352 (8-129, 134-363) |
| No. of water/Zinc/NAP molecules | 279/4/2 |
| Mean *B* factors: protein/waters/ligands/overall (Å<sup>2</sup>) | 15/33/24 |
| PDB accession code | 8A3N |

Values in parentheses are for the outer resolution shell.
<sup>a</sup> $R_{\text{merge}} = \sum_{hkl} \sum_i |I_i(hkl) - \langle I(hkl) \rangle| / \sum_{hkl} \sum_i I_i(hkl)$ .
<sup>b</sup> $R_{\text{meas}} = \sum_{hkl} [N/(N-1)]^{1/2} \times \sum_i |I_i(hkl) - \langle I(hkl) \rangle| / \sum_{hkl} \sum_i I_i(hkl)$ , where $I_i(hkl)$ is the *i*th observation of reflection *hkl*, $\langle I(hkl) \rangle$ is the weighted average intensity for all observations *i* of reflection *hkl* and *N* is the number of observations of reflection *hkl*.
<sup>c</sup> *CC*<sub>1/2</sub> is the correlation coefficient between symmetry equivalent intensities from random halves of the dataset.
<sup>d</sup> The data set was split into "working" and "free" sets consisting of 95 and 5% of the data respectively. The free set was not used for refinement.
<sup>e</sup> The R-factors *R*<sub>work</sub> and *R*<sub>free</sub> are calculated as follows: $R = \sum(|F_{\text{obs}} - F_{\text{calc}}|) / \sum |F_{\text{obs}}|$ , where *F*<sub>obs</sub> and *F*<sub>calc</sub> are the observed and calculated structure factor amplitudes, respectively.
<sup>f</sup> As calculated using MolProbity [32].

**Table S8.** Genbank accession for sequences used to construct tree of maximum likelihood in Figure 4A.

| Gene Name | Genbank accession |
| --- | --- |
| *Arabidopsis thaliana* cinnamyl alcohol dehydrogenase 5 (CAD5) | NM_119587.4 |
| *Populus tremuloides* sinapyl alcohol dehydrogenase (SAD) | AF273256.1 |
| *Camptotheca accuminata* 8-hydroxygeraniol oxidase (8HGO) | AY342355.1 |
| *Ocimum basilicum* geraniol dehydrogenase (GEDH) | AY879284.1 |
| *Rauwolfia serpentina* cinnamyl alcohol dehydrogenase (CAD) | KT369739.1 |
| *Catharanthus roseus* 8-hydroxygeraniol dehydrogenase (8HGO) | KF561458.1 |
| *Strychnos speciosa* Wieland-Gumlich aldehyde synthase (WS) | OM304303.1 |
| *Strychnos nux-vomica* Wieland-Gumlich aldehyde synthase (WS) | OM304294.1 |
| *Catharanthus roseus* Geissoschizine synthase (GS) | MF770507.1 |
| *Cinchona pubescens* dihydrocorinantheine aldehyde synthase (DCS) | MW456554 |
| *Catharanthus roseus* tabersonine 3- reductase (T3R) | KP122966.1 |
| *Catharanthus roseus* tetrahydroalstonine synthase (THAS) | KM524258.1 |
| *Catharanthus roseus* heteroyohimbine synthase (HYS) | KU865325.1 |
| *Rauwolfia serpentina* vomilenine reductase 2 (VR2) | KT369740.1 |
| *Rauwolfia tetraphylla* vomilenine reductase 2 (VR2) | KT369741.1 |
| *Tabernanthe iboga* dihydroprecondylocarpine acetate synthase 1 (DPAS1) | MK840855.1 |
| *Tabernanthe iboga* dihydroprecondylocarpine acetate synthase 2 (DPAS2) | MK840856.1 |
| *Catharanthus roseus* dihydroprecondylocarpine acetate synthase 1 (DPAS) | KU865331.1 |

## Notes

### Competing Interest Statement

The authors have declared no competing interest.

## References

[1] H. Jörnvall, J. Hedlund, T. Bergman, Y. Kallberg, E. Cederlund, B. Persson, in Chemico-Biological Interactions, Elsevier, 2013, pp. 91–96.

[2] S. B. Raj, S. Ramaswamy, B. V. Plapp, Biochemistry-us 2014, 53, 5791–5803.

[3] B. V. Plapp, B. R. Savarimuthu, D. J. Ferraro, J. K. Rubach, E. N. Brown, S. Ramaswamy, Biochemistry-us 2017, 56, 3632–3646.

[4] D. S. Auld, T. Bergman, Cell Mol Life Sci 2008, 65, 3961.

[5] L. Caputi, J. Franke, S. C. Farrow, K. Chung, R. M. E. Payne, T.-D. Nguyen, T.-T. T. Dang, I. S. T. Carqueijeiro, K. Koudounas, T. D. de Bernonville, B. Ameyaw, D. M. Jones, I. J. C. Vieira, V. Courdavault, S. E. O’Connor, Science 2018, 360, 1235–1239.

[6] S. C. Farrow, M. O. Kamileen, L. Caputi, K. Bussey, J. E. A. Mundy, R. C. McAtee, C. R. J. Stephenson, S. E. O’Connor, J Am Chem Soc 2019, 141, 12979–12983.

[7] L. Caputi, J. Franke, K. Bussey, S. C. Farrow, I. J. C. Vieira, C. E. M. Stevenson, D. M. Lawson, S. E. O’Connor, Nat Chem Biol 2020, 16, 383–386.

[8] Y. Qu, M. E. A. M. Easson, R. Simionescu, J. Hajicek, A. M. K. Thamm, V. Salim, V. D. Luca, Proc National Acad Sci 2018, 115, 3180–3185.

[9] D. Williams, Y. Qu, R. Simionescu, V. D. Luca, Plant J 2019, 99, 626–636.

[10] A. S. Sandholu, S. P. Mujawar, R. Krithika, H. V. Thulasiram, K. Kulkarni, Proteins Struct Funct Bioinform 2020, 88, prot.25891.

[11] J. Strommer, Plant J 2011, 66, 128–142.

[12] C. Lee, D. L. Bedgar, L. B. Davin, N. G. Lewis, Org Biomol Chem 2013, 11, 1127–1134.

[13] P. J. Baker, K. L. Britton, M. Fisher, J. Esclapez, C. Pire, M. J. Bonete, J. Ferrer, D. W. Rice, Proc National Acad Sci 2009, 106, 779–784.

[14] A. Vitale, N. Thorne, S. Lovell, K. P. Battaile, X. Hu, M. Shen, S. D’Auria, D. S. Auld, Plos One 2013, 8, e63828.

[15] E. C. Tatsis, I. Carqueijeiro, T. D. D. Bernonville, J. Franke, T.-T. T. Dang, A. Oudin, A. Lanoue, F. Lafontaine, A. K. Stavrinides, M. Clastre, V. Courdavault, S. E. O’connor, Nat Commun 2017, 8, 316.

[16] A. Stavrinides, E. C. Tatsis, L. Caputi, E. Foureau, C. E. M. Stevenson, D. M. Lawson, V. Courdavault, S. E. O’Connor, Nat Commun 2016, 7, 12116.

[17] A. Stavrinides, E. C. Tatsis, E. Foureau, L. Caputi, F. Kellner, V. Courdavault, S. E. O’Connor, Chem Biol 2015, 22, 336–41.

[18] M. Geissler, M. Burghard, J. Volk, A. Staniek, H. Warzecha, Planta 2016, 243, 813–824.

[19] P. Stockinger, S. Roth, M. Müller, J. Pleiss, Chembiochem 2020, 21, 2689–2695.

## References

[1] L. Caputi, J. Franke, S. C. Farrow, K. Chung, R. M. E. Payne, T.-D. Nguyen, T.-T. T. Dang, I. S. T. Carqueijeiro, K. Koudounas, T. D. de Bernonville, B. Ameyaw, D. M. Jones, I. J. C. Vieira, V. Courdavault, S. E. O’Connor, Science 2018, 360, 1235–1239.

[2] S. C. Farrow, M. O. Kamileen, L. Caputi, K. Bussey, J. E. A. Mundy, R. C. McAtee, C. R. J. Stephenson, S. E. O’Connor, J Am Chem Soc 2019, 141, 12979–12983.

[3] M. Jarret, V. Turpin, A. Tap, J. Gallard, C. Kouklovsky, E. Poupon, G. Vincent, L. Evanno, Angewandte Chemie Int Ed 2019, 58, 9861–9865.

[4] A. Stavrinides, E. C. Tatsis, E. Foureau, L. Caputi, F. Kellner, V. Courdavault, S. E. O’Connor, Chem Biol 2015, 22, 336–41.

[5] Y. Qu, M. E. A. M. Easson, R. Simionescu, J. Hajicek, A. M. K. Thamm, V. Salim, V. D. Luca, Proc National Acad Sci 2018, 115, 3180–3185.

[6] L. Caputi, J. Franke, K. Bussey, S. C. Farrow, I. J. C. Vieira, C. E. M. Stevenson, D. M. Lawson, S. E. O’Connor, Nat Chem Biol 2020, 16, 383–386.

[7] N. S. Berrow, D. Alderton, S. Sainsbury, J. Nettleship, R. Assenberg, N. Rahman, D. I. Stuart, R. J. Owens, Nucleic Acids Res 2007, 35, e45–e45.

[8] S. S. Jeong, J. E. Gready, Anal Biochem 1994, 221, 273–277.

[9] T. Bruhn, A. Schaumlöffel, Y. Hemberger, G. Bringmann, Chirality 2013, 25, 243–249.

[10] D. Liebschner, P. V. Afonine, M. L. Baker, G. Bunkóczi, V. B. Chen, T. I. Croll, B. Hintze, L.-W. Hung, S. Jain, A. J. McCoy, N. W. Moriarty, R. D. Oeffner, B. K. Poon, M. G. Prisant, R. J. Read, J. S. Richardson, D. C. Richardson, M. D. Sammito, O. V. Sobolev, D. H. Stockwell, T. C. Terwilliger, A. G. Urzhumtsev, L. L. Videau, C. J. Williams, P. D. Adams, Acta Crystallogr Sect D 2019, 75, 861–877.

[11] P. Emsley, B. Lohkamp, W. G. Scott, K. Cowtan, Acta Crystallogr Sect D Biological Crystallogr 2010, 66, 486–501.

[12] W. Kabsch, Acta Crystallogr Sect D Biological Crystallogr 2010, 66, 125–132.

[13] G. Winter, J Appl Crystallogr 2010, 43, 186–190.

[14] P. R. Evans, G. N. Murshudov, Acta Crystallogr Sect D Biological Crystallogr 2013, 69, 1204–1214.

[15] N. Stein, J Appl Crystallogr 2008, 41, 641–643.

[16] A. J. McCoy, R. W. Grosse-Kunstleve, P. D. Adams, M. D. Winn, L. C. Storoni, R. J. Read, J Appl Crystallogr 2007, 40, 658–674.

[17] G. N. Murshudov, P. Skubák, A. A. Lebedev, N. S. Pannu, R. A. Steiner, R. A. Nicholls, M. D. Winn, F. Long, A. A. Vagin, Acta Crystallogr Sect D Biological Crystallogr 2011, 67, 355–367.

[18] K. Cowtan, Acta Crystallogr Sect D Biological Crystallogr 2006, 62, 1002–1011.

[19] O. Trott, A. J. Olson, J Comput Chem 2010, 31, 455–461.

[20] Y. Kochnev, E. Hellemann, K. C. Cassidy, J. D. Durrant, Bioinformatics 2020, 36, btaa579-.

[21] W. Tian, C. Chen, X. Lei, J. Zhao, J. Liang, Nucleic Acids Res 2018, 46, gky473-.

[22] R. C. Edgar, Biorxiv 2021, 2021.06.20.449169.

[23] J. Trifinopoulos, L.-T. Nguyen, A. von Haeseler, B. Q. Minh, Nucleic Acids Res 2016, 44, W232–W235.

[24] I. Letunic, P. Bork, Nucleic Acids Res 2021, 49, gkab301-.

[25] S. Zhao, R. B. Andrade, J Org Chem 2017, 82, 521–531.

[26] M. E. Kuehne, U. K. Bandarage, A. Hammach, Y.-L. Li, T. Wang, J Org Chem 1998, 63, 2172–2183.

[27] B. Hong, D. Grzech, L. Caputi, P. Sonawane, C. E. R. López, M. O. Kamileen, N. J. H. Lozada, V. Grabe, S. E. O’Connor, Nature 2022, 1–6.

[28] F. Trenti, K. Yamamoto, B. Hong, C. Paetz, Y. Nakamura, S. E. O’Connor, Org Lett 2021, 23, 1793–1797.

[29] Y. Qu, M. L. A. E. Easson, J. Froese, R. Simionescu, T. Hudlicky, V. DeLuca, Proc National Acad Sci 2015, 112, 6224–6229.

[30] A. Stavrinides, E. C. Tatsis, L. Caputi, E. Foureau, C. E. M. Stevenson, D. M. Lawson, V. Courdavault, S. E. O’Connor, Nat Commun 2016, 7, 12116.

[31] M. Geissler, M. Burghard, J. Volk, A. Staniek, H. Warzecha, Planta 2016, 243, 813–824.

[32] I. W. Davis, A. Leaver-Fay, V. B. Chen, J. N. Block, G. J. Kapral, X. Wang, L. W. Murray, W. B. Arendall, J. Snoeyink, J. S. Richardson, D. C. Richardson, Nucleic Acids Res 2007, 35, W375–W383.

